# Coupled effects of salinity and host phylogeny on niche breadth and viral evolution from seawater to salt saturation

**DOI:** 10.64898/2026.06.18.733169

**Authors:** Jaime Alcorta, María Dolores Ramos-Barbero, Fernando Santos, Josefa Antón

## Abstract

Virus-host interactions are fundamental drivers of microbial community structure, yet whether viral ecological niches are confined within individual host niches (nested host niche scenario) or span multiple hosts and exceed any single host niche (expanded host niche scenario) remains poorly understood. To explore these patterns, we characterized prokaryotic and viral distributions and predicted virus-host interactions along a salinity gradient at Bras del Port salterns (Spain), ranging from seawater (3.6% salinity) to salt saturation (39.0%). We analyzed metagenomes and viromes from six ponds supplemented by 27 additional published viromes from the same hypersaline system, recovering 170 metagenome-assembled genomes (MAGs) dereplicated at the genomospecies level (MAGs clustered at 95 % average nucleotide identity), approximately 55,000 viral operational taxonomic units (vOTUs), and nearly 4,000 predicted virus-host pairs. Viruses exhibited broader niches than their putative hosts at the highest salinities, while at lower salinities the pattern was reversed or inconsistent depending on the site, and niche breadths of both viruses and hosts increased steadily toward higher salinities. Host taxonomy at the class level and below was the primary driver of viral genomic clustering, explaining more variance than salinity provenance (approximately 30% *vs.* approximately 19%), while the contribution of salinity to viral genomic composition appeared indirect, mediated through the salinity-driven distribution of distinct host classes rather than direct environmental filtering of viral sequences. Together, these findings support the expanded host niche scenario as the predominant virus-host interaction strategy, with evolutionary and ecological dynamics jointly shaped by salinity and host identity.

## Introduction

Microorganisms occupy distinct ecological niches defined by the environmental conditions where they can survive and compete successfully [1–2]. Their metabolic capabilities and genomic adaptations determine which resources can be exploited within these environments. Metagenomics has revolutionized our ability to simultaneously characterize whether microorganisms are distributed across diverse ecosystems and what metabolic functions they perform, therefore determining the breadth of their ecological niches [1]. Consequently, these patterns of niche occupancy and environmental preferences have been revealed for microbial communities across numerous ecosystems [3–5].

Biotic interactions, and virus-host relationships in particular, play crucial roles in shaping the realized niches of microorganisms [2], yet the association between viral and host niche breadths remains complex and poorly understood. This relationship may be essentially a consequence of the host-range dynamics [6]: viruses infecting a single host lineage would be confined within its ecological niche (the “nested host niche” scenario), whereas broad host-range viruses may occupy the union of multiple host niches, potentially exceeding the environmental range of any individual host (the “expanded host niche” scenario). However, host-range alone does not fully determine viral niche breadth, as viruses may additionally require direct physiological adaptation to abiotic conditions, evidenced by environment-specific signatures in viral proteome composition (e.g., as those reported for polar communities [7], saline lakes [8], or desert viromes [9]). Moreover, the realized niche of a virus is not static, as viruses continuously modify the physiology, population structure, and competitive dynamics of their hosts through infection, reshaping the very ecological space in which subsequent viral generations propagate [10].

Whether the phylogenetic relationships among viruses are driven primarily by the evolutionary history of their hosts or by environmental filtering remains an open question. Network analyses reveal that viral communities are significantly modular and that viral sequences tend to cluster by host phylum, suggesting that host evolutionary history is a dominant structuring force at broad taxonomic scales [11]. Consistent with the nested host niche scenario, most known viral species infect only one or two host taxa [12], and the host-range constraints that underlie this pattern are highly condition-dependent and variable across environmental contexts [6]. On the other hand, both the direct genomic adaptation of viruses to abiotic conditions and the indirect reshaping of host communities through infection suggest that environmental selection leaves an imprint on viral genomic diversity beyond what host phylogeny alone predicts. This is evident at the assemblage level: marine viromes carry a detectable “marine-ness” signature [13], while viromes from near-saturation hypersaline environments display an analogous “hypersaline-ness” quality [14], suggesting that the environment itself may shape viral community composition regardless of host phylogeny.

Natural environmental gradients are ideal systems for studying microbial niche breadth (e.g., [15]), and salinity represents one of the most important environmental factors shaping microbial community structure and diversity in both aquatic and terrestrial ecosystems [16–19]. Hypersaline environments, particularly multi-pond solar salterns, exhibit pronounced shifts in microbial community composition, functional profiles, and viral assemblages along salinity gradients [20–23]. These environments typically show a transition from bacterial-dominated communities at lower salinities to archaeal-dominated communities at higher salinities, with considerable proportions of halophilic *Bacteroidota* persisting across the gradient [21,24]. Viruses are particularly abundant in these systems, with distinct patterns of host range and high microdiversity emerging across salinity gradients [23,25–26]. Solar salterns therefore offer a unique opportunity to simultaneously address viral and prokaryotic niche breadths and disentangle whether viral genomic relationships are structured primarily by salinity or by host taxonomy, given that they harbor phylogenetically diverse microbial groups with broad salinity tolerances co-occurring across a well-defined and connected environmental gradient.

To address these knowledge gaps, we characterized prokaryotic and viral communities and their predicted interactions along a comprehensive salinity gradient at Bras del Port salterns (Santa Pola, Alicante, Spain), spanning from seawater (3.6% salinity) to salt saturation (39.0%) across six ponds (Figure 1A). Using paired metagenomes and viromes, supplemented by 27 additional published viromes from the same system [26], we recovered vOTUs and dereplicated MAGs (genomospecies) to identify putative virus-host pairs and determine their ecological niches and genomic compositional relatedness across the gradient. This allowed us to address two fundamental questions in viral ecology: (i) how salinity shapes viral *versus* host niche breadths, and (ii) whether viral genomic relationships are structured primarily by the evolutionary history of their hosts or by environmental filtering along the salinity gradient. Our findings support an “expanded host niche” scenario as the predominant virus-host interaction strategy across the gradient with contributions of the “nested host niche” scenario particularly at lower salinities. These results also show that niche breadth of both viruses and hosts increases toward hypersaline conditions, and reveal that host evolutionary history leaves a stronger imprint on viral genomic composition than the salinity environment itself.

**Figure 1.**
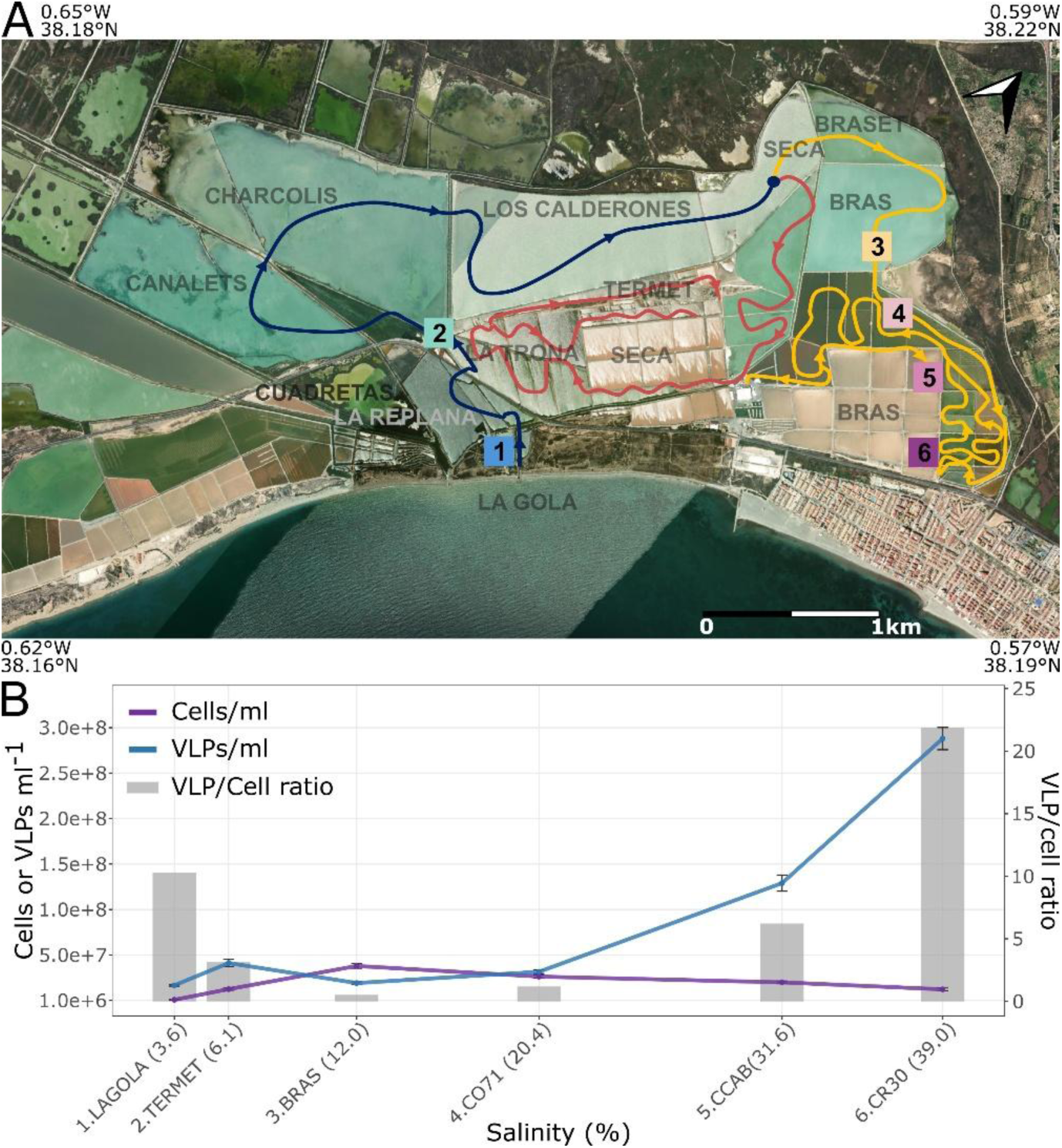
Sampling sites and microbial abundance along the salinity gradient at the Bras del Port solar salterns. (A) Aerial view of the Bras del Port solar salterns (Santa Pola, Alicante, Spain). Satellite imagery © Esri obtained via the maptiles R package. Coloured outlines delineate the flow circuits of the salterns. Numbered squares indicate the six sampling sites with increasing salinity: LAGOLA (1, 3.6%), TERMET (2, 6.1%), BRAS (3, 12.0%), CO71 (4, 20.4%), CCAB (5, 31.6%), and CR30 (6, 39.0%). Scale bar, 1 km. (B) Abundance of virus-like particles (VLPs; blue line) and prokaryotic cells (purple line) per milliliter, and VLP-to-cell ratio (grey bars) across the salinity gradient. VLP counts were determined by SYBR Gold epifluorescence microscopy and cell counts by DAPI staining. Error bars represent standard deviation of triplicate counts.

## Materials and methods

### Sample collection

Samples were collected in September 2016 from six ponds spanning a salinity gradient of 3.6-39.0% at the Bras del Port (BdP) solar salterns in Santa Pola, Alicante, Spain (38.15555° N,-0.7161° W) (Table S1). The sampled ponds comprised two low-salinity ponds (Lagola and Termet), one medium-salinity brine pond (BRAS), two concentrator ponds (CO71 and CCAB), and one crystallizer pond (CR30), from each of which one liter of brine was collected. Salinity was measured in situ with a hand refractometer (Sper Scientific, Scottsdale, AZ, USA), and ionic composition was determined by atomic absorption spectrometry at the Technical Services of the University of Alicante.

### Microbial abundance (viral particles and cells) determined by DAPI, FISH, and SYBR Gold

Viral abundance was determined using SYBR Gold staining as previously described by Boujelben et al. [20]. Briefly, 20 μl of each sample fixed with formaldehyde (4% final concentration, v/v) was filtered through 0.02 μm Anodisc 25 Whatman filters (Whatman, Merck KGaA, Darmstadt, Germany). Viral particles were enumerated using a Leica DM4000B epifluorescence microscope equipped with a 100X oil immersion fluorescence objective (Leica Microsystems, Wetzlar, Germany). Total cells were enumerated by DAPI staining, and archaeal and bacterial populations were identified by fluorescence in situ hybridization (FISH). Samples were fixed with formaldehyde (Sigma-Aldrich, St. Louis, MO, USA) at a final concentration of 6.7% (v/v) for 16 h at 4 °C. Fixation was stopped by diluting 10 times with 1× phosphate-buffered saline (PBS), and the samples were filtered through 0.22 μm pore-size, 16 mm diameter filters (Isopore GTTP; Merck Millipore, Darmstadt, Germany). FISH was performed using the EUB338 and ARC915 probes followed by staining with 4′,6-diamidino-2-phenylindole (DAPI; 1 μg ml⁻¹) as described by Antón et al. [27].

### Cell and viral DNA extraction and sequencing

One liter of brine samples was processed to separate cell and viral fractions. For cell collection, samples were centrifuged at 30,000 × g for 40 min at 20 °C (Beckman Coulter, Brea, CA, USA; JI40 rotor); the resulting pellet was resuspended in approximately 10 ml, re-centrifuged at maximum speed for at least 15 min, and stored at-80 °C until DNA extraction with the UltraClean® Soil DNA Isolation Kit (Mo Bio Laboratories, Carlsbad, CA, USA). The supernatant from the first centrifugation was filtered through 0.22 µm to remove residual cells, and the filtrate was used for viral fraction purification. The viral fraction was concentrated using a Vivaflow 200 PES system (Sartorius, Göttingen, Germany), reducing the initial volume to 20 ml with a 30 kDa cutoff, followed by ultracentrifugation at 286,000 × g for 4 h at 20 °C and resuspension of the pellet in 500 μl of the supernantant (Beckman Coulter, Brea, CA, USA; SW41T rotor). An aliquot of 50 μl from the concentrated sample was used to determine viral genome sizes by pulsed-field gel electrophoresis (PFGE).

Viral DNA extraction was performed as previously described [14]. Briefly, virus concentrates were mixed with equal volumes of 1.6% low-melting agarose (Pronadisa, Alcobendas, Madrid, Spain) and dispensed into 100 ml molds to solidify at room temperature, yielding agarose plugs that physically embed and protect the viral particles. These plugs were then incubated with Turbo DNase (Ambion, Thermo Fisher Scientific, Waltham, MA, USA) prior to ESP treatment to minimize contamination from cellular DNA [14,23]. The absence of cell DNA contamination in viral fractions was verified by PCR using archaeal 16S rRNA gene primers 341F and 907R [21]. No cellular contamination was detected in any viral DNA sample. Then, quality and quantity of cell and viral DNA were assessed using a Qubit® 2.0 fluorometer (Thermo Fisher Scientific, Waltham, MA, USA) with the high-sensitivity assay kit (2-100 ng). Sequencing of both cell and viral DNA was carried out on the Illumina MiSeq platform (Illumina, San Diego, CA, USA) using Nextera XT libraries with 2 × 300 bp paired-end reads.

Fastq files were initially processed with the TrimGalore v0.6.6 software, which uses Cutadapt v2.8 [28], with paired-end mode and the following options:-q 28 --length 50 --max_n 1 --trim-n and --illumina. Then, the sequencing effort was assessed with the Nonpareil v3.401 software [29–30] with the-T alignment and-f fasta options. Viral sequence enrichment of the samples was estimated with the ViromeQC v1.0.2 software [31] with the-w environmental parameter.

### Metagenomic assembly with contig and CDS-level profiling

*De novo* assemblies of trimmed reads were generated with MEGAHIT v1.1.3 [32] using default parameters and the --presets meta-large option, and assemblies were evaluated with QUAST v5.0.2 [33]. Coding sequences (CDS) were predicted with Prodigal v2.6.3 [34] using the-p meta option. Contigs were then classified hierarchically: viral and plasmid contigs were identified with geNomad v1.8.1 (end-to-end option [35]), unclassified contigs ≥1 kb were assigned to prokaryotes or eukaryotes with Whokaryote [36], and the remaining contigs were left unclassified. Taxonomic affiliations of prokaryotic, unclassified and plasmid contigs and their CDS were obtained with RefineM v0.4.0 [37], using the scaffold_stats and taxon_profile modules with these predicted CDS and a custom DIAMOND v2.1.9.163 database [38] built from CDSs of representative genomes of the GTDB R220 and formatted with a custom Python script.

Functional annotation of predicted CDS was performed using three complementary tools: emapper v2.1.12 [39–40], run with the-m diamond option against eggNOG DB v5.0.2; DRAM v1.5.0 [41], using the annotate_genes and distill modules with results retrieved from the KOfam [42] and PFAM [43] databases; and METABOLIC-G.pl v4.0 [44]. To determine abundances, trimmed reads were mapped against contig and predicted CDS nucleotide sequences using CoverM v0.6.1 [45] with the following specific parameters: --min-read-percent-identity 95 --min-read-aligned-percent 50 --min-covered-fraction 0. Functional, taxonomic, and abundance information was then integrated for each contig and CDS using custom bash and Python scripts.

### Metagenome assembled genomes

The metaWRAP binning module [46] was used to group contigs with the following software: MetaBAT2 v2.12.1 [47], MaxBin2 [48] and CONCOCT v1.1.0 [49], with default parameters. Next, the bin_refinement module of metaWRAP [46] was used to obtain merged bins with the options-c 50 (completeness threshold) and-x 10 (contamination threshold). Potential contamination of contigs with different genomic properties was cleaned with the RefineM v0.4.0 tool [37], through the modules scaffold_stats, outliers and filter_bins, and by different taxonomic assignment with the module call_genes, taxon_profile and taxon_filter with the GTDB R95 custom protein database, and ssu_erroneous, with the GTDB R80 custom SSU database.

Taxonomic assignment of MAGs was performed with GTDB-Tk v2.4.0 using database version R220 [50]. Genome quality was assessed with CheckM v1.0.12 [51] and CheckM2 v1.0.2 [52]. Non-coding RNA elements and other genomic features, including tRNAs and ribosomal RNA subunits, were annotated with Bakta v1.9.4 [53]. Following quality filtering, MAGs were dereplicated using the dereplicate function of dRep v3.5.0 [54], selecting one representative genome per species cluster at 95% ANI (i.e., genomospecies; parameters-comp 50-con 10 --S_algorithm skani-pa 0.75-sa 0.95-nc 0.75). The distribution of dereplicated MAGs across the salinity gradient was then determined by mapping reads back to all samples with CoverM v0.6.1 [45] with above parameters. To account for uneven read mapping across genomes, presence-absence was additionally evaluated with metapresence v1.0 [55] under both default and relaxed settings (-ber_threshold 0.7 - fug_threshold 0.45); this tool internally uses Bowtie2 v2.5.4 [56] and SAMtools v1.21 [57].

Phylogenomic reconstructions were performed from a total of 120 and 53 single-copy concatenated genes for bacteria and archaea, respectively, which were recovered from the intermediate files of the GTDB-tk v2.4.0 analysis. Maximum-likelihood trees were generated using IQtree v.1.5.5 software [58], with the TESTNEW option to choose the best substitution models [59], after which a nonparametric ultrafast bootstrap (-bb) support of 1000 replicates was applied [60]. Node collapse and rooting of phylogenetic reconstructions were managed using the iTOL web server [61].

### Determination of viral operational taxonomic units (vOTUs)

To cluster the viral contigs into vOTUs, the MVP v1.1.4 software [62] was implemented. This tool uses at the first step (MVP_02_filter_genomad_checkv) geNomad v1.7.4 [35] and checkV v1.0.3, with the 1.5 version of the database [63]. Then, viral contigs were clustered (MVP_03_do_clustering) with default options (--min_ani 95 and --min_tcov 85). From the MVP pipeline, two sets of vOTUs were obtained and used for downstream analyses: (i) one called the “relaxed MVP representatives” (related to the MVP_02_filter_genomad_checkv step), and (ii) the “conservative MVP representatives”, which are obtained in the step MVP_05_create_vOTU_table and which take into account viral sequences longer than 5 kb or longer than 1 kb when qualified as complete, high-, or medium-quality by checkV [62].

To further investigate potential vOTUs not detected by the MVP pipeline (which is reliant mostly on geNomad), all the contigs were mapped with coverM v0.6.1 [45] (specific parameters: --min-covered-fraction 0 --min-read-percent-identity 95 --min-read-aligned-percent 50) against a set of 27 Bras del Port viromes previously obtained [26] and available under the NCBI Bioproject PRJEB69609. Considering that the 27 samples were from five sites with salinities >20 % (ponds CM1, CM2, CCAB, CR30 and CR41), and that the maximum number of samples per site was 7, those initial contigs that were mapped with over 10% of horizontal coverage in ≥7 samples (therefore potentially present either in all the samples from one site or in two or more sites) were used for further analyses. Therefore, a total of 139,216 mapped contigs were collapsed in 76,915 vOTUS with the ani_cluster utility of the MGV package (https://github.com/snayfach/MGV/ [64]), clustering at 95% of identity and 85% of the aligned fraction based on a blastn v2.15.0+ reciprocal search [65] between all selected viral contigs to choose the representative mapped vOTUs.

Thus, two sets of vOTUs were obtained combining the analyses from MVP and the previously available viromes: the “relaxed” set of vOTUs (55,691 vOTUs), which comprised the relaxed MVP vOTUs with sizes ≥1 kb, plus the vOTUs ≥1 kb obtained by the virome mapping strategy, and the “conservative” set (8,139 vOTUs) which comprised the conservative MVP OTUs plus the vOTUS ≥5 kb of the virome mapping strategy. Both sets were quantified through the same read mapping procedures used for the dereplicated MAGs: coverM (mapping with bwa-mem at 95 % identity) and filtering by aligned fraction (>0, >10, >50, and >80% aligned fraction) and with both sets of parameters using metapresence. The threshold to determine the presence of a vOTU in a sample was set to ≥50% aligned fraction as determined in Supplementary Text. Differential abundance analysis of vOTUs was performed by comparing virome and metagenome samples using log_2_ fold-change calculations on transcript per million abundances (TPM), with statistical significance assessed through the DESeq2 R package [66] with Benjamini-Hochberg correction (adjusted p-value < 0.05).

The relationships among the vOTUs, in terms of their shared protein clusters, were explored through a network approach. CDS were predicted with the pharokka v1.7.5 software [67], with the --meta mode which uses Pyrodigal_gv v0.2.0 [68] and then clustered with vConTACT v2 [69–70], with parameters: --rel-mode’Diamond’ --db’ProkaryoticViralRefSeq211-Merged’ --pcs-mode MCL --vcs-mode ClusterONE.

### Virus-host relationships inference

For prediction of virus-host relationships, the association between vOTUs and dereplicated MAGs was explored using two different tools. Since metagenomic binning precedes viral sequence identification, contigs subsequently classified as viral may have been inadvertently incorporated into MAGs. To avoid spurious virus-host associations derived from this pipeline artifact, vOTUs sharing ≥99% nucleotide identity and ≥10% query coverage (blastn v2.15.0+ [65]) with contigs binned into MAGs were excluded prior to prediction analyses. Then, software PHIST v1.2.1 [71] was used with default parameters, and software WiSH v1.1 [72] with the build and predict tools with no null-parameters provided. Virus-host pairs were analyzed in three subsets for both the relaxed and the conservative vOTUs: a) those associations detected by PHIST, b) those detected by PHIST and with a minimum of 3 shared k-mers, and c) those detected by PHIST and WiSH. The presence breadth of virus-host pairs across these subsets was determined from presence/absence patterns using the methods described above. Validation of virus-host relationship prediction tools with experimentally confirmed associations for genera present in the MAGs is available in Supplementary Material.

Niche position (abundance-weighted mean salinity) and niche breadth (abundance-weighted standard deviation of salinity around the niche position), modified from the ecospat.nichePOSNB min-max approach by using a sum-col strategy as detailed in Supplementary Text, were calculated for each vOTU and MAG using a custom R script (available at https://github.com/jaalcort/col-sum_nichePosBdt_calculator) following the conceptual framework of the ecospat package [73], including only samples where detection thresholds were met (aligned fraction >50% and metapresence default for vOTUs and MAGs, respectively). For MAGs, calculations were performed directly across the six metagenomic samples. For vOTUs, to avoid bias from differences in sequencing depth and viral capture efficiency between library types, niche position was estimated independently for metagenomic and viromic libraries and then combined as a detection-rate weighted average (weighted by the fraction of sites with TPM > 0, requiring a minimum of two sites per library type); niche breadth was recalculated as the square root of the TPM-weighted variance around this consensus position, pooling detections from both library types. Full equations and comparisons with the standard ecospat and Gaussian fitting approaches are provided in Supplementary Text.

### Determining influence of biotic and abiotic factors in genomic relatedness

To determine if there was an effect of the salinity over the genomic ANI between all related viral contigs (the pre-clustering contigs used to obtain the relaxed vOTUs), they were processed with the ani_cluster utility of the MGV package [64], and ANI values were filtered for qcov >0, tcov >0/85 and ANI >70 %. Then, a difference of the salinity value was calculated for each pair of viral contigs regarding their origin samples. The “salinity decay” (i.e., influence of differences of salinity on the genetic distance) was investigated via simple linear regression analysis between the ANI and the absolute difference of salinity for each pair of viral contigs with the lm() R function. These results were contrasted with the ANI differences of non-dereplicated MAGs obtained from the intermediary files of the drep analysis.

To evaluate the influence of environmental and biological factors on viral genomic composition as a proxy for evolutionary relationships among vOTUs, tetranucleotide frequencies were calculated for all relaxed vOTUs using the scaffold_stats utility from RefineM v0.4.0 [37], producing 136-dimensional vectors subsequently normalized to relative frequencies. Three distance metrics were computed to capture compositional similarity: Bray-Curtis dissimilarity, as the primary metric for compositional data [74]; Euclidean distance, to assess geometric relationships in multivariate space [75]; and cosine distance (1 - cosine similarity), to focus on compositional patterns independent of vector magnitude [76]. All metrics were calculated using the vegan package v2.6-4 in R. Principal Coordinate Analysis (PCoA) was then performed on the Bray-Curtis dissimilarity matrix using the ape package v5.7-1 [77] to visualize major compositional patterns across vOTUs. Then, the contribution of categorical and continuous variables to tetranucleotide composition variation was assessed through two complementary approaches. For categorical variables tested individually, permutational multivariate analysis of variance (PERMANOVA) was conducted using the adonis2 function in the vegan package [78] with 999 permutations. For continuous variables, Mantel tests [79] were performed using the vegan package with 999 permutations.

## Results and Discussion

### Overall taxonomic and functional characterization of samples

Virus-like particles (VLPs) ranged from ∼2 to ∼30 × 10⁷ per milliliter, with virus-to-cell ratios of ∼10 in the marine pond (LAGOLA) and ∼22 in saturated brines (CR30) (Table S1), consistent with previous reports [24,80]. The virus-to-cell ratio exhibited a U-shaped pattern along the salinity gradient, with a pronounced minimum at intermediate salinities (BRAS; ratio = 0.52), where cell abundance peaked (∼4 × 10⁷ cells ml⁻¹) but VLP concentrations remained relatively low (∼2 × 10⁷ VLPs ml⁻¹) (Figure 1B, Table S1). This may reflect a bacterial bloom of fast-growing, moderately halophilic bacteria in the transition zone at the time of sampling, where viral predation may not yet fully equilibrate with the host population growth [81–82].

Sequencing depth and assembly quality metrics are summarized in Table 1. Nonpareil coverage analysis demonstrated that sequencing depth was adequate for reliable community comparisons across all samples, with saturation curves reaching plateaus at lower sequencing efforts in higher salinity samples (Figure S1). Coverage values increased with salinity while diversity indices decreased (Figure S1), consistent with the lower prokaryotic diversity previously observed at the top of the salinity gradient [83]. ViromeQC analysis confirmed successful viral enrichment across all viromes, with particularly high enrichment scores (>35, [31]) in the viromes of intermediate-to-high salinity ponds CO71 and CCAB (Table 1). Assembly quality was robust across both metagenomes and viromes, with most reads mapping back to assembled contigs (Table 1).

**Table 1.** Sequence features of metagenomic and virome samples. Sequencing metrics and assembly statistics for metagenomes and viromes across the salinity gradient. Summary of quality control, sequencing coverage, and assembly metrics for six metagenomic and six viromic samples collected along the salinity gradient at Bras del Port salterns (Santa Pola, Spain). Total trimmed reads represent the number of sequences after quality filtering. Non-pareil C indicates sequencing coverage (proportion of community sampled), while non-pareil diversity represents the estimated microbial diversity index. ViromeQC total enrichment score reflects the quality of viral enrichment in metaviromic samples. Assembly statistics include: number of contigs (≥0 bp), total assembled sequence length (≥0 bp), length of the largest contig, GC content (%), and N50 (contig length at 50% of total assembly). Final columns show the percentage of total trimmed reads successfully mapped to the complete assembly and to contigs >1,000 bp.

| Sample | Salinity (%) | Total trimmed reads | non-pareil C | non-pareil diversity | viromeQC total enrichment score | n contigs ($\geq 0$ bp) | Total length ($\geq 0$ bp) | Largest contig | GC (%) | N50 | Percentage of total reads mapped to assembly (all contigs) | Percentage of total reads mapped to assembly (contigs $> 1000$bp) |
| --- | --- | --- | --- | --- | --- | --- | --- | --- | --- | --- | --- | --- |
| MG LAGOLA | 3.6 | 99363846.0 | 0.86 | 18.53 | 0.58 | 549856 | 464849715 | 324572 | 47.05 | 1881 | 81.13 | 60.62 |
| MV LAGOLA | 3.6 | 12129382.0 | 0.78 | 17.47 | 8.48 | 80277 | 70029242 | 156905 | 44.29 | 1759 | 73.49 | 58.24 |
| MG TERMET | 6.1 | 254485438.0 | 0.84 | 18.98 | 0.23 | 984838 | 660896290 | 447586 | 49.24 | 1274 | 76.29 | 50.66 |
| MV TERMET | 6.1 | 60006848.0 | 0.91 | 18.35 | 5.36 | 491667 | 413258686 | 681525 | 47.50 | 1574 | 84.81 | 62.96 |
| MG BRAS | 12.0 | 105897316.0 | 0.88 | 18.35 | 0.47 | 399434 | 340388252 | 328700 | 48.74 | 2080 | 78.29 | 64.37 |
| MV BRAS | 12.0 | 24652338.0 | 0.94 | 16.84 | 1.63 | 141370 | 116382101 | 1020022 | 47.98 | 1630 | 86.88 | 63.26 |
| MG CO71 | 20.4 | 72915146.0 | 0.93 | 17.43 | 0.90 | 159597 | 186689278 | 351642 | 56.20 | 3462 | 90.01 | 79.46 |
| MV CO71 | 20.4 | 11201730.0 | 0.97 | 14.96 | 79.42 | 64363 | 38180052 | 103119 | 51.14 | 1120 | 93.16 | 36.66 |
| MG CCAB | 31.6 | 57818056.0 | 0.85 | 18.40 | 0.64 | 198755 | 213406409 | 240446 | 59.20 | 2816 | 80.84 | 72.27 |
| MV CCAB | 31.6 | 4907872.0 | 0.96 | 14.55 | 35.87 | 21865 | 13701459 | 37203 | 56.52 | 1008 | 76.92 | 37.67 |
| MG CR30 | 39.0 | 57477810.0 | 0.96 | 16.63 | 0.95 | 107265 | 116424831 | 274994 | 57.31 | 2900 | 93.51 | 81.40 |
| MV CR30 | 39.0 | 3874368.0 | 0.97 | 15.14 | 1.82 | 20390 | 12759709 | 23377 | 57.46 | 960 | 73.98 | 40.23 |

The taxonomic classification of the assembled contigs revealed distinct patterns between metagenomes and viromes, with viromes containing a substantially higher proportion of viral and unclassified contigs (∼20 and ∼40 %, respectively; Figure 2), reflecting the greater taxonomic novelty characteristic of viral sequences, particularly at higher salinities. Prokaryotic communities were dominated by *Pseudomonadota*, *Bacteroidota*, and *Actinomycetota* among bacteria, and by *Halobacteriota*, particularly *Haloarculaceae* and *Haloferacaceae* among archaea, consistent with previous results [21] (Figure S2; Table S2). Across the salinity gradient, bacterial relative abundance in cellular metagenomes decreased from 72.8% (LAGOLA) to 2.0% (CR30), while archaeal values increased from 0.5% to 79.6%, driven by the decline of *Pseudomonadota* and the rise of *Haloferacaceae* towards higher salinities. Dominant bacterial genera varied across samples: genus HIMB11, closely related to the marine *Roseobacter* clade [84], dominated in LAGOLA; genus HIMB30, within the marine Gammaproteobacteria family *Litorivicinaceae* [85] in TERMET; and the moderately halophilic bacterium *Spiribacter* [86] and the extremely halophilic genus *Salinibacter* [87] dominated in intermediate and high salinity samples, respectively (Figure S2). Archaeal communities showed a clear succession pattern from the BRAS pond onwards. At intermediate-high salinity, *Halorubrum* [88] and *Hallobellus* [89] co-dominated alongside emerging populations of *Haloquadratum* [90]. At the highest salinity, *Haloquadratum* dominated (>37% of reads), with *Halobaculum* [91] as a secondary player. This pattern reflects a progressive ecological filtering along the salinity gradient, where increasing ionic stress selects for extremely halophilic archaeal taxa.

**Figure 2.**
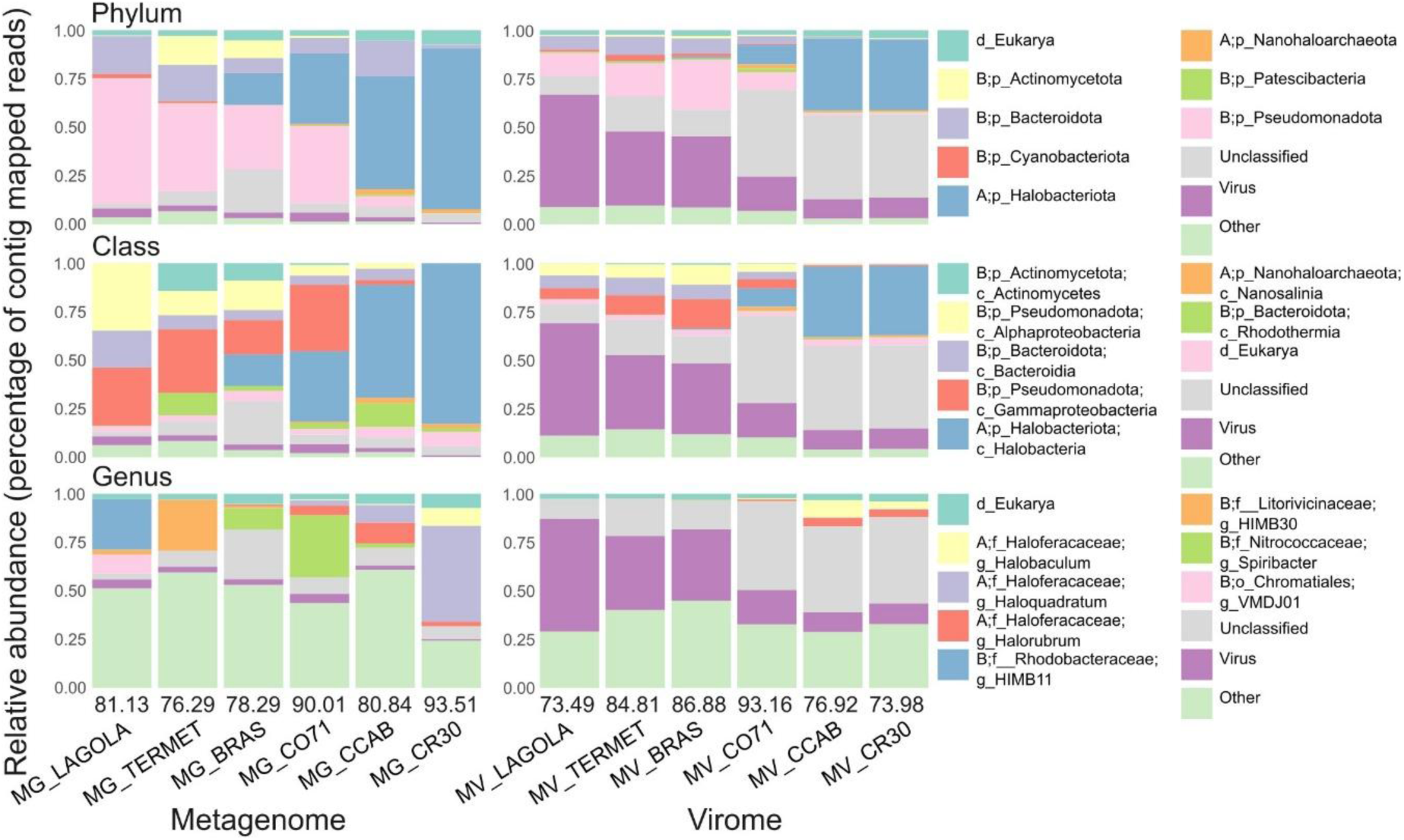
Taxonomic composition of assembled contigs across metagenomes and viromes along the salinity gradient. Stacked bar charts show the relative abundances of contigs at phylum (top), class (middle), and genus (bottom) levels, expressed as the percentage of total contig-mapped reads per sample. Read mapping was performed with CoverM (≥95% nucleotide identity, ≥50% read aligned fraction). Contigs were hierarchically classified as follows: viral and plasmid contigs were identified with geNomad; remaining contigs ≥1 kb were assigned to prokaryotes or eukaryotes with Whokaryote; and prokaryotic, unclassified and plasmid contigs were taxonomically annotated using RefineM against GTDB R220 representative genomes based on predicted ORF consensus. Prefixes indicate domain of origin: A, Archaea; B, Bacteria. Numbers above sample labels indicate the percentage of total trimmed reads successfully mapped to assembled contigs. The ten most abundant taxa are shown per panel; remaining taxa are grouped as “Other.” MG, metagenome; MV, virome.

Functional annotations of predicted CDS revealed consistent COG category patterns across metagenomes, contrasting with higher variability in viromes (Figure S3; Table S3). Both dataset types were dominated by replication, recombination and repair functions (category L), followed by amino acid metabolism in metagenomes (category E) and cell membrane biogenesis in viromes (category M, in which the most abundant functions were K01449 and PF07486, corresponding to cell wall hydrolases in viral and unclassified contigs, among others), reflecting the differential functions of their major entities. Also, the high prevalence of unknown functions in viromes (category S) reflects the unique characteristics of viral proteins [92]. More specific functional classifications revealed transposases as the most abundant KEGG category in metagenomes, while viromes were enriched in phage thymidylate synthases, involved in DNA replication and thymidine hypermodification for host defense evasion [93], and terminases, responsible for DNA packaging into viral capsids. Notably, across the salinity gradient, major biogeochemical pathways exhibited functional redundancy, with the same metabolic functions performed by taxonomically distinct communities (Figure S4), which, at this level of resolution, is an emergent property observed in diverse ecosystems [2].

### High prokaryotic genomic novelty and salinity-dependent distribution patterns

A total of 217 MAGs were obtained (205 from metagenomes and 12 from viromes, mostly from BRAS and TERMET) and dereplicated to 170 genomospecies (31 from Archaea and 139 from Bacteria; Figure 3, Table S4) with an average completeness of 80.75% ± 14.13% and contamination of 2.11% ± 1.82%. Of these, 39 qualified as high-quality and 131 as medium-quality according to Genomic Standards Consortium criteria [94] (Table S4). The taxonomic novelty, according to the GTDB R220 version of the database, comprised one possible new family within *Flavobacteriales* (*Bacteroidota*), 10 possible new genera, and 114 possible new species (accounting for 67.06% of MAGs; Figure S5), underscoring the high degree of unexplored prokaryotic diversity in hypersaline environments [95]. Most potential new genera belonged to *Pseudomonadota*, *Bacteroidota*, *Nanoarchaeota*, and *Dependentiae*, with most phyla containing >50% MAGs classified as potential new species.

**Figure 3.**
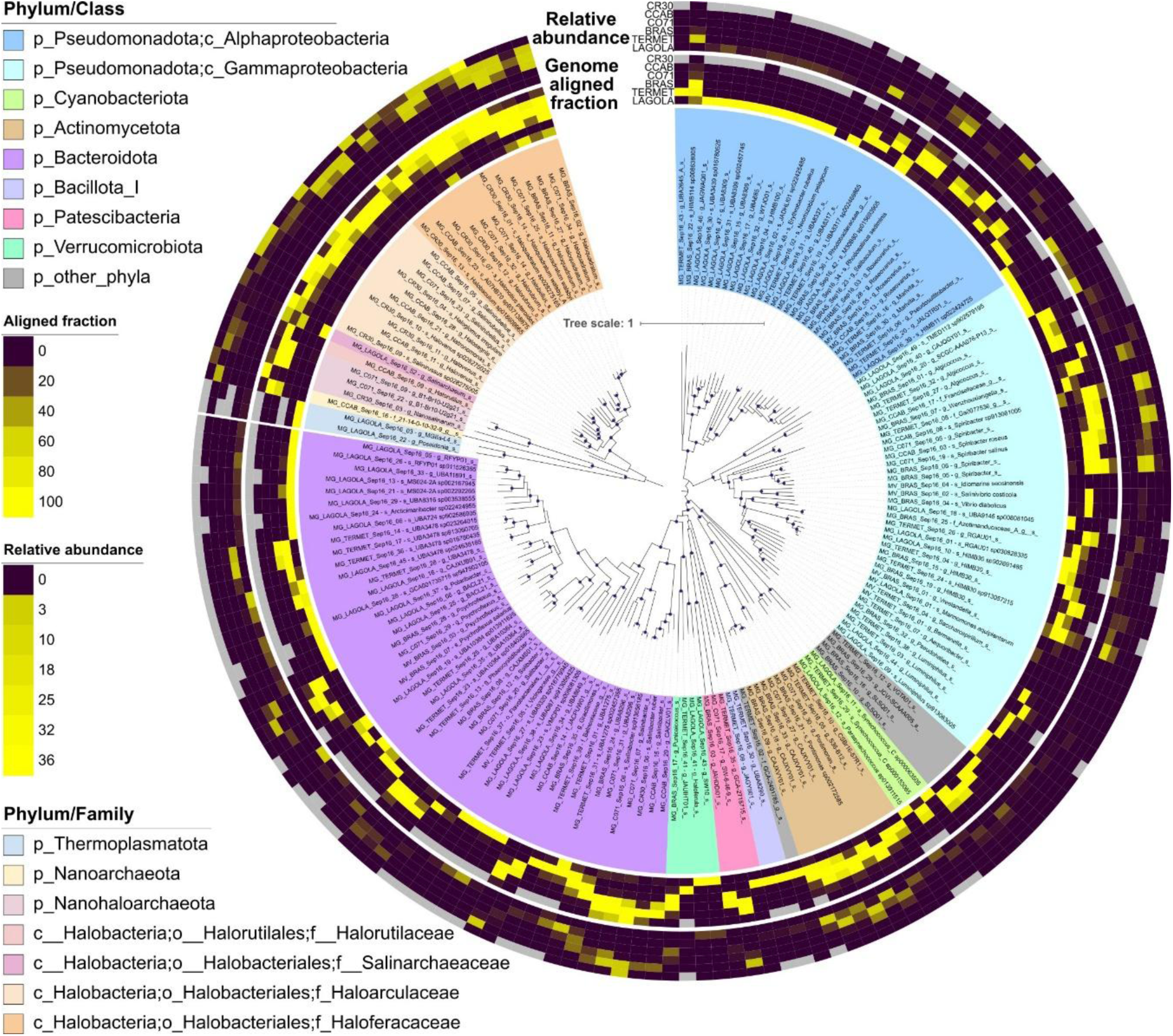
Phylogenomic reconstruction of the 170 dereplicated MAGs recovered across the salinity gradient. Maximum-likelihood phylogenomic tree inferred from concatenated alignments of 120 universal single-copy marker genes for Bacteria and 53 for Archaea (LG+F+R5 and LG+R8 chosen substitution models, respectively), using IQ-TREE with ultrafast bootstrap support (1,000 replicates); nodes with bootstrap values ≥95% are indicated by filled purple circles. MAGs were dereplicated at 95% average nucleotide identity (ANI) using dRep, retaining one representative per genomospecies cluster. Tip labels indicate MAG identifiers followed by their closest GTDB R220 taxonomic assignment. Inner color blocks indicate phylum/class (Bacteria) or phylum/family (Archaea) affiliation, as shown in the legends. Outer ring heatmaps show, for each of the six sampling sites in order of decreasing salinity from outermost to innermost ring (CR30, CCAB, CO71, BRAS, TERMET, LAGOLA), the relative abundance (TPM, outer ring per site) and genome-aligned fraction (inner ring per site) of each MAG, determined by mapping trimmed metagenomic reads with CoverM (≥95% nucleotide identity, ≥50% read aligned fraction). Tree scale bar represents one substitution per site.

Genomospecies presence-absence along the salinity gradient was determined using the metapresence tool with default parameters after comparison with other methods (Figure S6, Supplementary Text). Those from classes *Rhodothermia*, *Gammaproteobacteria*, *Bacteroidia*, *Actinomycetes, Verrucomicrobiae*, and *Halobacteria* were detected throughout most of the salinity gradient (Figure S7), consistent with previous observations [96]. Under more stringent mapping thresholds (50% aligned fraction), *Gammaproteobacteria*, *Alphaproteobacteria*, and *Bacteroidia* were absent from the highest salinity sample, suggesting these groups approach their physiological limits under near-saturation conditions. In contrast, several taxa with ultra-small cells, including *Nanosalinia*, *Paceibacteria* (c_Minisyncoccia in GTDB R226), and *Nanoarchaeia* (c_Nanobdellia in GTDB R226), showed distributions restricted to higher salinities, consistent with observations from other saltern systems [97]. *Cyanobacteriia* were confined to lower salinity environments, reflecting osmotic constraints on oxygenic photosynthetic microorganisms [98].

Nearly half of the genomospecies (46.5%) were restricted to single ponds (Figure S6). Only two MAGs were detected across five sites and with a wide horizontal genome coverage (Figure S8): MAG MG_BRAS_Sep16_21, affiliated to *Pontimonas* sp002172585, spanned from the marine sample LAGOLA to the hypersaline CCAB, a cross-gradient distribution from 3.6 % to 31.6%, expanding the previously reported moderate halophilic nature of the genus (1-9% sea salts tolerance [99]); similarly widespread, MAG MG_BRAS_Sep16_27, affiliated with *Haloquadratum*, spanned from LAGOLA to CR30 (excluding TERMET) and clustered within a group of four unclassified genomospecies (i.e., with no related genome at 95% ANI in either the dataset or GTDB R220). These unclassified *Haloquadratum* MAGs in this study showed their highest abundances at intermediate to high salinities (12-31.6% salts), contrasting with previously described *Hqr. walsbyi* isolates and the only one *Hqr. walsbyi* MAG recovered here, which peaked at 20.4-39.0% salinity (Figure 3). While cultured *Hqr. walsbyi* requires >14% salts for growth with optimal conditions at 18-36% salinity [100], these unclassified species appear adapted to somewhat lower salinities, representing an ecologically significant expansion of the range typically associated with this extreme halophilic genus. This broader salinity tolerance is consistent with two recent findings: the replacement of dominant *Hqr. walsbyi* populations by adapted congeneric species under strong osmotic dilution stress [101], and the identification of a single globally dominant *Hqr. walsbyi* genomovar whose abundance was further enhanced under moderate recurrent dilution disturbances [102]. Together, these results suggest that salinity range partitioning may be a broader ecological strategy within the genus.

### vOTU recovery and salinity-driven structuring of viral communities

Analysis of viral contigs yielded a dataset of 55,691 relaxed vOTUs ≥1 kb (36,801 identified by the MVP pipeline and 18,884 recovered through mapping against 27 extra viromes from the same hypersaline system), of which 8,139 constituted conservative vOTUs ≥5 kb (7,075 from the conservative MVP, plus 1,064 from virome mapping), after applying the identification and clustering strategies (see Materials and Methods; Table S5). vOTU presence-absence was determined using a 50% genome aligned fraction threshold after comparison with other thresholds (Figure S9; see Supplementary Text). Of the total relaxed dataset, 63.85% of vOTUs were classified as *Caudoviricetes*. Differential recovery of vOTUs between viromes and metagenomes provided insights into viral life cycle stages at the time of sampling (Figure S10): 36 vOTUs were enriched in viromes (mostly from TERMET), consistent with active production as extracellular particles, while 2,229 vOTUs were enriched in metagenomes (mostly from CCAB and CR30). These findings support the complementary use of metagenomes and viromes for characterizing viral communities: while differential abundance analysis captures vOTUs present in both library types at distinct replication stages, 28.8% of vOTUs in the relaxed set were detected exclusively in viromes, reflecting potentially active extracellular viral populations that would be missed by metagenome-only approaches [103–105].

Protein-based clustering revealed distinct viral communities structured by salinity. A large, interconnected network of 15,760 vOTUs along with reference genomes (Figure 4A), dominated by sequences from low-to-intermediate salinities (Super Viral Clusters SVC_1 to SVC_4; Figure 4A-B), was mainly composed of caudoviruses with predicted hosts within *Pseudomonadota* and *Bacteroidota*. In contrast, high-salinity sites harbored multiple discrete clusters (SVC_5 to SVC_18; Figure 4A-B), present in two or more sites, with predicted hosts primarily from *Halobacteriota* and *Bacteroidota*. These clusters were constituted by vOTUs mostly classified as unknown and captured by the virome-mapping strategy.

**Figure 4.**
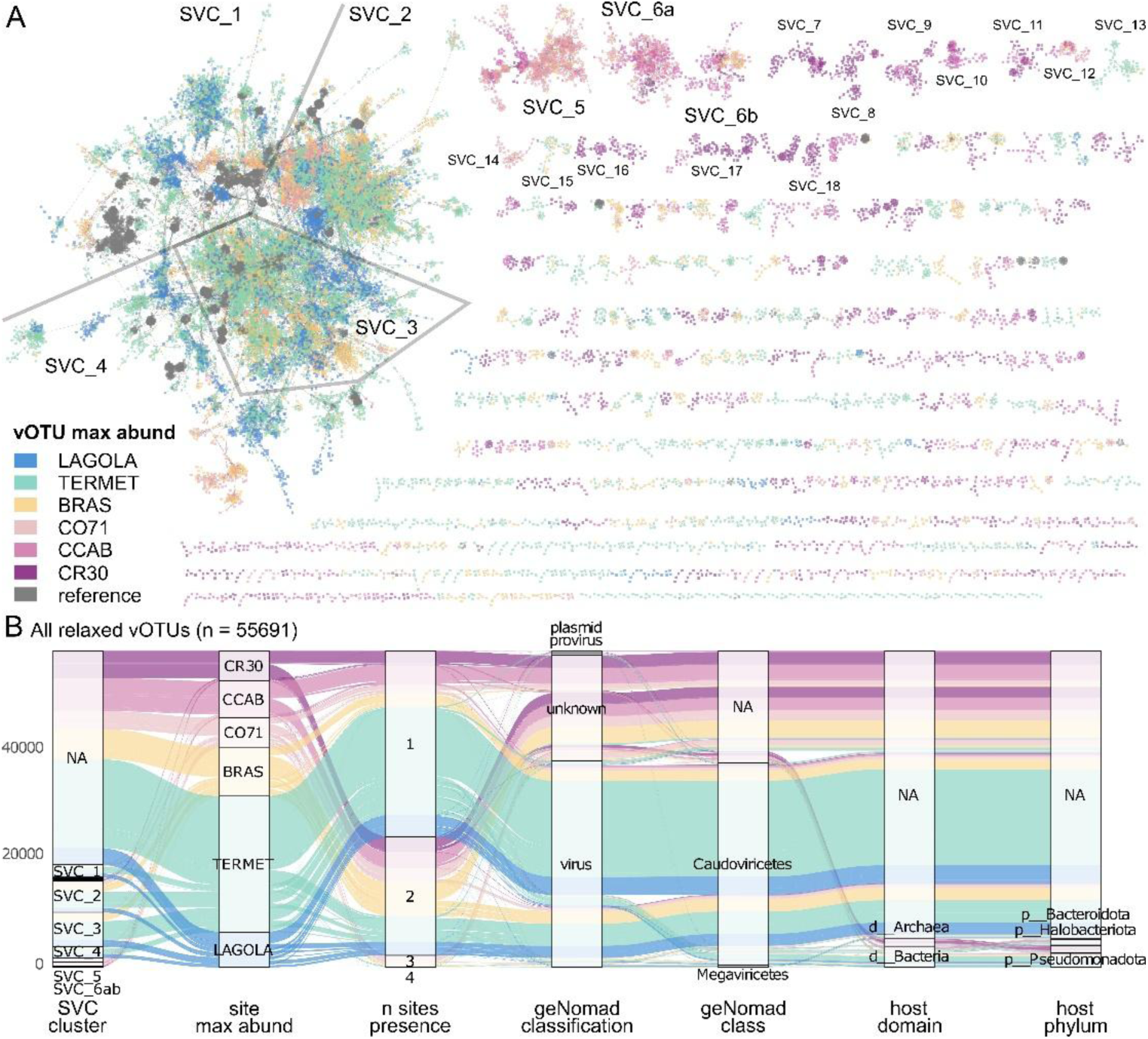
Protein-sharing network and ecological metadata of vOTUs across the salinity gradient. (A) Protein-cluster-based network of 15,760 relaxed vOTUs constructed with vConTACT2, using shared predicted protein clusters as edges. Each node represents one vOTU, colored by the site of its highest abundance (TPM). Reference genomes from the ProkaryoticViralRefSeq211-Merged database are shown in grey. Major Super Viral Clusters (SVCs) are custom groups with high numbers of vOTUs for further descriptions. (B) Alluvial diagram summarizing metadata associations for all 55,691 relaxed vOTUs. Flows connect, from left to right: SVC cluster membership; site of maximum abundance; number of sites where the vOTU was detected (presence at ≥50% genome-aligned fraction); geNomad sequence classification (virus, provirus, plasmid, or unknown); geNomad taxonomic class; predicted host domain; and predicted host phylum based on PHIST analysis against dereplicated MAGs. NA indicates that the corresponding metadata was unavailable or could not be assigned for that vOTU.

Several high-salinity clusters represented novel viral groups (Figure 4A-B). SVC_5 contained 755 vOTUs without cultured references, predicted to infect *Haloferacaceae*, *Rhodobacteraceae*, and *Salinibacteraceae*, while SVC_6a (454 vOTUs) included 12 reference viruses infecting *Haloferax*, *Halorubrum*, *Haloarcula*, and *Halogranum* [106], alongside additional references targeting *Haloquadratum* and *Gammaproteobacteria*. SVC_6b (202 vOTUs), mostly from CCAB and CO71, included the *Halorubrum* tailed virus HRTV-4 [106] and vOTUs predicted to infect *Salinibacter* or *Haloquadratum*. Together, these clustering patterns indicate that viral diversity and host range shift markedly across the salinity gradient, with high-salinity environments harboring compositionally distinct and largely uncharacterized viral assemblages.

### Niche breadth of viruses and hosts increases with salinity

To identify potential virus-host relationships within samples without relying on external databases, three complementary approaches were applied (Table S6): PHIST with a stringency filter of 3 or more shared k-mers, PHIST without filtering, and PHIST-WiSH common pairs, yielding 3,952, 5,590, and 1,369 relaxed vOTU-genomospecies pairs, respectively. The stringent PHIST subset was used as the primary dataset, with the remaining subsets serving to assess robustness of the results (Figures S11-S15). These methods of virus-host relationship prediction were further supported by an analysis against experimentally confirmed pairs from genera associated with MAGs present in the samples: *Haloquadratum spp.*, *Sal. ruber*, and *Synechococcus* sp. WH7803, with available sequenced viruses in databases (Table S7), recovering 85.48% precision (correct assignments) but limited recall (6.39% of total possible correct assignments) with the ≥3 k-mer threshold, suggesting that stringency improves reliability at the cost of sensitivity.

Considering the presence-absence patterns of each member of the virus-host pairs (Figure 5A-B), three predominant co-occurrence patterns were identified, and the taxonomic composition of these groups differed markedly across the gradient (Figure S16). The 1-*vs*-1 and 2-*vs*-1 groups, that is, those virus-host pairs where the vOTU was detected in one or two sites, while the corresponding host was detected only in one site, respectively, were dominated by *Caudoviricetes* from low-salinity Super Viral Clusters (SVC_1 to SVC_4), predicted to infect hosts characteristic of marine and low-salinity environments, including *Roseobacter*-affiliated genera such as HIMB11, *Roseovarius*, and *Marinomonas*. Roseophages support this tendency as they are characterized by narrow host ranges and high genetic diversity, with known isolates targeting specific strains and failing to infect closely related species (e.g., [107]). Other predicted low-salinity hosts included other *Alphaproteobacteria* genera such as HIMB30 and *Luminiphilus*. In contrast, the 3-*vs*-4 group, with vOTUs detected in three samples and their corresponding hosts found in four sites, was characterized by taxonomically unclassified vOTUs and some affiliated with SVC_8 (Figure 4A), predominant at intermediate-to-high salinity sites (BRAS to CR30), with predicted hosts almost exclusively from *Halobacteriota*, *Salinibacter*, and *Spiribacter*, mirroring virus-host associations previously described in arid-region salt lakes [8]. Analysis of non-co-occurrent pairs, those where only one member of the pair was detected in a site, revealed contrasting patterns across the gradient (Figure 5C): high-salinity sites (CCAB to CR30) contained more non-co-occurrent MAGs than vOTUs, while low-to-medium salinity sites (LAGOLA to CO71) showed the opposite pattern. This asymmetry, supported by a significant difference in site distribution between entities (Wilcoxon paired test, p < 0.05; Figure 5D), could reflect a salinity-dependent structuring of viral communities, consistent with patterns reported across wetland and saline lake habitats with contrasting salinity regimes [8,108]. Alternatively, it may partly result from differences in virus-host prediction efficacy across the gradient, as computational tools may identify with higher confidence associations at higher salinities where host diversity is lower and community composition is more constrained.

**Figure 5.**
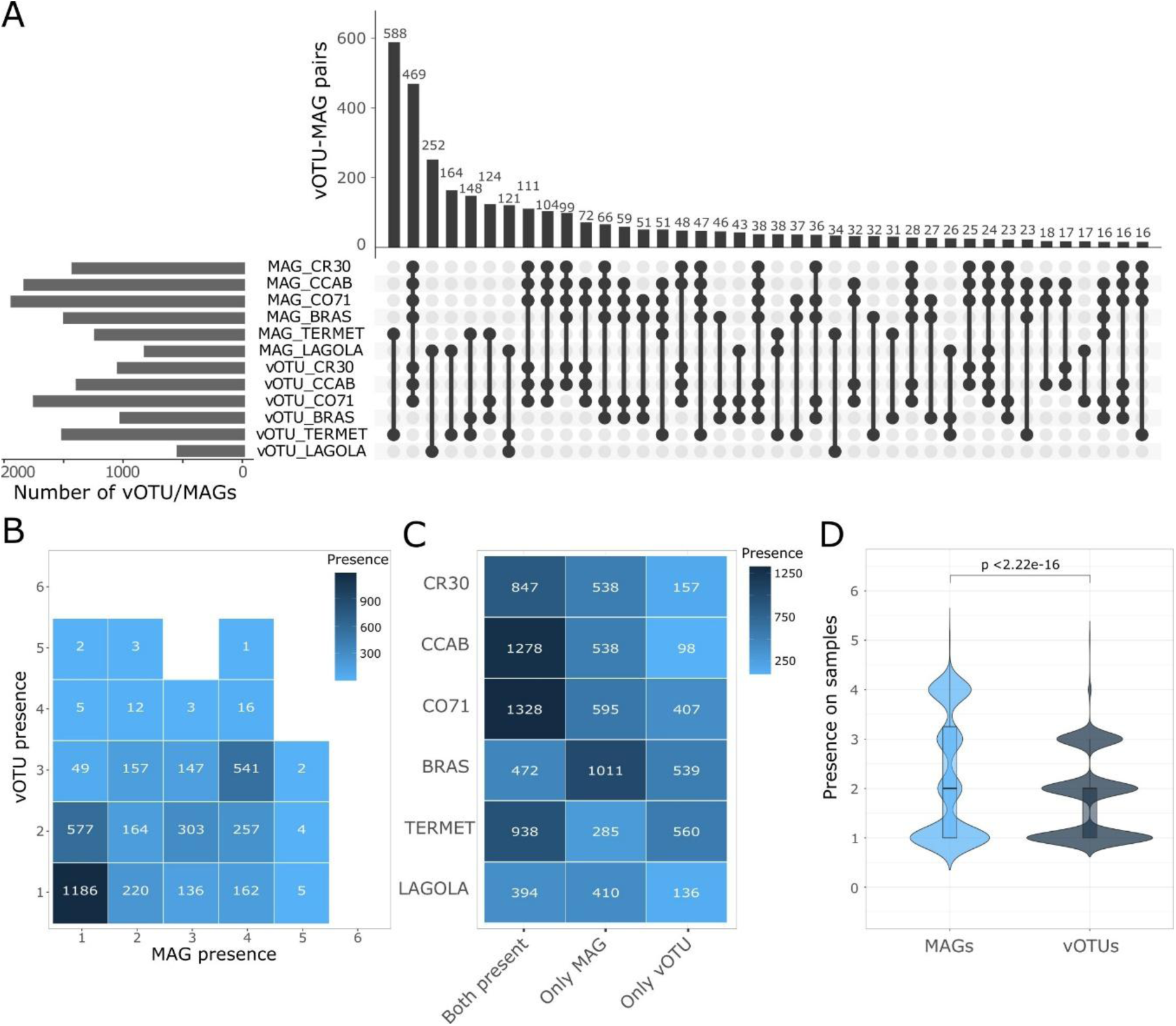
Presence-absence patterns of predicted virus-host pairs across the salinity gradient. Virus-host pairs were predicted using PHIST with a stringency threshold of ≥3 shared k-mers (n = 3,952 relaxed vOTU-MAG pairs). MAG presence was determined with metapresence and vOTU presence by ≥50% genome-aligned fraction across both metagenomic and viromic libraries. (A) UpSet plot displaying the distribution of predicted pairs by per-site presence-absence of each member. Horizontal bars (left) show the total number of MAG or vOTU detected at each sample. Vertical bars (top) show the count of pairs sharing a given combination of site detections, with filled circles indicating which pairs of MAGs and vOTUs contributed to each intersection. (B) Two-dimensional histogram of the number of sites where each vOTU (y-axis) and its cognate MAG (x-axis) were independently detected, across the six sampling sites. Cell values indicate the number of pairs falling in each combination. (C) Site-level co-occurrence matrix showing, for each site, the count of predicted pairs where both members were detected (Both present), only the MAG was detected (Only MAG), or only the vOTU was detected (Only vOTU), based on the presence-absence thresholds described above. (D) Violin plots comparing the number of sites where MAGs and their cognate vOTUs were independently detected. Boxes show median and interquartile range. Statistical significance was assessed with a Wilcoxon signed-rank test (p-value shown).

Abundance relationships between vOTUs and their putative hosts revealed distinctive dynamics across the salinity gradient, with linear regression models of log_10_-transformed TPM values showing significant positive slopes for all sites except TERMET (Figure 6A), increasing progressively from LAGOLA to CR30, and consistent across most data subsets (Figures S17-S21). While direct abundance relationships alone are not reliable as *de novo* virus-host relationships predictors and can be biased by prediction methodologies [109–110], they have been proposed as proxies for viral infection strategies. Both kill-the-winner (KtW [111]) and piggyback-the-winner (PtW [112]) dynamics predict positive virus-host abundance correlations but differ in slope: KtW predicts strong near-linear tracking where viruses suppress dominant hosts, while PtW predicts sublinear increases where proviruses replicate alongside hosts at high host densities. The progressive increase in slope toward higher salinities could therefore suggest a gradual shift from mixed strategies at lower salinities toward a predominance of lytic dynamics at the highest salinities, a pattern consistent with the observed decrease in integrative lysogenic markers abundances towards CR30 (Figure S22). However, this interpretation should be considered cautiously: experimental evidence from *Sal. ruber* and its isolated viruses suggests that high viral pressure can promote episomal virus maintenance and host population recovery through pseudolysogeny, enabling virus-host coexistence rather than host killing under high virus-to-host ratios [113], while co-evolutionary arms-race dynamics at the host strain level add further complexity to these community-scale patterns [104].

**Figure 6.**
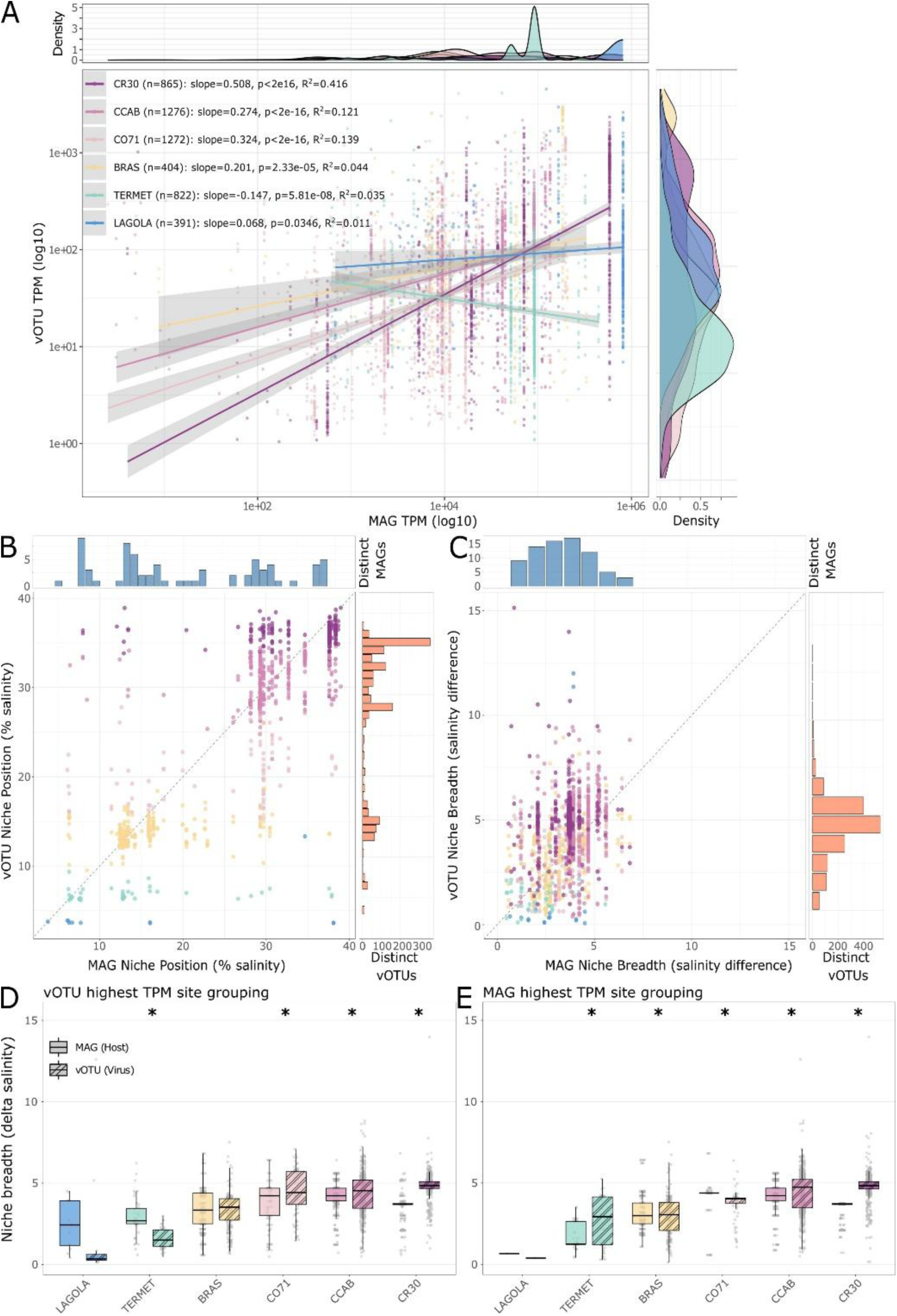
Abundance relationships and niche dynamics of predicted virus-host pairs along the salinity gradient. Virus-host pairs were predicted using PHIST with a stringency threshold of ≥3 shared k-mers (n = 3,952 unique pairs; 5,030 total site-level observations), quantified by mapping trimmed reads with CoverM (≥95% nucleotide identity, ≥50% read aligned fraction). (A) Linear regression of log₁₀-transformed TPM abundances of vOTUs versus their cognate MAGs, fitted independently for each sampling site for which the vOTU had the highest abundance. Pairs where either member had no mapped reads at a given site are excluded. Regression statistics per site include the number of pairs (n), slope, coefficient of determination (R²), and p-value. Marginal density plots show the distribution of MAG (top) and vOTU (right) abundances. (B-C) Niche characteristics of virus-host pairs, calculated for the subset of 1,480 pairs with detections in at least two sites (MAGs: ≥2 metagenomic samples; vOTUs: ≥2 samples across metagenomic and/or viromic libraries). (B) Niche position, defined as the abundance-weighted mean salinity optimum, shown for each vOTU-MAG pair. (C) Niche breadth, defined as the abundance-weighted standard deviation of salinity around the niche position, shown for each pair. Marginal histograms show the number of distinct MAGs (top) and vOTUs (right) across 1% salinity intervals; dot colors indicate the site of highest vOTU abundance. (D-E) Pairwise comparison of niche breadth between vOTUs and their cognate MAGs, grouped by the site of highest TPM for vOTUs (D) or MAGs (E). Boxes show median and interquartile range. Asterisks indicate statistically significant differences between paired vOTU and MAG niche breadths within each site (Wilcoxon signed-rank test, p < 0.05).

While presence-absence analysis captures all predicted virus-host pairs regardless of detection frequency, niche modeling requires abundance values across at least two sites and therefore focuses on a subset of pairs with sufficient distributional data. Notably, the 1-*vs*-1, 2-*vs*-1, and 1-*vs*-2 co-occurrence groups, predominantly from low salinities and likely representing highly specialized single-pond associations such as those for roseophages and their *Roseobacter*-like hosts (Figure S16), are excluded from niche analyses by this constraint, though they remain ecologically important components of the low-salinity viral community. For the remaining pairs, abundance distributions across the salinity gradient revealed two key niche characteristics. Niche position, defined here as the salinity at which abundances should peak, showed remarkable agreement between vOTUs and related hosts (Figures 6B and S21; Pearson’s r = 0.846, p < 0.001), with a strong correlation indicating consistent positioning despite a few deviated pairs, mostly hosts with low-to-medium salinity niches paired with vOTUs favoring high salinities. These patterns reflect the niche position specificity for viruses and hosts, as has been shown in other systems [114], and which may reflect co-evolutionary adaptation to the salinity gradient [115].

Niche breadth (Figure 6C), defined as the salinity range centered around the niche position and weighted by abundances, showed consistently higher mean values for vOTUs relative to their cognate hosts across all relaxed datasets (4.28 ± 1.61% *vs.* 3.65 ± 1.10%, p = 9.81e-61, Wilcoxon Signed-Rank Test), though this difference was not significant in conservative datasets (p > 0.05), suggesting it may be partly attributable to the shorter vOTU lengths or greater number of pairs in relaxed sets. When analyzed by site, vOTUs consistently showed significantly broader niches than their hosts at CR30 across all datasets, with the opposite pattern observed at TERMET, while intermediate salinities showed inconsistent results depending on the grouping used (Figure 6D-E; Table S7). More robustly, site-mean niche breadths of both vOTUs and hosts increased significantly with salinity (Spearman r = 0.83-1.00, p < 0.05), and at the individual level, vOTU breadth correlated positively with salinity in all six datasets (r = 0.36-0.66, p < 0.001), while host breadth showed a weaker and less consistent relationship (Figure S24). Taking this into account, increasing niche breadth toward environmental extremes has been described for prokaryotes [116], suggesting that the broader salinity tolerance of halophiles relative to marine microorganisms extends to their associated viral communities, with both halophilic hosts and viruses showing progressively broader ecological ranges toward hypersaline conditions. This pattern may further reflect the well-documented asymmetry in microbial dispersal along salinity gradients, where colonization from low-to high-salinity environments is more constrained than the reverse [117], potentially limiting the establishment of marine-adapted viruses and their hosts at higher salinities (Figure S25).

### vOTUs tetranucleotide frequencies group due to a combination of host taxonomy and site provenance

To assess the effect of salinity on the genetic distance among either viruses or prokaryotic genomes, two strategies were employed. First, linear regression analysis was performed between ANI and absolute salinity differences between the respective origin ponds for genome pairs (either viral contigs or MAGs). For all viral contigs with >70% ANI, salinity differences explained little variation in genetic distance (low R² values of 0.0002 and 0.0003; Figure S26A-B). In contrast, prokaryotic MAGs showed higher, albeit low, positive association (slope = 0.27, R² = 0.06; Figure S26C), indicating a higher salinity effect on prokaryotic genetic distance than on viruses. Thus, although salinity gradients have a stronger influence on prokaryotic genetic divergence, the overall explanatory power remains limited for both viruses and prokaryotes.

For the second strategy, alignment-free methods were used due to their power to determine patterns of sequence clustering [118], as a proxy of evolutionary relationships, and to detect the influence of metadata on these patterns. Principal coordinate analysis (PCoA) of Bray-Curtis dissimilarities between tetranucleotide frequencies from all relaxed vOTUs with host assignments (n = 3,952) showed a distribution mostly driven by the GC content over PC axis 1 (Mantel r = 0.921; Figure 7A), while salinity showed a contribution mostly over PC axis 2 (Mantel r = 0.127). When considering discrete variables, the host class was more explicative (R^2^ = 0.306) than the site where they were found with higher abundances (Highest TPM Site; R^2^ = 0.187); however, highest TPM site was more explanatory than host phylum (R^2^ = 0.083). Therefore, there was a higher influence of the host taxonomy rather than the site with highest abundance, but this was dependent on the taxonomic level considered, which is sustained considering other similarity matrices (Figure 7B) and other data subsets (Figure S27). Similar patterns have been seen regarding salinity in an estuarine system in which host genus had a greater effect than salinity on genomic properties of viruses [115]. The current class-level differences are seemingly driven by division of high-and low-GC members from two phyla: *Pseudomonadota* (*Alpha*- and *Gammaproteobacteria*, respectively) and *Bacteroidota* classes (*Rhodothermia* and *Bacteroidia*) (Figure 7A) which show distinct distribution patterns in the salinity gradient (Figure S7) and in other studies [119]. Therefore, viral genomic composition clusters primarily according to the taxonomy of their hosts at or below class level, consistent with the previously described tendency of viral sequences to group by host phylum, reflecting shared evolutionary history between viruses and their hosts across broad taxonomic ranks [11]. This pattern carries an additional contribution of salinity provenance, likely mediated by the salinity-driven distribution of distinct host classes across the gradient rather than by a direct environmental effect on viral genomic composition, in agreement with the stronger influence of salinity on prokaryotic than on viral genetic distances observed above (Figure S26).

**Figure 7.**
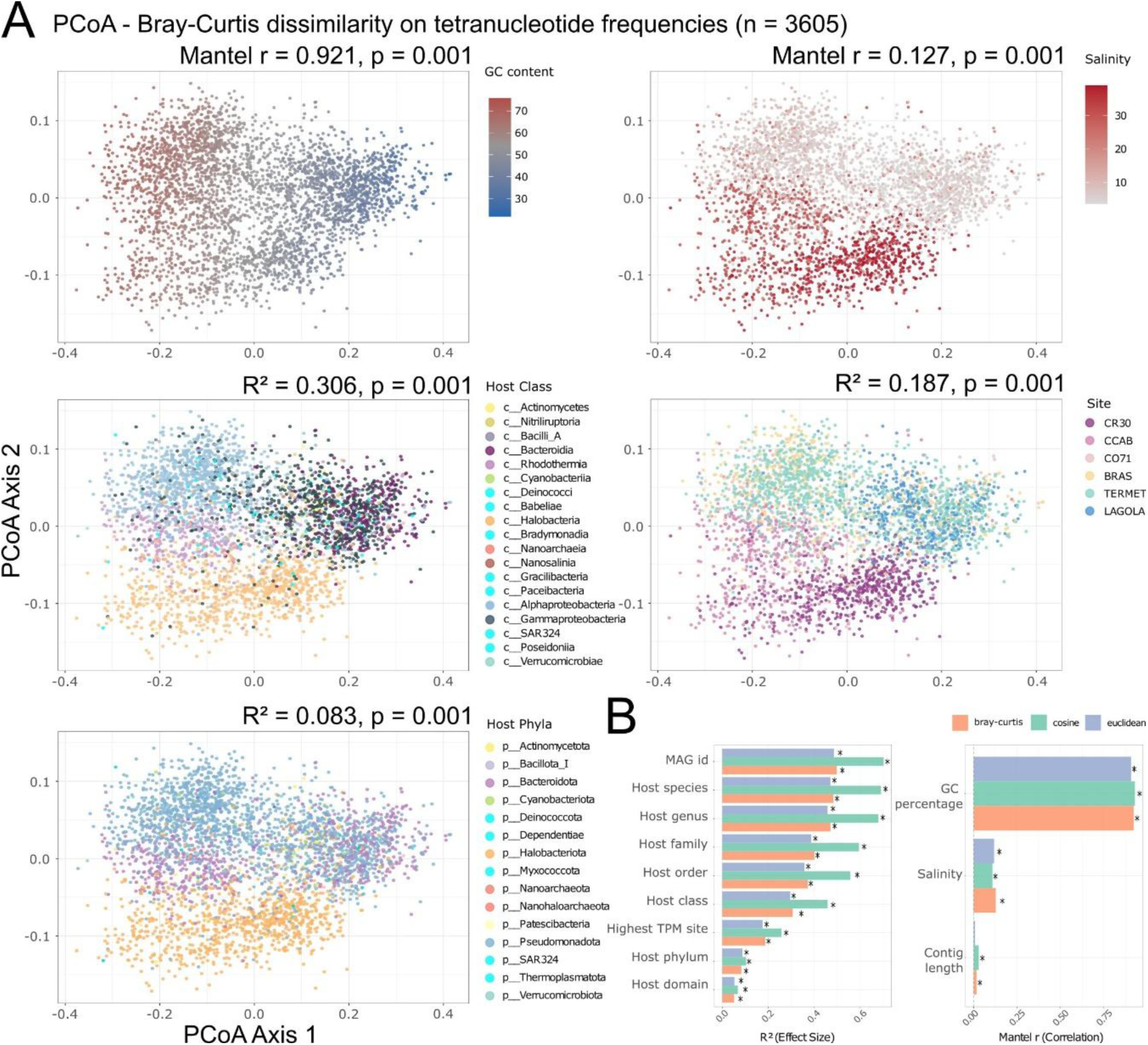
Tetranucleotide frequency patterns of vOTUs and drivers of compositional variation. (A) Principal Coordinate Analysis (PCoA) based on Bray-Curtis dissimilarity of vOTU tetranucleotide frequencies (n = 3,605 vOTUs with host assignment via PHIST ≥3 shared k-mers). Points are colored by GC content (upper left), salinity of the site of highest vOTU abundance (upper right), predicted host class (middle left), sampling site (middle right), and predicted host phylum (lower left). Mantel tests were used for continuous variables (GC content and salinity) and PERMANOVA for categorical variables (host taxonomy and sampling site). (B) Effect sizes of all tested variables on vOTU tetranucleotide frequency composition using three distance metrics: Bray-Curtis (orange), cosine (green), and Euclidean (blue). Left panel shows R² effect sizes from PERMANOVA for taxonomic and ecological variables ranked from higher to lower effect size; asterisks indicate significant associations (p < 0.05) after permutation testing. Right panel shows Mantel r correlation coefficients for continuous variables (GC percentage, salinity, and contig length) against the three distance matrices; asterisks indicate significant associations (p < 0.05).

## Conclusion

Overall, our findings support an “expanded host niche” scenario as the predominant virus-host interaction strategy along the salinity gradient, likely facilitated by high microdiversity among closely related hosts (e.g., *Sal. ruber* [120]) and congeneric species replacement (e.g., *Haloquadratum* [101]), as well as by the enhanced physical stability of viral particles under hypersaline conditions [121], and dynamic shifts in viral life cycle strategies across salinity gradients [8]. However, “nested host niche” examples, where viruses occupy restricted niches within individual hosts, were also detected mostly at lower salinities, indicating that diverse virus-host interaction strategies coexist along the gradient. Salinity emerges as a fundamental environmental parameter shaping not only virus-host diversity but also the niche breadth of both entities, while host taxonomy at the class and lower levels determines the evolutionary signal in viral genomic composition. These results advance our understanding of how environmental gradients and host evolutionary history jointly structure viral ecological niches, with specific implications for predicting viral diversity and infection dynamics in extreme environments and interpreting virus-host co-evolutionary patterns across other physicochemical gradients.

## Data availability

Raw reads of the 12 samples and 82 MAGs are available in the NCBI SRA and genome databases, respectively, under the NCBI BioProject accession PRJNA1465206. Remaining MAGs with < 90 % completeness are available in the figshare repository DOI 10.6084/m9.figshare.32676357. Code for niche calculation through three methodologies is available on the GitHub repository https://github.com/jaalcort/nichePosBdt_calculator.

## Author Contributions (CRediT)

Jaime Alcorta: Methodology, Formal analysis, Investigation, Writing - Original Draft, Visualization. María Dolores Ramos-Barbero: Methodology, Investigation, Writing - Review & Editing. Fernando Santos: Conceptualization, Writing - Review & Editing, Visualization, Funding acquisition. Josefa Antón: Conceptualization, Resources, Writing - Review & Editing, Supervision, Funding acquisition.

