## Supplementary information and figures for "Coupled effects of salinity and host phylogeny on niche breadth and viral evolution from seawater to salt saturation"

### Supplementary text

*Presence-absence across mapping methods*

No universally accepted parameters exist for determining MAG or reference genome presence-absence in metagenomic samples through read mapping. Castro et al. [122] provided the first metagenomic systematic assessment using imGLAD, finding confident detection (probability of presence >99%) at sequencing breadths of 3-4% of the total genome when mapping at >95% nucleotide identity. Notably, other tools evaluated in the same study, including PathoScope and MetaPhlAn, required at least 10% genome coverage to achieve comparable specificity and sensitivity with spiked genomes in real metagenomes. The issue was further addressed by Sanguineti et al. [55], who developed metapresence, a tool that incorporates additional metrics beyond simple aligned fraction: the Breadth-Expected Ratio (BER) and the Fraction of Unexpected Gaps (FUG), which together account for both coverage depth and the spatial distribution of mapped reads across the genome. The underlying mapping strategy uses Bowtie2 in sensitive mode, corresponding to approximately 98% identity for read recruitment.

To select the most appropriate threshold for our dataset, we compared MAG distribution patterns (Figure S6) across samples using four aligned fraction cutoffs with read mapping at 95% of identity (AF0, AF10, AF50, and AF80) alongside metapresence run under default parameters (BER > 0.8, or BER > 0.8 and FUG > 0.5 for low-coverage genomes) and a more restrictive configuration (BER > 0.7, or BER > 0.7 and FUG > 0.45). Although presence-absence patterns were broadly consistent across approaches, the closest agreement in detection numbers was observed between metapresence with default parameters and the AF10 threshold, with less patchy distributions than AF50. Intersection analysis (Figure S6) further supported metapresence default parameters as a robust framework for establishing prokaryotic genome presence-absence, and this threshold was adopted for comparisons with vOTU mapping data, which itself exceeds the 3-4% detection threshold established by Castro et al. [122].

Unlike prokaryotic genomes, several large-scale viral metagenomic studies have adopted aligned fraction thresholds ranging from 10%, as used in the IMG pipelines [123-124], to 75%, as used in the TARA ocean pipeline [125]. However, the appropriate threshold varies substantially with genome size and completeness. Ramos-Barbero et al. [126] explicitly argued against the 10% threshold for crAss-like phages, noting that for genomes of ~100 kb, a single shared gene could represent 10% of the genome and produce false positives; in that study instead it was adopted a 40%

horizontal coverage cutoff, supported by observed average coverages of ~47% in human feces and ~37% in wastewater.

Here, when comparing vOTU detection numbers across the different aligned fraction and metapresence settings for the three vOTU datasets (Figure S9), most detections were shared across all methods. However, for the larger dataset of the relaxed vOTUs, the metapresence settings under both configurations detected fewer vOTUs than the AF80 threshold, while for the conservative vOTU set the metapresence values were closer to AF50 and AF80. In contrast to the prokaryotic analysis, where metapresence default parameters were most concordant with AF10, for vOTUs the metapresence settings aligned more closely with higher aligned fraction thresholds, indicating that direct application of the same thresholds across cellular and viral genomes requires caution and further validation.

Given that the vOTUs from this study have a minimum length of 1 kb, an aligned fraction of 10% would correspond to only 100 bp of covered sequence, which is insufficient to reliably confirm genome presence and is susceptible to false positives from conserved genomic regions shared across multiple vOTUs. Therefore, consistent with the reasoning of Ramos-Barbero et al. [126] for larger phage genomes, we selected an aligned fraction threshold of 50% (AF50) for vOTU presence-absence analyses, which represents a more biologically meaningful coverage level for contigs of this size range and is consistent with the higher thresholds observed to perform comparably to metapresence settings in the conservative vOTU dataset (Figure S8). It is worth mentioning that most analyses and figures also were performed with the AF10 threshold for the vOTUs, being mostly consistent with the results presented with the AF50 threshold.

*Comparison with experimental virus-host pairs*

To validate predicted virus-host relationships with experimentally confirmed associations, we identified representative genomes for our MAG genomospecies using the GTDB version R220 taxonomy from the GTDB database (https://gtdb.ecogenomic.org/). For representative genomes derived from culture-based analyses, associated phages were retrieved from literature and the NCBI nucleotide database using "genus_name phage" queries. Reference viral genomes (Table S7) were then compared through BLASTn searches to relaxed vOTUs (≥90% identity, ≥85% horizontal coverage) determining possibly related viruses from this study to those references.

From 56 MAGs with species affiliation, 27 were associated with a defined genus and 18 contained a cultured strain as the representative genome (Table S4). Sequenced viruses were found for six of them (Table S7): *Haloquadratum walsbyi* (n = 27), *Salinibacter ruber* (n = 18), *Synechococcus* sp. WH7803 (n = 14), *Erythrobacter rubellus* (n = 5), *Vibrio diabolicus* (n = 2), and *Salinivibrio costicola* (n = 1). A comparison against these references identified 170 relaxed vOTUs associated with *Hqr. walsbyi* viruses (n references = 25 [26]), 19 with *Sal. ruber* viruses (n references = 12 [23,104]), and 15 with a *Synechococcus* sp. WH7803 virus (S-BM1 [127]). From these vOTUs, only 6.39% were associated with a predicted host using PHIST, with 85.48% precision; true positives were restricted to *Hqr. walsbyi* assignments, with false positives including assignments from *Hqr.* to g__HIMB30-s and from *Sal.* to g__Halorutilus, and no true positives were recovered for *Salinibacter* or *Synechococcus*. Although the original PHIST study reported 64.9% accuracy at the family level [71], the more stringent ≥3 k-mer cutoff applied here improves prediction reliability at the cost of sensitivity, likely missing virus-host relationships where tetranucleotide patterns diverge substantially between virus and host pairs.

*Niche prediction based on weighted-abundances*

Niche position and niche breadth were calculated using a custom abundance-weighted approach (called here column-sum normalisation) following the conceptual framework of the ecospat package by Di Cola et al. [73]. The column-sum and ecospat implementations differ in weight normalisation and variance denominator, as described below.

Let $s_j$ denote the value of the environmental variable of interest (here, salinity in % total dissolved salts) for sample j, and $t_{ij}$ the normalised abundance of entity i (vOTU or MAG) in sample j expressed as a measure enabling cross-sample comparisons (RPKM, RPKG or TPM; see below), and $J_i$ the set of samples in which entity i was detected above the detection threshold (aligned fraction >50% for vOTUs; metapresence default for MAGs).

Abundance values used as weights must be normalised to enable meaningful comparisons across samples with different sequencing depths. Raw read counts are unsuitable because they reflect sequencing effort as much as biological abundance, and compositional relative abundances (proportions of total reads per sample) introduce dependencies between entities that distort weighted means when community composition changes across the gradient. TPM normalisation addresses both issues by scaling each entity's read count to a common library size of one million reads after correcting for gene or genome length, producing values that would be comparable across samples and proportional to true abundance differences between sites. All three methods described below use TPM as the abundance measure $t_{ij}$.

For a given entity i, **niche position** $P_i$ was defined as the abundance-weighted mean of the environmental variable (here, salinity) across all detected samples:

$$\text{[Equation 1]} \quad P_i = \sum_{j \in J_i} w_{ij} \cdot s_j$$

where $w_{ij}$ are the column-sum weights:

$$\text{[Equation 2]} \quad w_{ij} = \frac{t_{ij}}{\sum_{k \in J_i} t_{ik}}$$

Here k indexes all samples in the detected set $J_i$, so that $t_{ik}$ refers to the TPM of entity i in each sample k used to compute the normalising sum, ensuring that weights are proportional to the abundance of each entity at each site relative to its total across all detected samples, and sum to one.

This formula is structurally similar to that used by the ecospat function ecospat.nichePOSNB [73], as both compute a weighted mean of the environmental variable, using abundance as weights. However, because the two implementations differ in weight normalisation (column-sum vs max-normalisation over all sites), niche position estimates are not identical, though differences in the current dataset are generally small.

The ecospat niche position (Equation 1b) is computed as the weighted mean of the environmental variable (here, salinity) using max-normalised weights (Equation 2b). In Equation 2b, k indexes all samples including those where abundance is zero. Because most entities are absent from at least one site, the minimum term in Equation 2b is typically zero in practice, and the formula reduces to $t_{ij}$ divided by the maximum abundance of entity i. Unlike column-sum weights, max-normalised weights do not sum to one, so the position formula (Equation 1b) explicitly divides by their sum:

$$\text{[Equation 1b]} \quad P_i^{ecospat} = \frac{\sum_{j \in J_i} w_{ij}^{ecospat} \cdot s_j}{\sum_{j \in J_i} w_{ij}^{ecospat}}$$

$$\text{[Equation 2b]} \quad w_{ij}^{ecospat} = \frac{t_{ij} - min\ t_{ik}}{max\ t_{ik} - min\ t_{ik}}$$

**Niche breadth** was defined as the abundance-weighted standard deviation of the environmental variable (here, salinity) around the niche position. Both our column-sum implementation and ecospat share the same general structural formula for the weighted variance, but differ in how abundance weights are normalised and in the variance denominator, as detailed below:

$$B_i = \sqrt{\sum_{j \in J_i} w_{ij} \cdot \left(s_j - P_i\right)^2} \quad \text{[Equation 3]}$$

where $s_j$ is the value of the environmental variable for sample j (here, salinity), $P_i$ is the niche position of entity i (Equation 1), and $w_{ij}$ are the abundance weights. The column-sum breadth (Equation 3) uses weights from Equation 2, while the ecospat breadth (Equation 3b) uses max-normalised weights from Equation 2b and a different variance denominator.

The corresponding ecospat niche breadth (Equation 3b) uses the same squared-deviation structure but with max-normalised weights (Equation 2b) and a frequency-weight bias correction denominator of $\sum w^{ecospat} - 1$ (as implemented in Hmisc::wtd.var), rather than $\sum w = 1$ as in column-sum:

$$B_i^{ecospat} = \sqrt{\frac{\sum_{j \in J_i} w_{ij}^{ecospat} \cdot \left(s_j - P_i^{ecospat}\right)^2}{\sum_{j \in J_i} w_{ij}^{ecospat} - 1}} \quad \text{[Equation 3b]}$$

To quantify the practical impact of normalisation choice, column-sum and ecospat breadth estimates were compared across the full datasets of 25,808 relaxed vOTUs and 89 MAGs detected in at least two sites. The ecospat implementation produced systematically larger breadth estimates than column-sum normalisation across the full dataset (Figure S28), due to the combined effect of max-normalised weights and the sum(w) − 1 variance denominator. Entities detected at only two sites represented the majority of vOTUs (n = 14,755; 57.2%) and 31.5% of MAGs (n = 28); for these entities, ecospat breadths were particularly inflated relative to column-sum, with a mean ratio of ecospat to column-sum breadth of 4.95 for vOTUs and 6.69 for MAGs at two detected sites. Across the full datasets, the mean ratio of ecospat to column-sum breadth was 3.90 for vOTUs and 4.71 for MAGs, and the ratio decreased progressively as the number of detected sites increased, with ecospat and column-sum breadths approaching closer agreement at five to six detected sites.

To provide an independent reference for niche breadth estimation, a Gaussian response curve was fitted to the TPM abundance of each entity as a function of the environmental variable (here, salinity) across the sampling sites. Gaussian curve fitting was performed in R using the nls() function from the base stats package, implementing a nonlinear least-squares algorithm with the port optimisation method. The model fitted was (Equation 6):

$$TPM_{ij} \sim amp_i \cdot \exp\left(-\frac{1}{2} \cdot \left(\frac{s_j - \mu_i}{\sigma_i}\right)^2\right) \quad \text{[Equation 6]}$$

where $amp_i$ is the amplitude (maximum predicted abundance), $\mu_i$ is the value of the environmental variable at the abundance peak (equivalent to niche position), and $\sigma_i$ is the standard deviation of the Gaussian (used as the niche breadth estimate). To avoid convergence to local optima, a multi-start approach was implemented in which three starting values for σ were tested prior to fitting (range/8, range/4, and range/2 of the observed range of the environmental variable), and the fit with the highest $R^2$ was retained. The starting σ value that produced the best fit is reported as a diagnostic parameter. Bounds were set to ensure biologically plausible fits: amplitude was constrained to positive values with an upper bound of twice the observed maximum abundance (here, TPM) to allow the fitted peak to exceed observed values; μ was constrained within the observed range of the environmental variable at detected sites; and σ was constrained to a minimum of 0.001 units of the environmental variable to prevent degenerate spike-like fits, with an upper bound of twice the observed range. Convergence was assessed by successful completion of the nls() algorithm, and goodness of fit was evaluated using $R^2$ calculated as $1 - SS_{residual} / SS_{total}$ from the observed non-zero TPM values. Only entities detected at three or more sites were eligible for Gaussian fitting, as a minimum of three points is required to constrain the three parameters of the model; fits with $R^2 < 0.7$ were excluded from comparative analyses as insufficiently reliable. Under these criteria, 10,956 relaxed vOTUs (99.1% of those detected at three or more sites) and 61 MAGs (100%) yielded retained Gaussian fits. The lower overall convergence rate for vOTUs (42.8% of the full dataset) reflects the predominance of entities detected at only two sites, which are ineligible for Gaussian fitting, rather than poor model fit among eligible entities.

Among entities with successful Gaussian fits, the Gaussian σ was larger than column-sum breadth estimates at three detected sites, with mean ratios of 1.29 for vOTUs and 1.62 for MAGs, narrowing at four detected sites (mean ratio 0.97 for vOTUs and 1.38 for MAGs; Figure S29). This pattern reflects the sensitivity of Gaussian fitting to the shape and symmetry of the abundance distribution across the gradient. Column-sum and Gaussian breadths showed consistent positive correlations across site number groups (Spearman r = 0.68, 0.37, and 0.58 for vOTUs at three, four, and five detected sites respectively; r = 0.71 and 0.89 for MAGs at three and four sites), supporting the mutual consistency of the two approaches. Column-sum and ecospat breadths were also positively correlated among entities detected at three or more sites (Spearman r = 0.33, 0.41, and 0.46 for vOTUs at three, four, and five detected sites respectively; r = 0.07 and 0.36 for MAGs at three and four sites, neither reaching significance), though correlations were weaker than those observed between column-sum and Gaussian, consistent with the larger and more variable inflation factor of ecospat relative to column-sum. Ecospat and Gaussian breadths showed weak and inconsistent correlations across site number groups (Spearman $r = -0.14$, $-0.31$, and $0.09$ for vOTUs at three, four, and five detected sites respectively ($p < 0.001$, $p < 0.001$, and $p = 0.105$); $r = -0.002$ and $0.29$ for MAGs at three and four sites (both non-significant)), indicating that the two methods deviate from column-sum through independent mechanisms and in opposing directions depending on the number of detected sites. Three illustrative examples of vOTUs detected at all six sites are shown in Figure S30, with Gaussian fits characterised by $R^2$ values of 1.000 (MG_CR30_k119_84439), 0.846 (MG_BRAS_k119_224842), and 0.960 (MG_CO71_k119_7442), showing that entities with more symmetric unimodal distributions yield higher goodness of fit and closer agreement between Gaussian and weighted position estimates. Column-sum normalisation was therefore used for all niche breadth calculations reported in the main text, as it provides a direct and interpretable estimate of the realised niche breadth without the systematic inflation of ecospat or the unpredictable directional bias of Gaussian fitting at low site numbers. The three calculation methods are available at the GitHub repository: https://github.com/jaalcort/nichePosBdt_calculator/.

For vOTUs, metagenomic (MG) and viromic (MV) libraries were treated separately to avoid bias from differences in sequencing depth and viral capture efficiency between library types. Niche position was estimated independently per library type using Equations 1 and 2 applied to MG and MV samples separately, and a consensus position was obtained as a detection-rate weighted average (Equation 7):

$$\text{[Equation 7]} \quad P_i^{consensus} = \frac{d_i^{MG} \cdot P_i^{MG} + d_i^{MV} \cdot P_i^{MV}}{d_i^{MG} + d_i^{MV}}$$

where $d_i^{MG}$ and $d_i^{MV}$ are the detection rates of entity i in metagenomic and viromic libraries respectively, defined as the fraction of sites in which the vOTU was detected ($TPM > 0$). Only library types where the vOTU was detected in at least two sites contributed to the consensus estimate; vOTUs detected in only one library type were retained using the single-library estimate. The consensus niche breadth was then recalculated by pooling all detected samples from both library types around the consensus position:

$$\text{[Equation 8]} \; B_i^{consensus} = \sqrt{\sum_{j \in J_i^{MG} \cup J_i^{MV}} w_{ij}^{pooled} \cdot \left(s_j - P_i^{consensus}\right)^2}$$

where $w_{ij}^{pooled}$ are abundance weights computed from the pooled TPM values of both library types normalised by their combined column sum, following Equation 2. This approach ensures that niche breadth reflects the true spread of the environmental variable (here, salinity) for each vOTU across all available evidence rather than an average of two within-library estimates.

### Supplementary figures

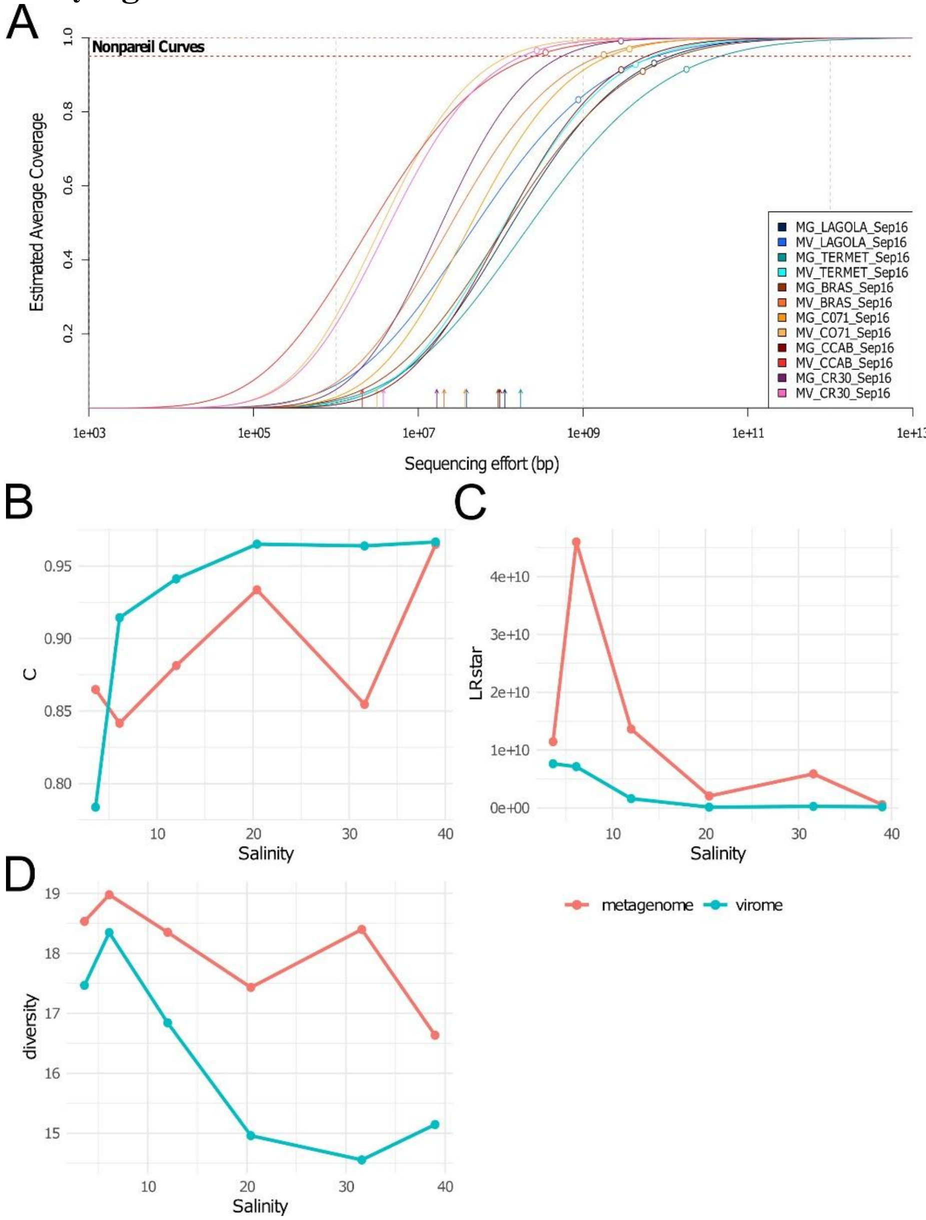


Figure S1. Sequencing coverage and diversity metrics across the salinity gradient. A) Non-pareil curves for metagenomes and viromes from the six sampling sites. B–D) Relationships between non-pareil statistics and salinity: B) coverage (C stat), C) estimated sequencing effort required (LRstar), and D) non-pareil diversity index, for both metagenomes and viromes.

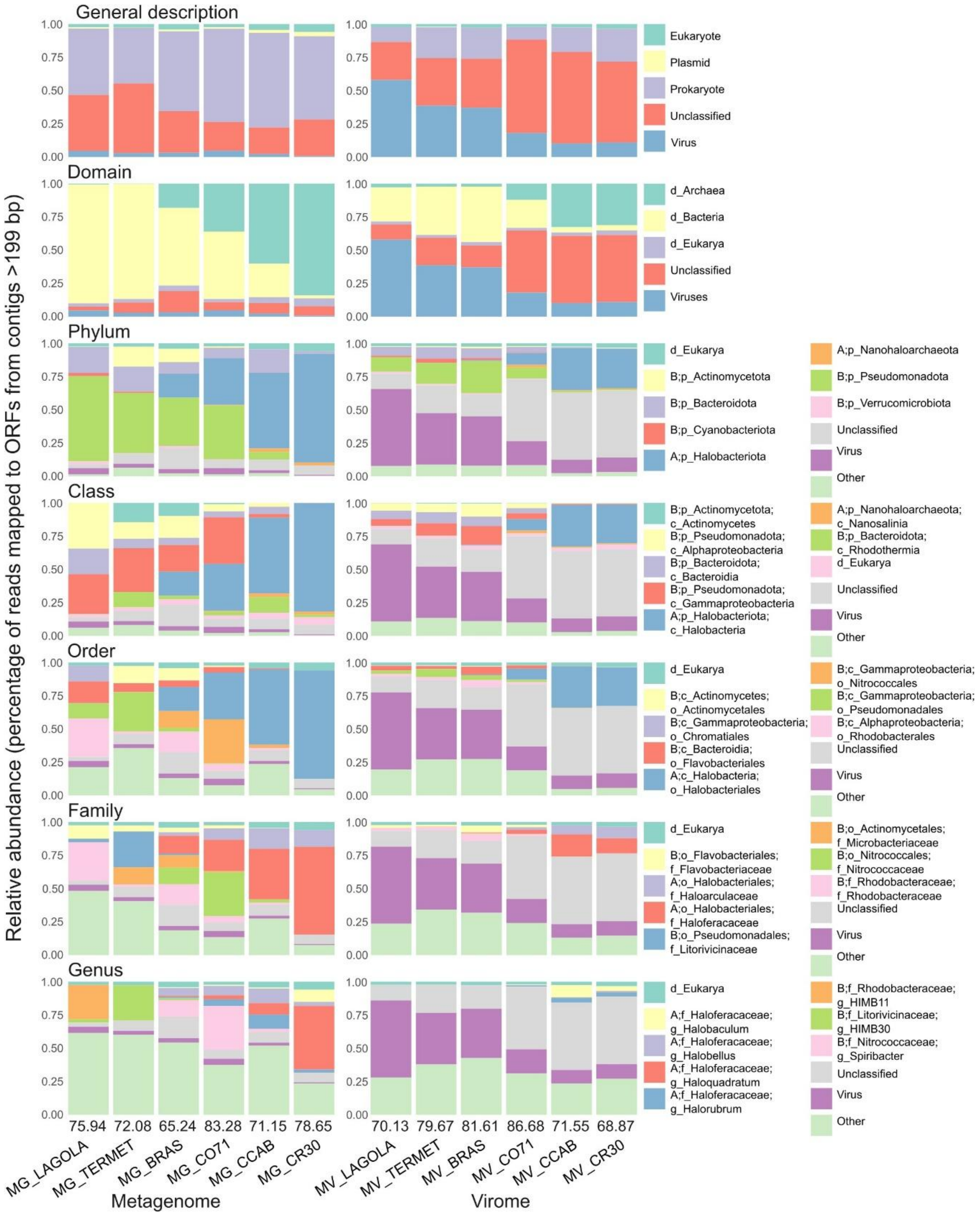


Figure S2. Relative abundances of mapped metagenomic reads across CDS classifications. Read mapping was performed with coverM (≥95% identity, ≥50% read coverage). CDSs from prokaryotic contigs were taxonomically annotated using refineM against GTDB-R220 representative genomes. Percentages above sample names indicate total mapped reads. The ten most abundant classifications are displayed; remaining classifications are grouped as "Other."

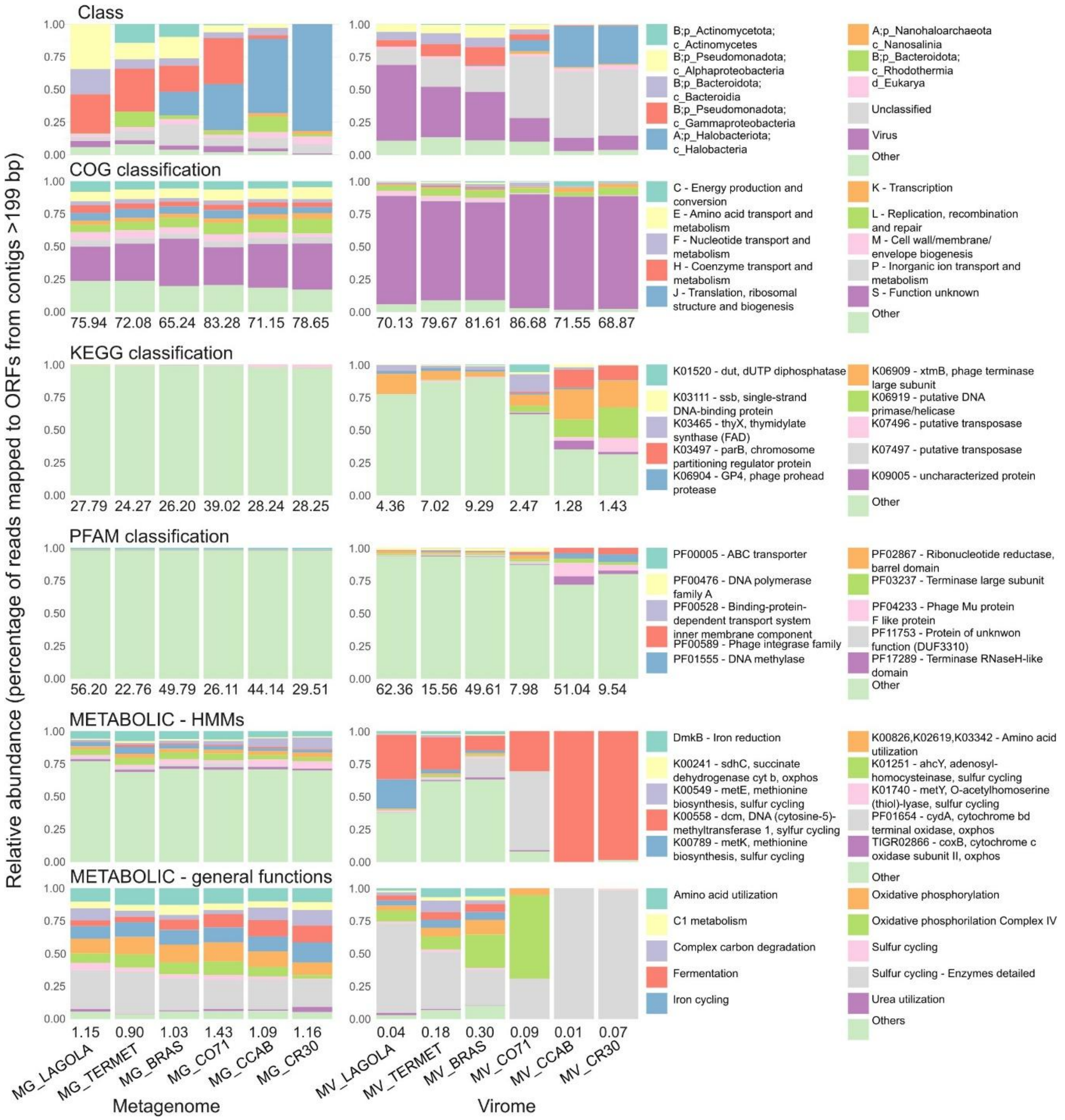


Figure S3. Relative abundances of predicted CDSs annotated with functional and metabolic databases. CDS annotations were performed at class level using refineM against GTDB-R220. COG classifications were assigned with eggnog-mapper. Functional annotations were obtained from DRAM (KEGG and PFAM databases) and METABOLIC (higher-level classifications and individual HMMs). Percentages below each bar indicate relative abundance of classified ORFs over total clean reads. The ten most abundant classifications are displayed; remaining classifications are grouped as "Other."

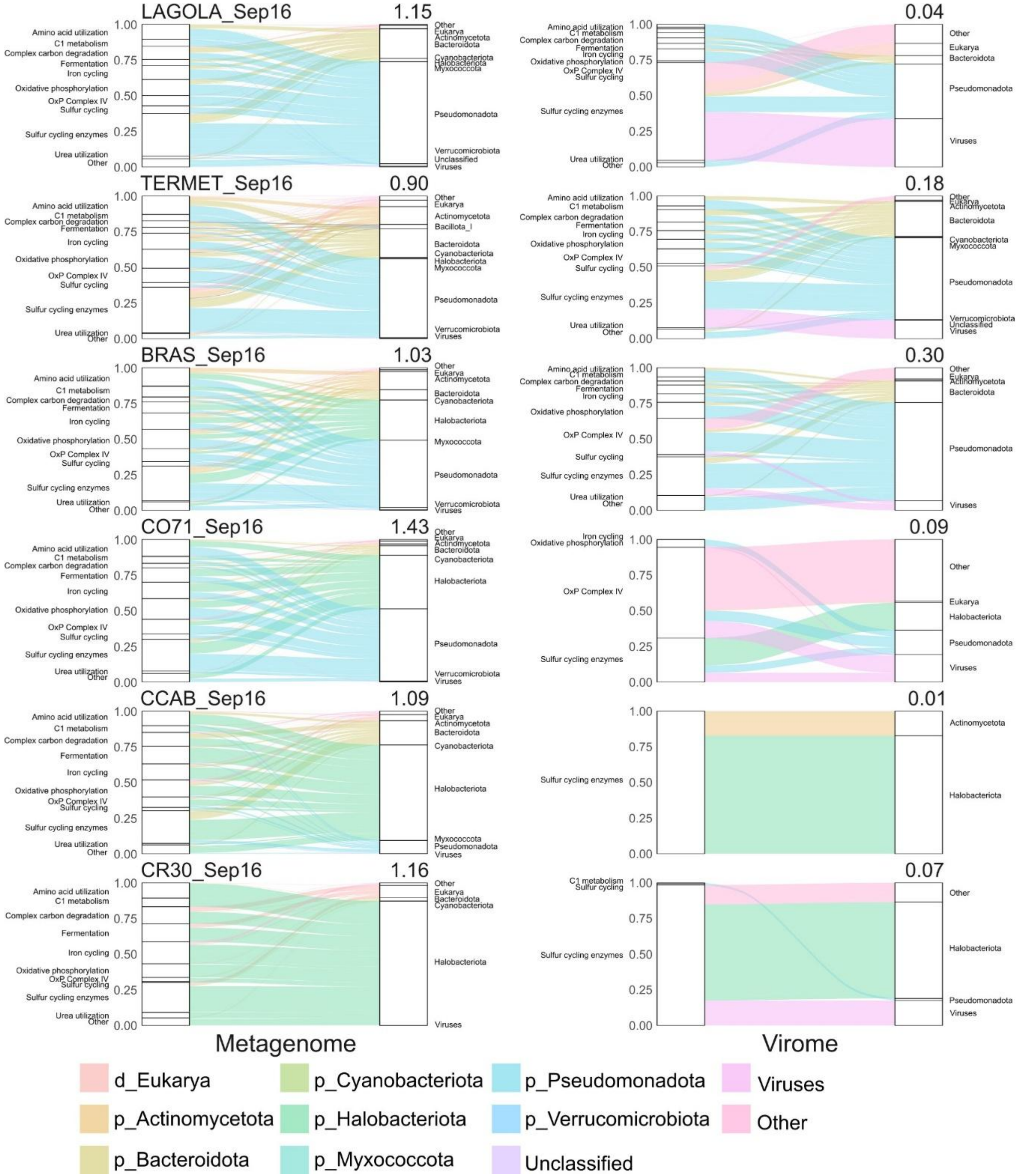


Figure S4. Relative abundances of metabolic functions across taxonomic groups along the salinity gradient. CDS annotations were assigned taxonomically using refineM against GTDB-R220 and functionally classified with METABOLIC at general function levels. Alluvial plots display the flow of relative abundances across taxonomic and functional categories. Percentages in the upper right corner of each panel indicate the proportion of reads of the respective sample.

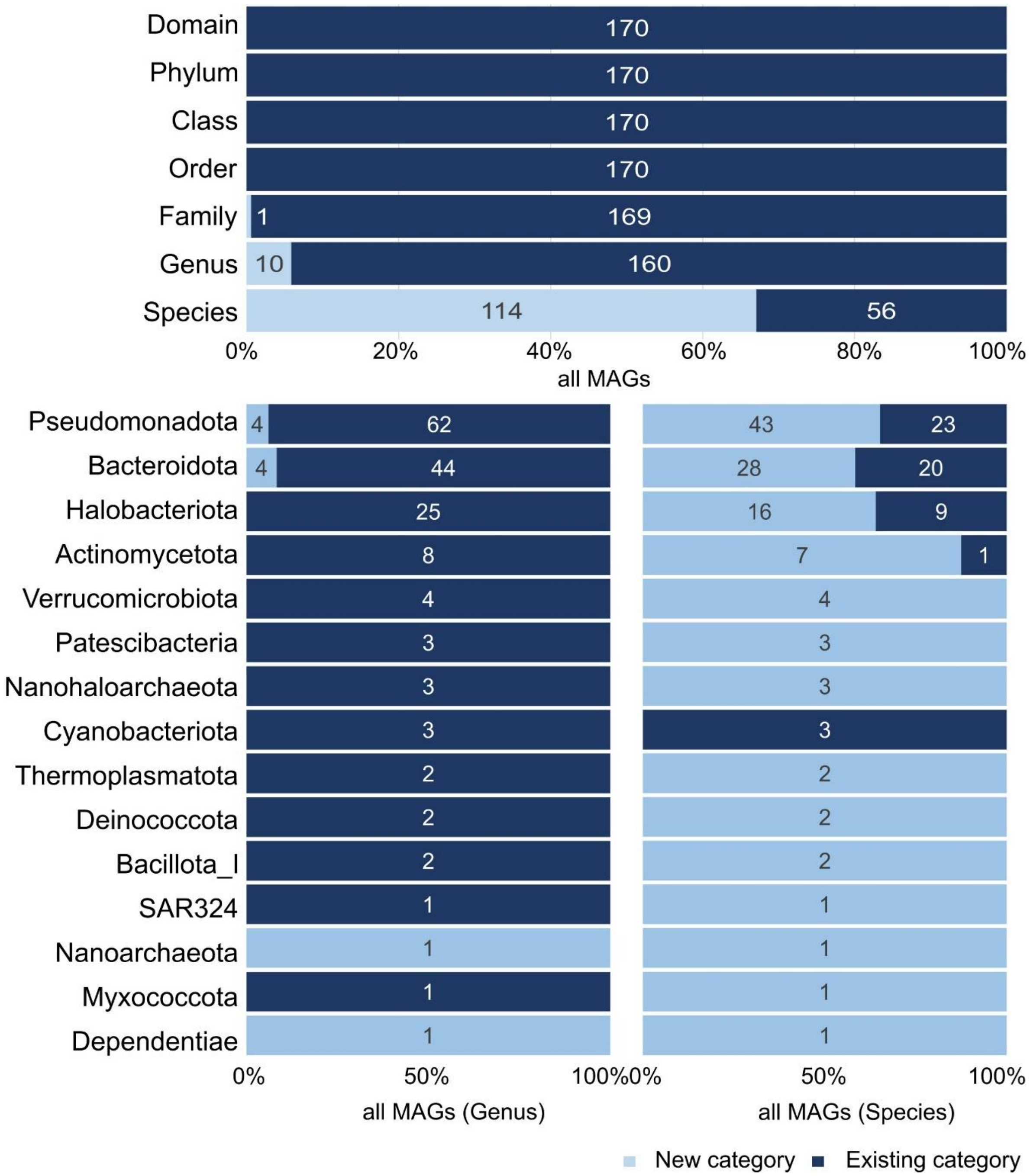


Figure S5. Taxonomic novelty of dereplicated MAGs. MAGs (n=170, completeness >50%, contamination <10%) were taxonomically classified using gtdb-tk with GTDB-R220. Upper panel shows the proportion of novel MAGs (without a representative genome associated in the database) across all taxonomic levels. Bottom panel displays the number and proportion of MAGs classified and unclassified at genus and species levels for each phylum.

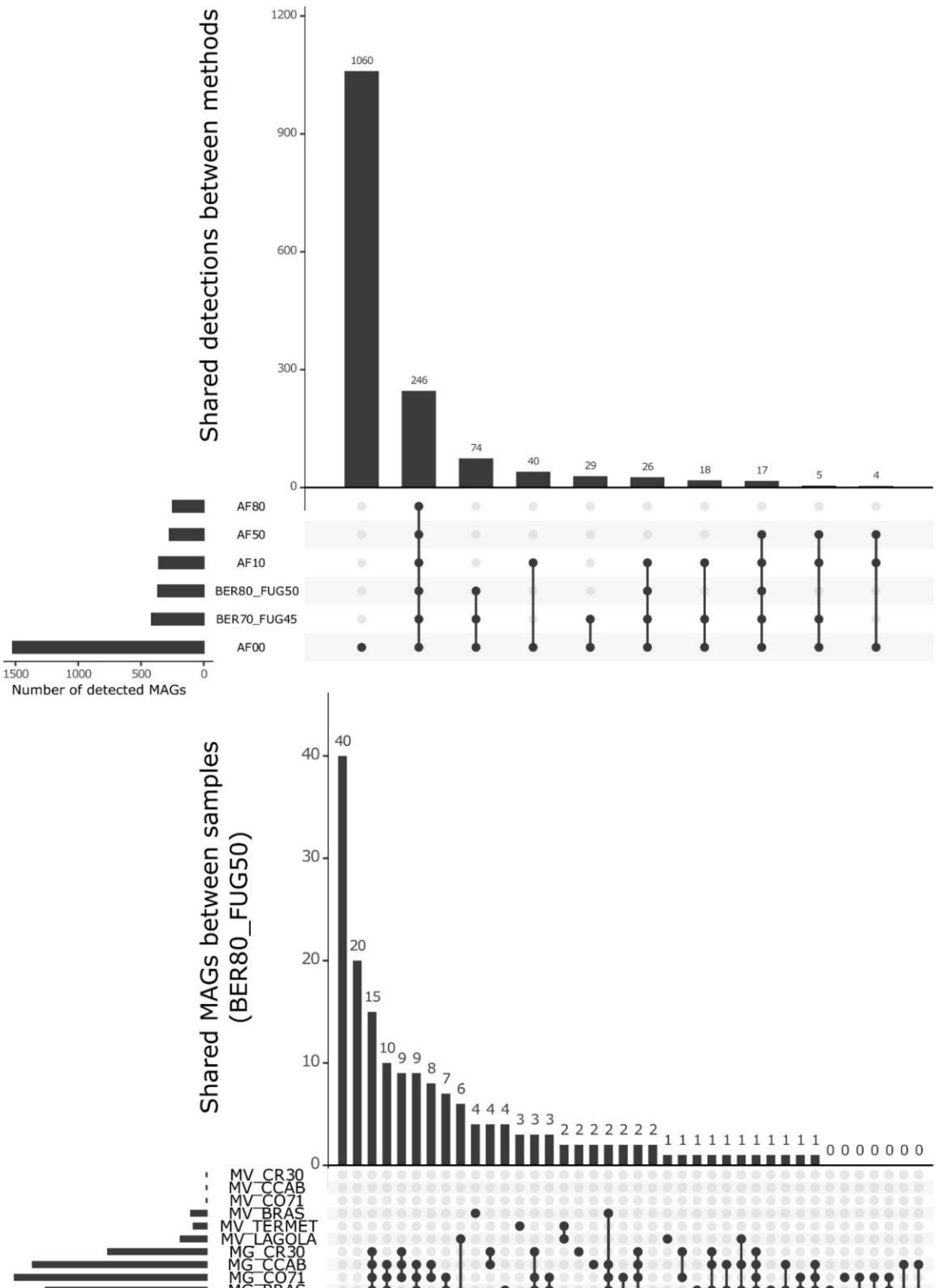


Figure S6. Consistency of MAG detection methods and sample-wise MAG distributions. Top) UpSet plot showing intersection of MAG presence-absence calls across six detection methods applied to all 12 samples, showing high consistency among approaches. Methods include: BER80_FUG50 (metapresence default), BER70_FUG45 (metapresence stringent), and aligned fraction thresholds of 0%, 10%, 50%, and 80% (AF00, AF10, AF50, AF80). Bottom) UpSet plot displaying the intersection of MAGs present across sample combinations using the metapresence default method (BER80_FUG50), revealing MAG distribution patterns along the salinity gradient.

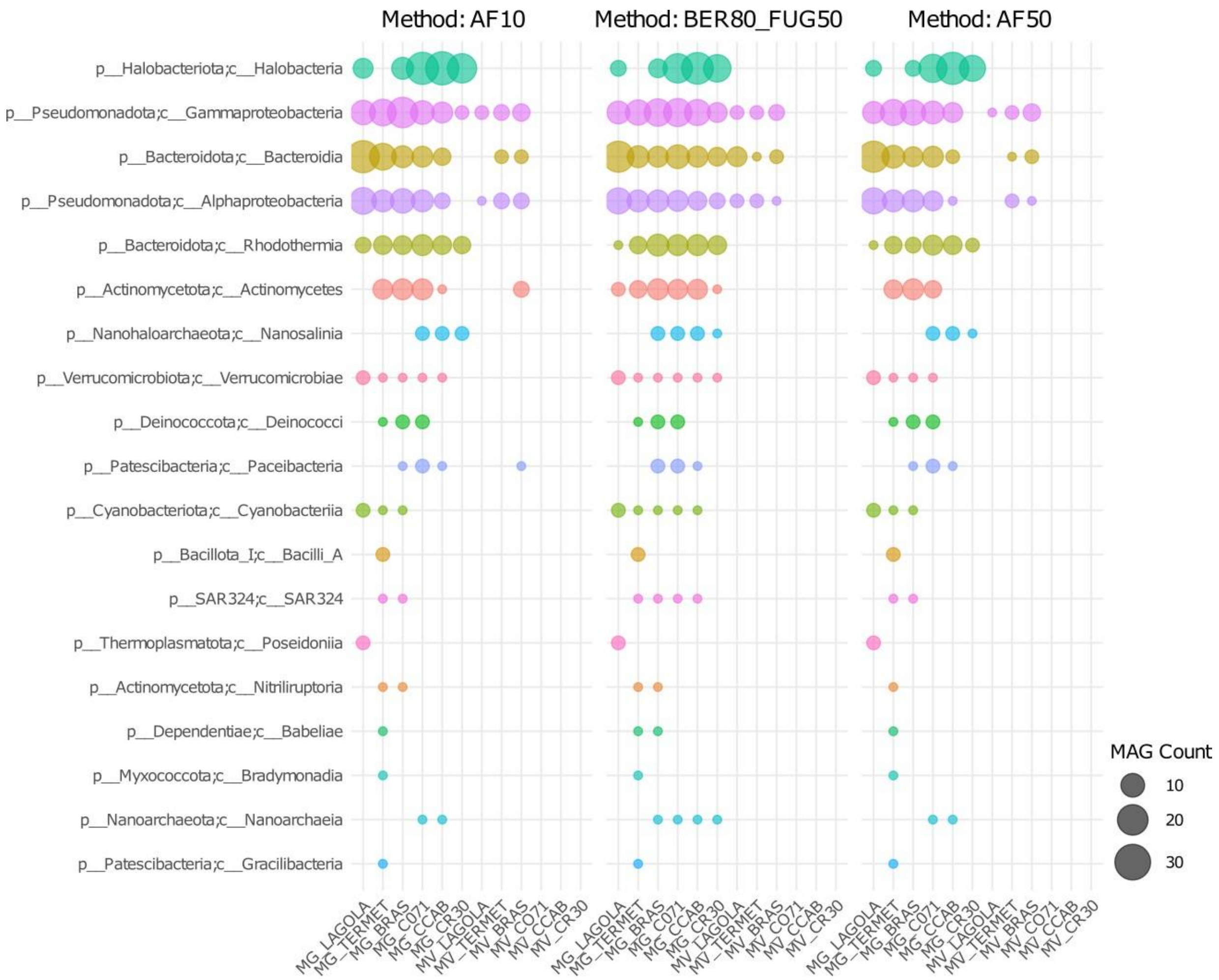


Figure S7. MAG detection across samples by taxonomic class using three methods. Bubble plots show the number of MAGs (circle size) detected in each sample according to their GTDB-R220 class-level taxonomy using three detection methods: aligned fraction threshold of 10% (AF10) and 50 % (AF50); and metapresence with default parameters (BER80_FUG50).

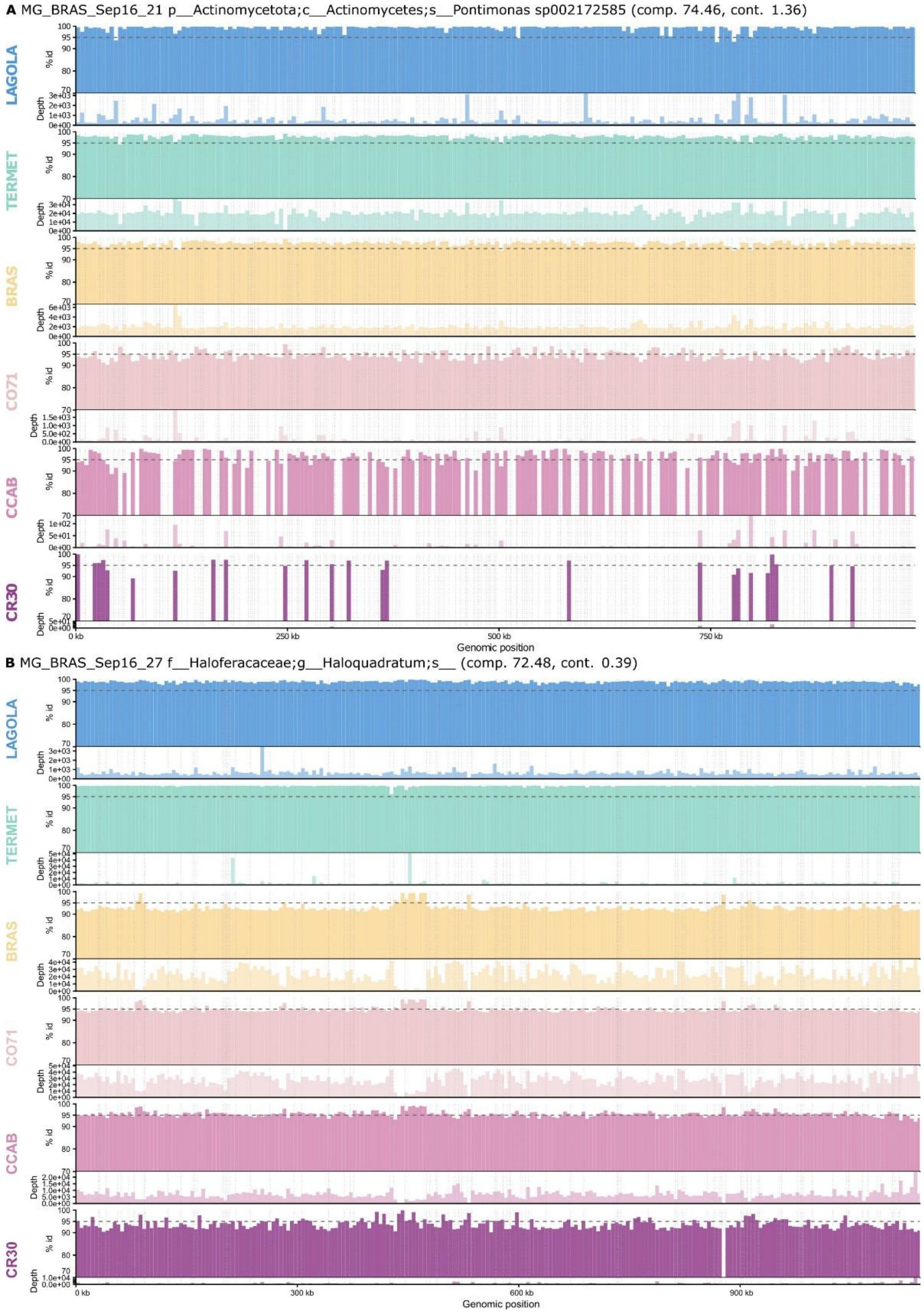


Figure S8. Fragment recruitment plots of the two cosmopolitan metagenome-assembled genomes across the salinity gradient. Trimmed metagenomic reads from each of the six sampling sites were mapped against the linearized genome sequence of (A) MAG MG_BRAS_Sep16_21, affiliated to *Pontimonas* sp002172585, and (B) MAG MG_BRAS_Sep16_27, affiliated with an unclassified *Haloquadratum* genomospecies. For each site, the upper track shows the mean nucleotide identity (%) of mapped reads per 5 kb genomic window, with a dashed line indicating the 95% identity threshold. The lower track shows the corresponding read depth per window. Dotted vertical lines indicate contig boundaries within each MAG. Only reads mapping with ≥70% nucleotide identity were retained. Sites are ordered along the increasing salinity gradient from top to bottom.

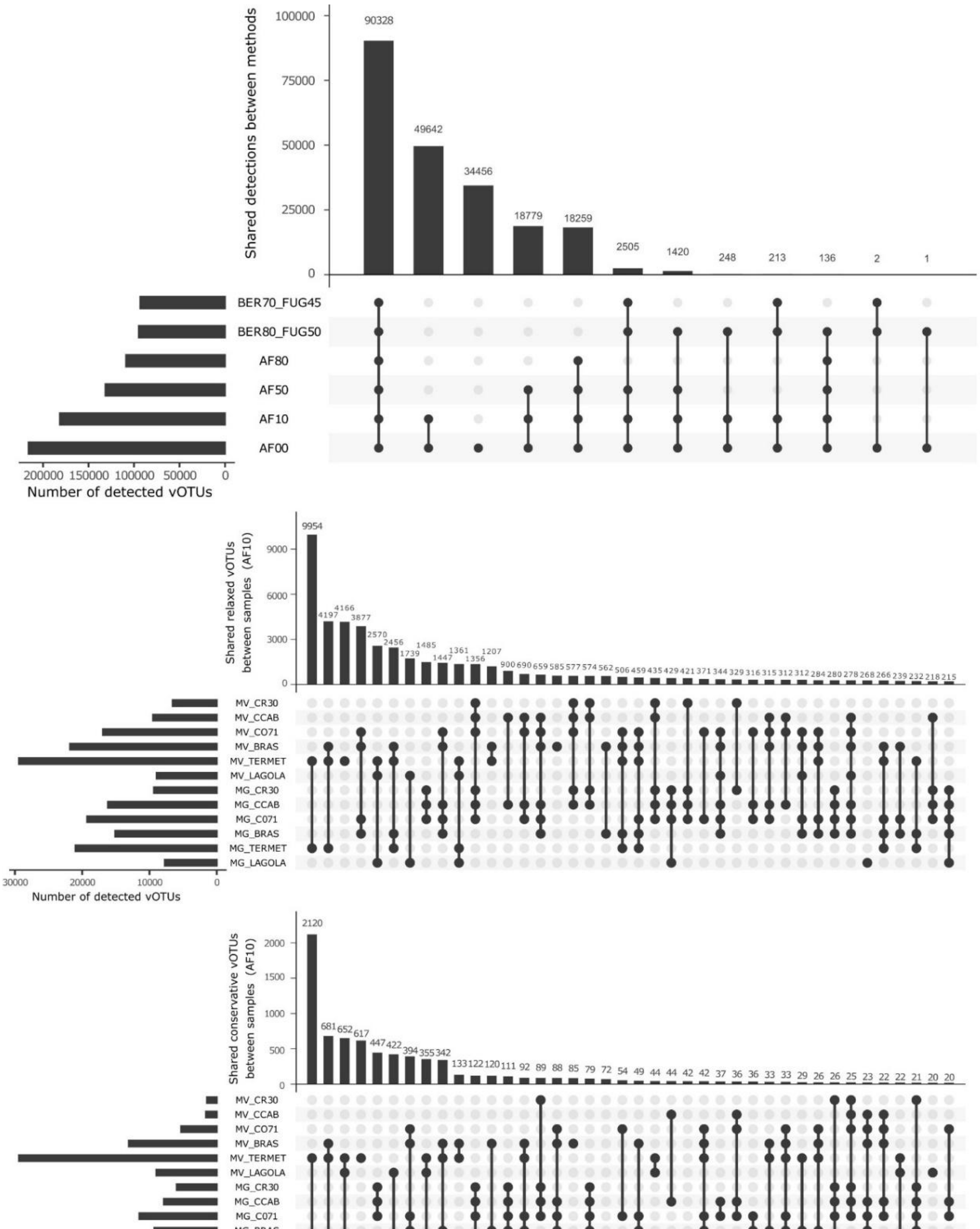


Figure S9. Consistency of vOTU detection methods and sample-wise vOTU distributions. Top) UpSet plot showing intersection of vOTU presence-absence calls across six detection methods applied to all 12 samples. Methods include: BER80_FUG50 (metapresence default), BER70_FUG45 (metapresence stringent), and aligned fraction thresholds of 0%, 10%, 50%, and 80% (AF00, AF10, AF50, AF80). Middle) UpSet plot displaying the intersection of relaxed vOTUs present across sample combinations using the aligned fraction 50% threshold (AF50), revealing vOTU distribution patterns along the salinity gradient. Bottom) UpSet plot displaying the intersection of conservative vOTUs present across sample combinations using the aligned fraction 50% threshold (AF50), showing more stringent vOTU occurrence patterns.

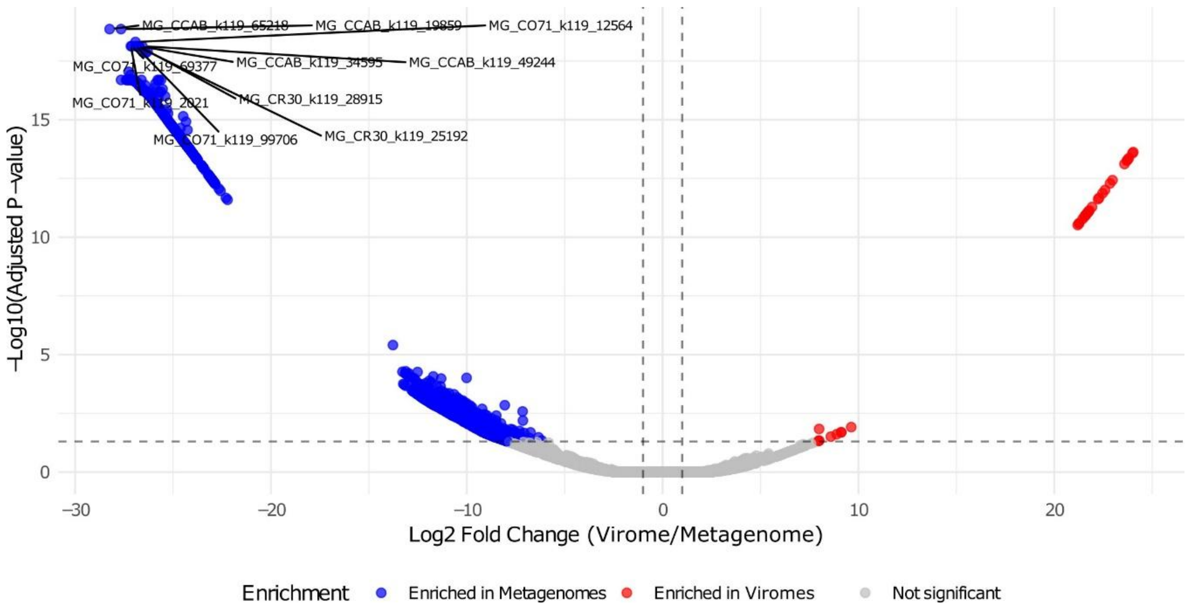


Figure S10. Differential abundance of vOTUs between metagenomes and viromes. Volcano plot showing enrichment of vOTUs in viromes relative to metagenomes (x-axis: $\log_2$ fold-change virome/metagenome; y-axis: $-\log_{10}$ adjusted p-value). vOTU abundances were quantified as TPM values and compared using DESeq2. Points above the significance threshold indicate differentially abundant vOTUs between the two sequence types.

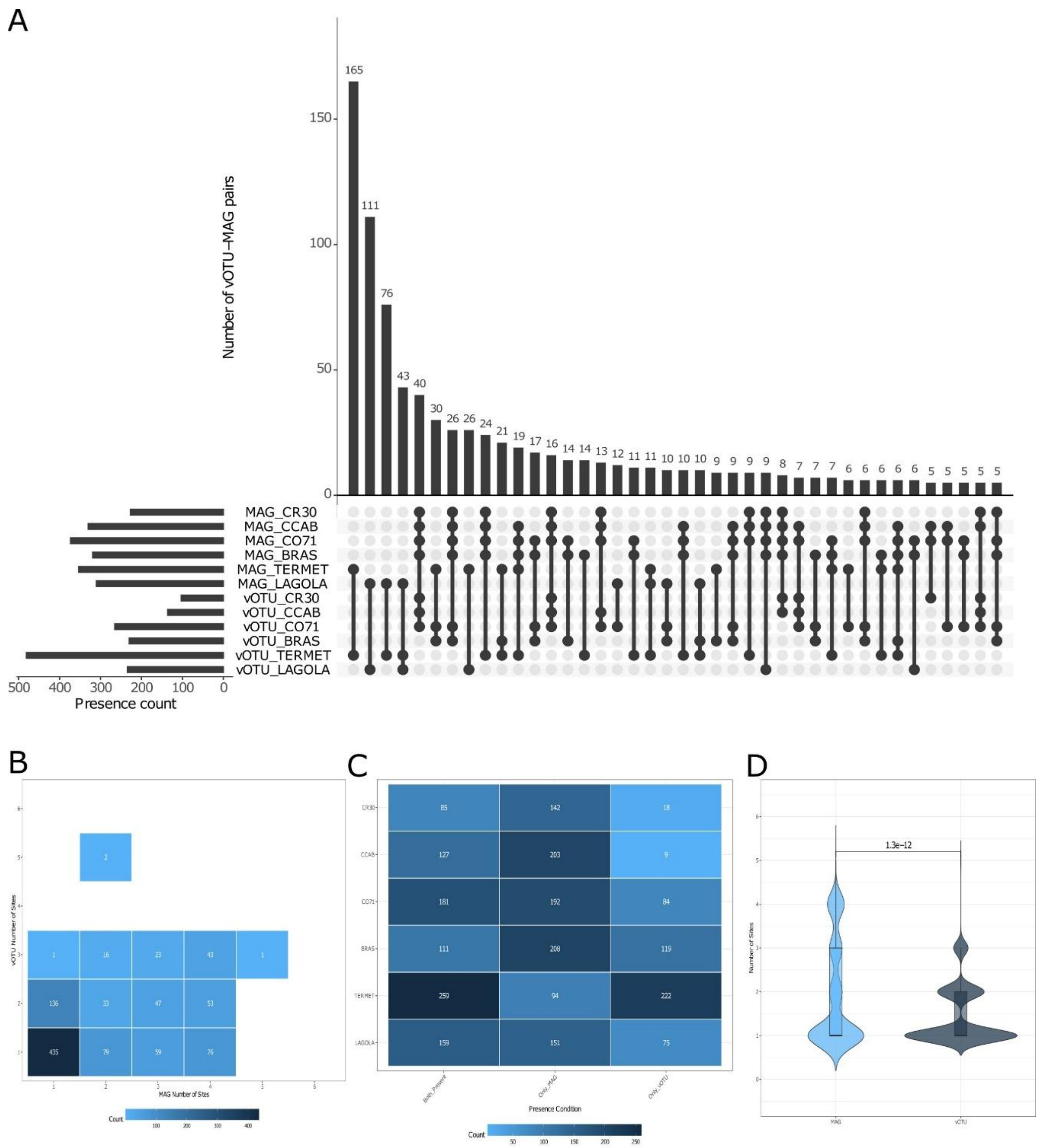


Figure S11. Virus-host pair distribution and co-occurrence patterns (conservative vOTUs, PHIST ≥3 shared k-mers; n = 1004). Virus-host pairs were predicted using PHIST (≥3 shared k-mers); MAG presence determined with metapresence and vOTU detection with ≥50% aligned fraction. A) UpSet plot showing MAG and vOTU per-site abundance (left histogram), site distribution of virus-host pairs (center), and interaction frequency (top histogram). B) Number of sites where each virus-host pair member is detected. C) Co-occurrence frequency within sites: whether pair members are predominantly found together or detected separately. D) Detection frequency of pair members (Wilcoxon paired t-test, p-value shown).

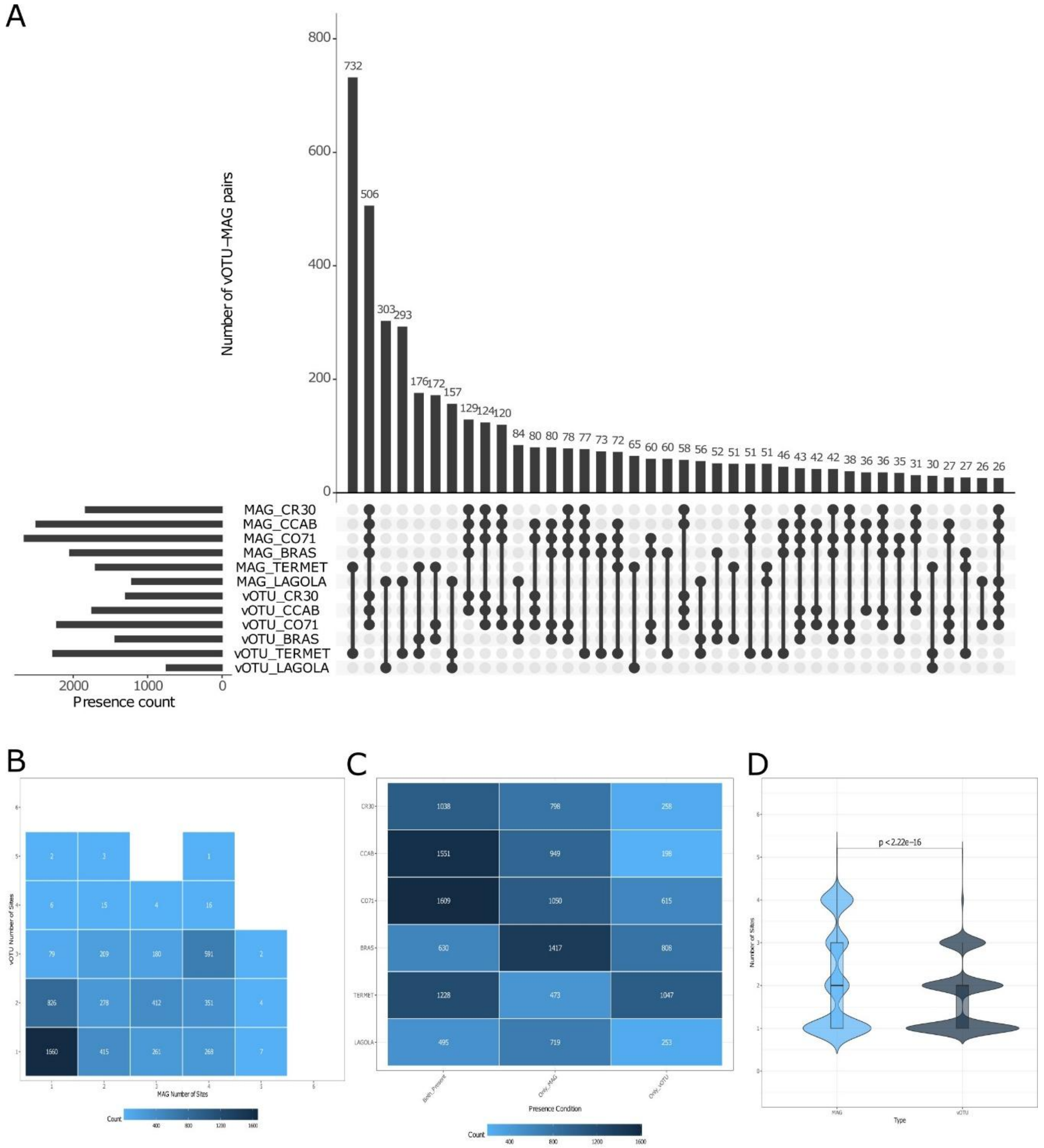


Figure S12. Virus-host pair distribution and co-occurrence patterns (relaxed vOTUs, all PHIST; n = 5590). Virus-host pairs were predicted using PHIST; MAG presence determined with metapresence and vOTU detection with ≥50% aligned fraction. A) UpSet plot showing MAG and vOTU per-site abundance (left histogram), site distribution of virus-host pairs (center), and interaction frequency (top histogram). B) Number of sites where each virus-host pair member is detected. C) Co-occurrence frequency within sites: whether pair members are predominantly found together or detected separately. D) Detection frequency of pair members (Wilcoxon paired t-test, p-value shown).

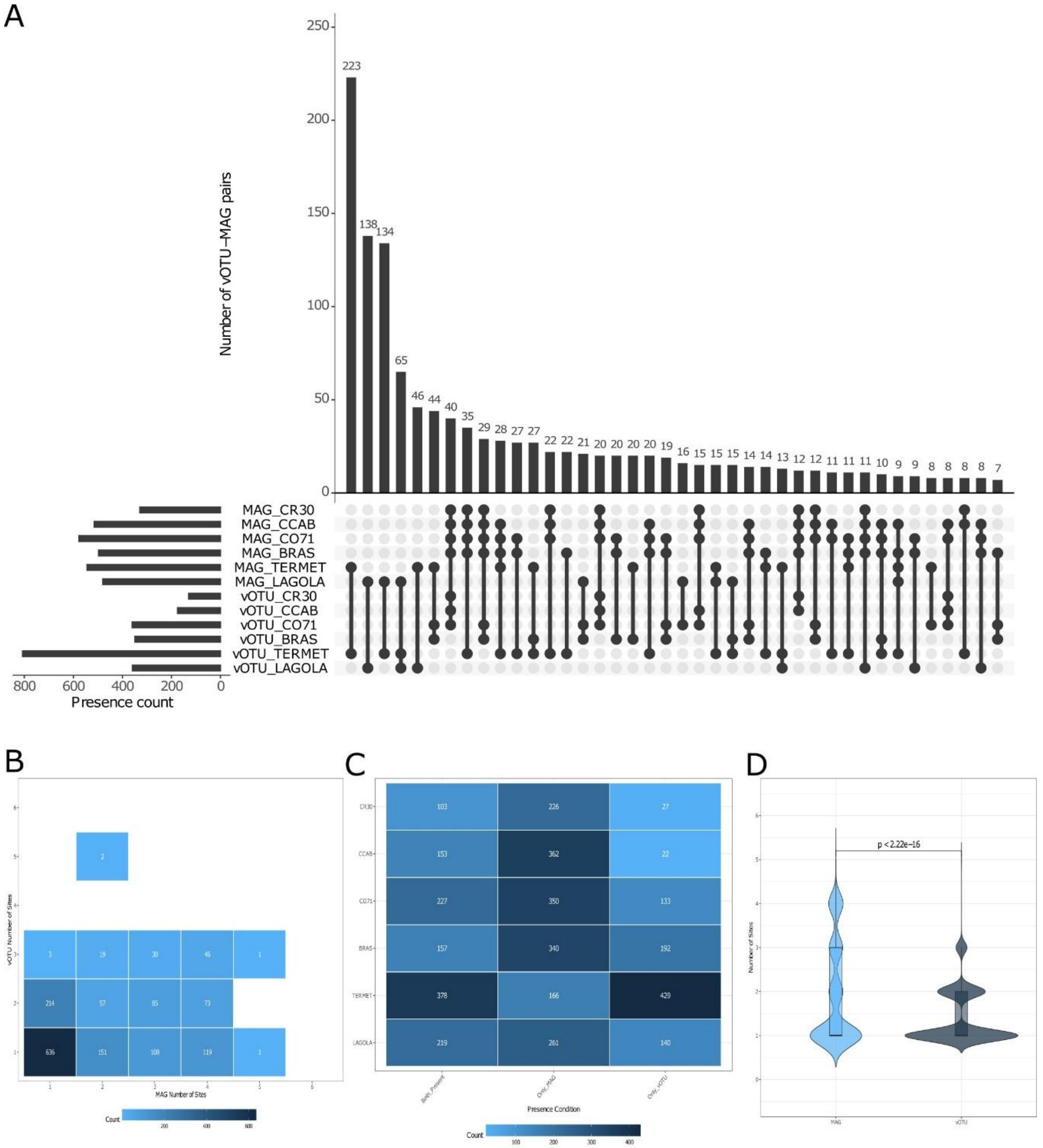


Figure S13. Virus-host pair distribution and co-occurrence patterns (conservative vOTUs, all PHIST; n = 1545). Virus-host pairs were predicted using PHIST; MAG presence determined with metapresence and vOTU detection with ≥50% aligned fraction. A) UpSet plot showing MAG and vOTU per-site abundance (left histogram), site distribution of virus-host pairs (center), and interaction frequency (top histogram). B) Number of sites where each virus-host pair member is detected. C) Co-occurrence frequency within sites: whether pair members are predominantly found together or detected separately. D) Detection frequency of pair members (Wilcoxon paired t-test, p-value shown).

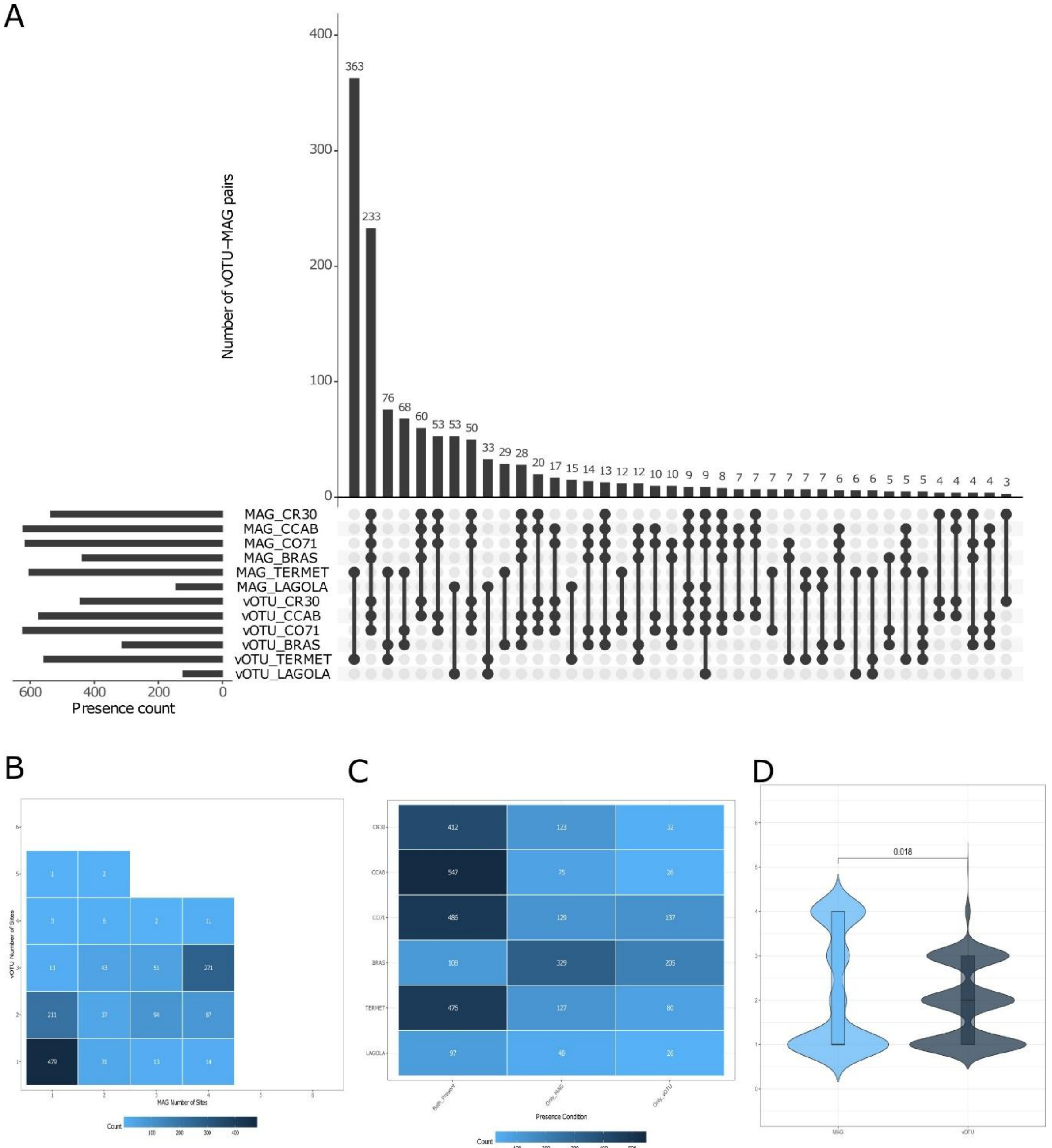


Figure S14. Virus-host pair distribution and co-occurrence patterns (relaxed vOTUs, PHIST-WiSH; n = 1369). Virus-host pairs were predicted using PHIST-WiSH; MAG presence determined with metapresence and vOTU detection with ≥50% aligned fraction. A) UpSet plot showing MAG and vOTU per-site abundance (left histogram), site distribution of virus-host pairs (center), and interaction frequency (top histogram). B) Number of sites where each virus-host pair member is detected. C) Co-occurrence frequency within sites: whether pair members are predominantly found together or detected separately. D) Detection frequency of pair members (Wilcoxon paired t-test, p-value shown).

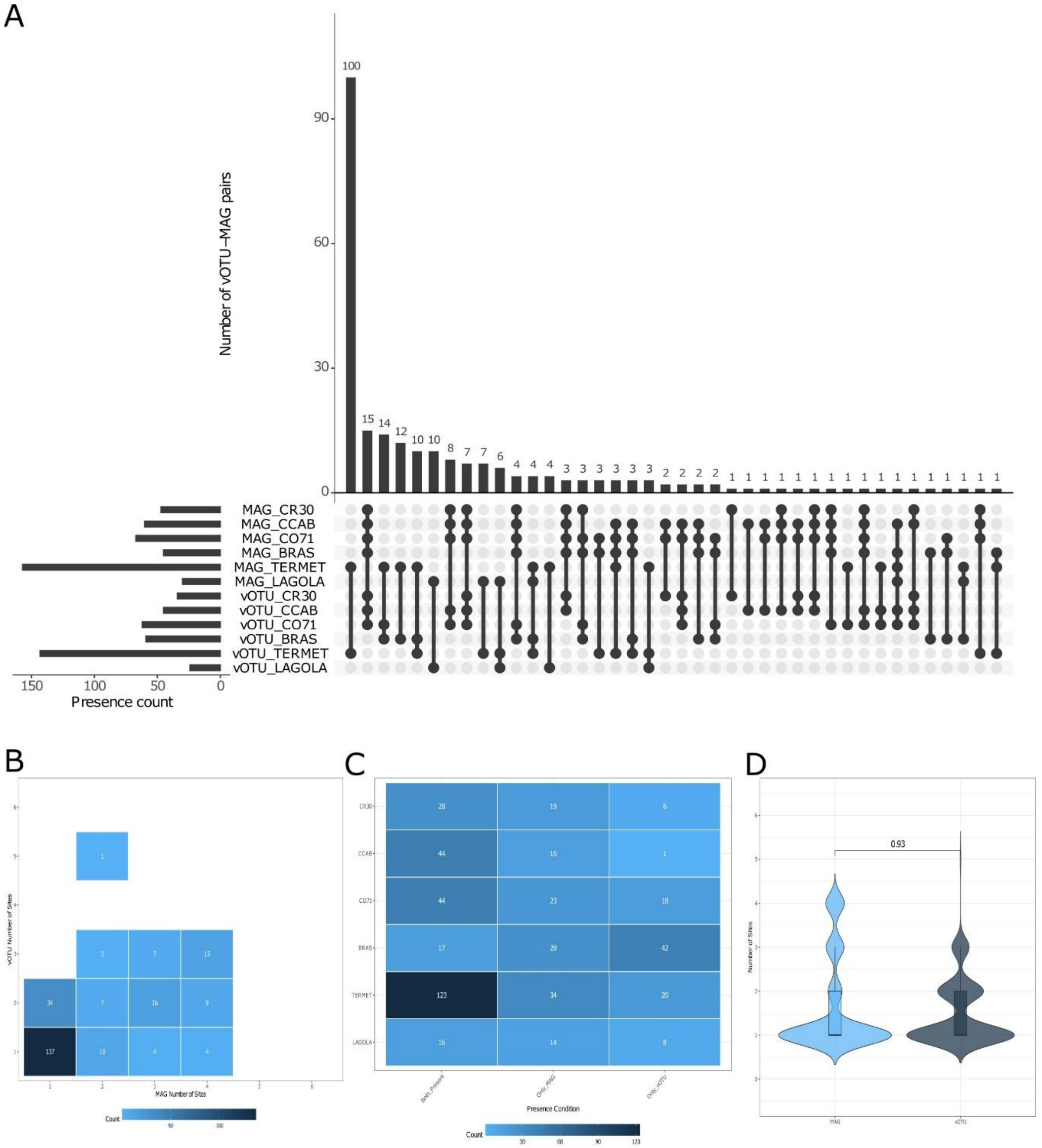


Figure S15. Virus-host pair distribution and co-occurrence patterns (conservative vOTUs, PHIST-WiSH; n = 247). Virus-host pairs were predicted using PHIST-WiSH; MAG presence determined with metapresence and vOTU detection with ≥50% aligned fraction. A) UpSet plot showing MAG and vOTU per-site abundance (left histogram), site distribution of virus-host pairs (center), and interaction frequency (top histogram). B) Number of sites where each virus-host pair member is detected. C) Co-occurrence frequency within sites: whether pair members are predominantly found together or detected separately. D) Detection frequency of pair members (Wilcoxon paired t-test, p-value shown).

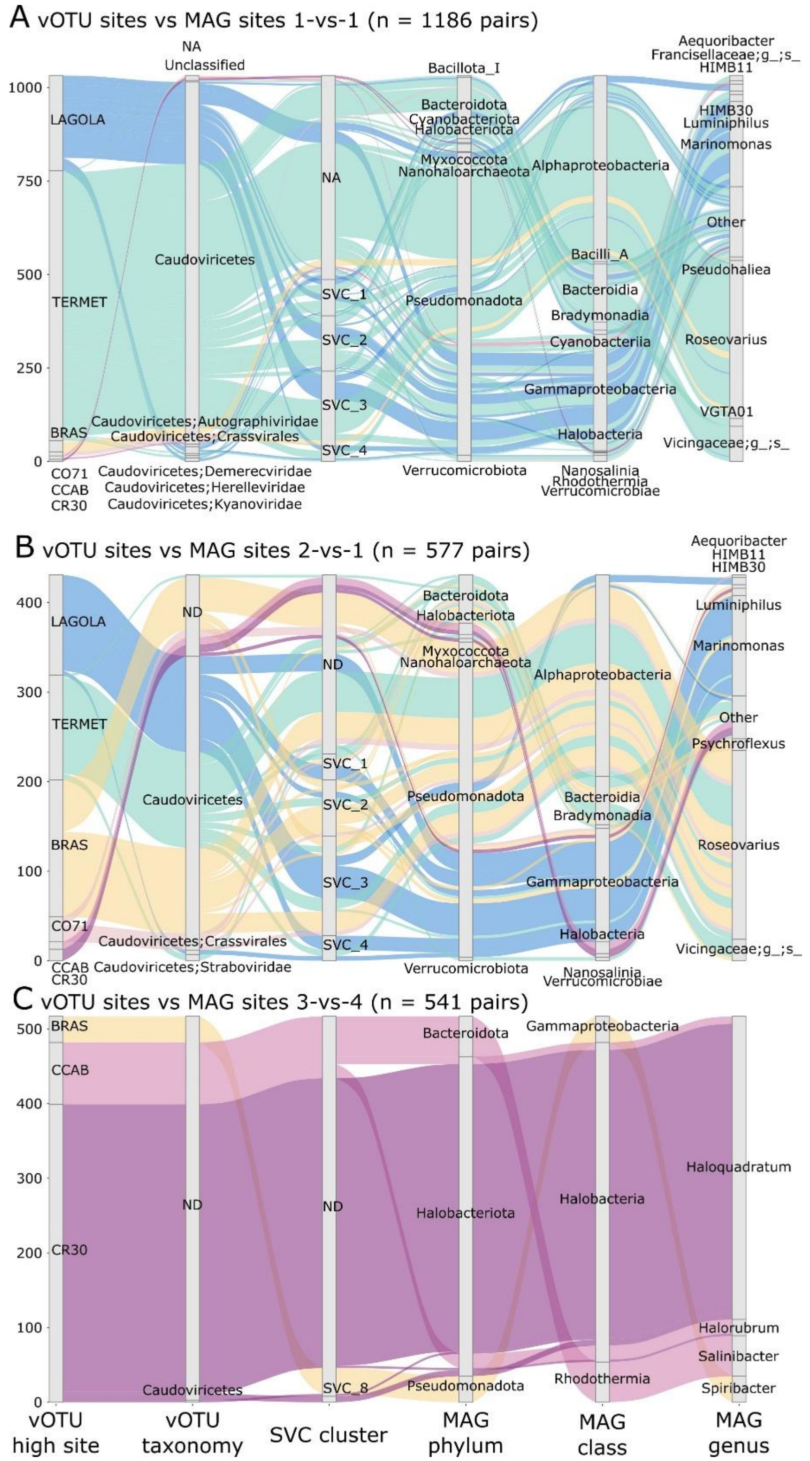


Figure S16. Alluvial diagrams of predicted virus-host pairs for three predominant co-occurrence patterns. Flows connect the site of highest vOTU abundance, vOTU taxonomic classification, Super Viral Cluster (SVC) affiliation (from Figure 5), and predicted host MAG taxonomy at phylum, class, and genus level, colored by the site of highest vOTU abundance following the color scheme used throughout the manuscript. (A) Pairs restricted to a single site for both entities (1-vs-1). (B) Pairs where the vOTU spanned two sites while the host MAG was detected at only one (2-vs-1). (C) Pairs where the vOTU was detected at three sites and its host MAG at four (3-vs-4). Virus-host predictions are based on PHIST with a minimum of three shared k-mers and relative to Figure 5.

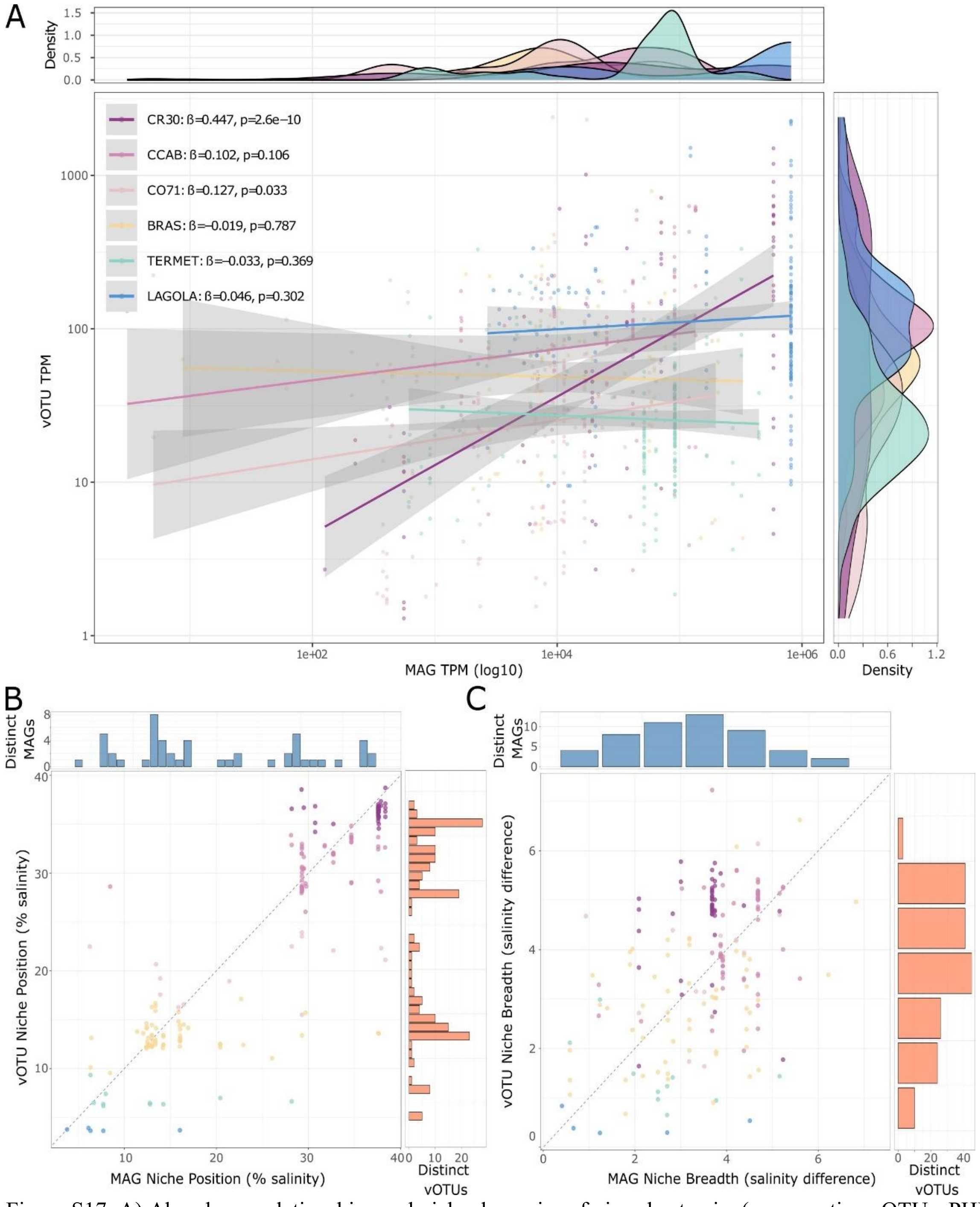


Figure S17. A) Abundance relationships and niche dynamics of virus-host pairs (conservative vOTUs, PHIST ≥3 shared k-mers). Linear regression of vOTU and MAG abundances (TPM, $\log_{10}$-transformed) for pairs at each site (n total= 849). Virus-host pairs predicted with PHIST (≥3 shared k-mers) were quantified by read mapping with coverM (≥95% identity, ≥50% read coverage). Pairs with unmapped members in a given sample are excluded. Linear model statistics for each site include: regression slope (β), and p-value. B-C) Niche position and breadth analyses for n = 191 vOTU-MAGs pairs. B) Niche position of virus-host pairs calculated as the salinity optimum using ecospat. C) Niche breadth expressed as the salinity range surrounding niche position. Lateral histograms display the number of distinct MAGs and vOTUs across 1% salinity intervals. Dot colors indicate the sample where each vOTU had highest abundance.

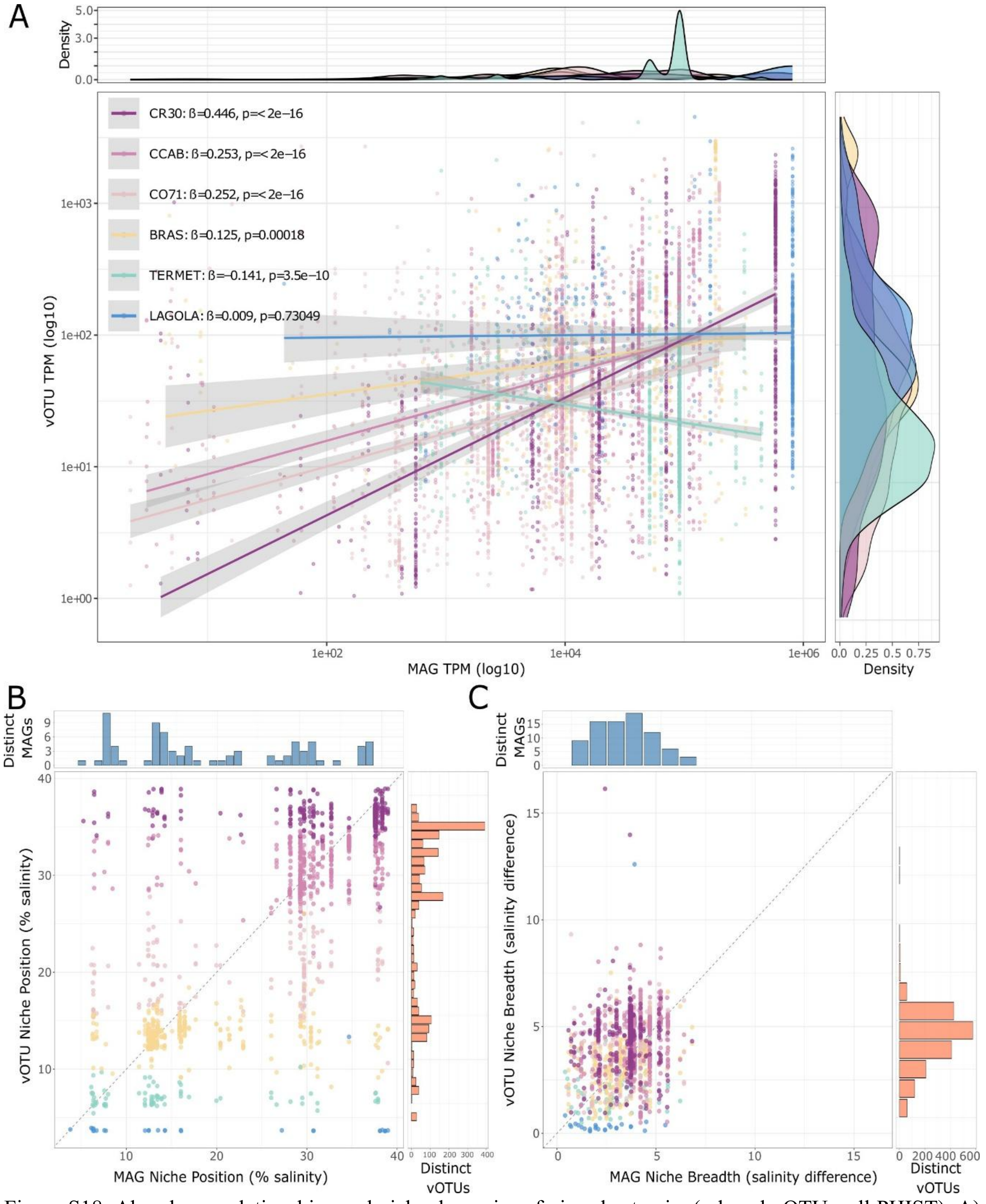


Figure S18. Abundance relationships and niche dynamics of virus-host pairs (relaxed vOTUs, all PHIST). A) Linear regression of vOTU and MAG abundances (TPM, $log_{10}$-transformed) for pairs at each site (n total= 6076). Virus-host pairs predicted with PHIST were quantified by read mapping with coverM (≥95% identity, ≥50% read coverage). Pairs with unmapped members in a given sample are excluded. Linear model statistics for each site include: regression slope (β), and p-value. B-C) Niche position and breadth analyses for n = 1875 vOTU-MAGs pairs. B) Niche position of virus-host pairs calculated as the salinity optimum using ecospat. C) Niche breadth expressed as the salinity range surrounding niche position. Lateral histograms display the number of distinct MAGs and vOTUs across 1% salinity intervals. Dot colors indicate the sample where each vOTU had highest abundance.

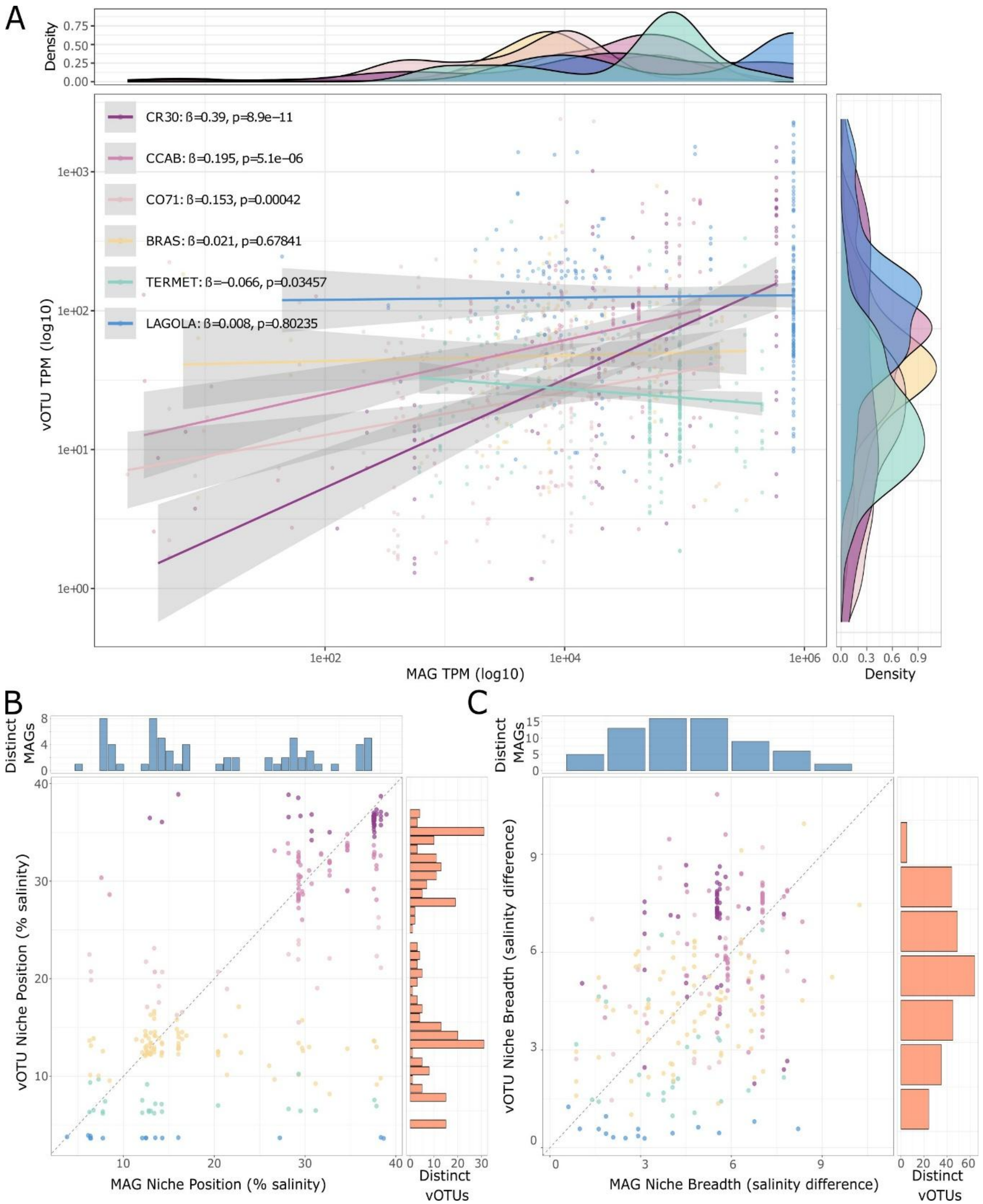


Figure S19. Abundance relationships and niche dynamics of virus-host pairs (conservative vOTUs, all PHIST). A) Linear regression of vOTU and MAG abundances (TPM, $\log_{10}$-transformed) for pairs at each site (n total= 1100). Virus-host pairs predicted with PHIST were quantified by read mapping with coverM (≥95% identity, ≥50% read coverage). Pairs with unmapped members in a given sample are excluded. Linear model statistics for each site include: number of pairs (n), regression slope (β), and p-value. B-C) Niche position and breadth analyses for n = 267 vOTU-MAGs pairs. B) Niche position of virus-host pairs calculated as the salinity optimum using ecospat. C) Niche breadth expressed as the salinity range surrounding niche position. Lateral histograms display the number of distinct MAGs and vOTUs across 1% salinity intervals. Dot colors indicate the sample where each vOTU had highest abundance.

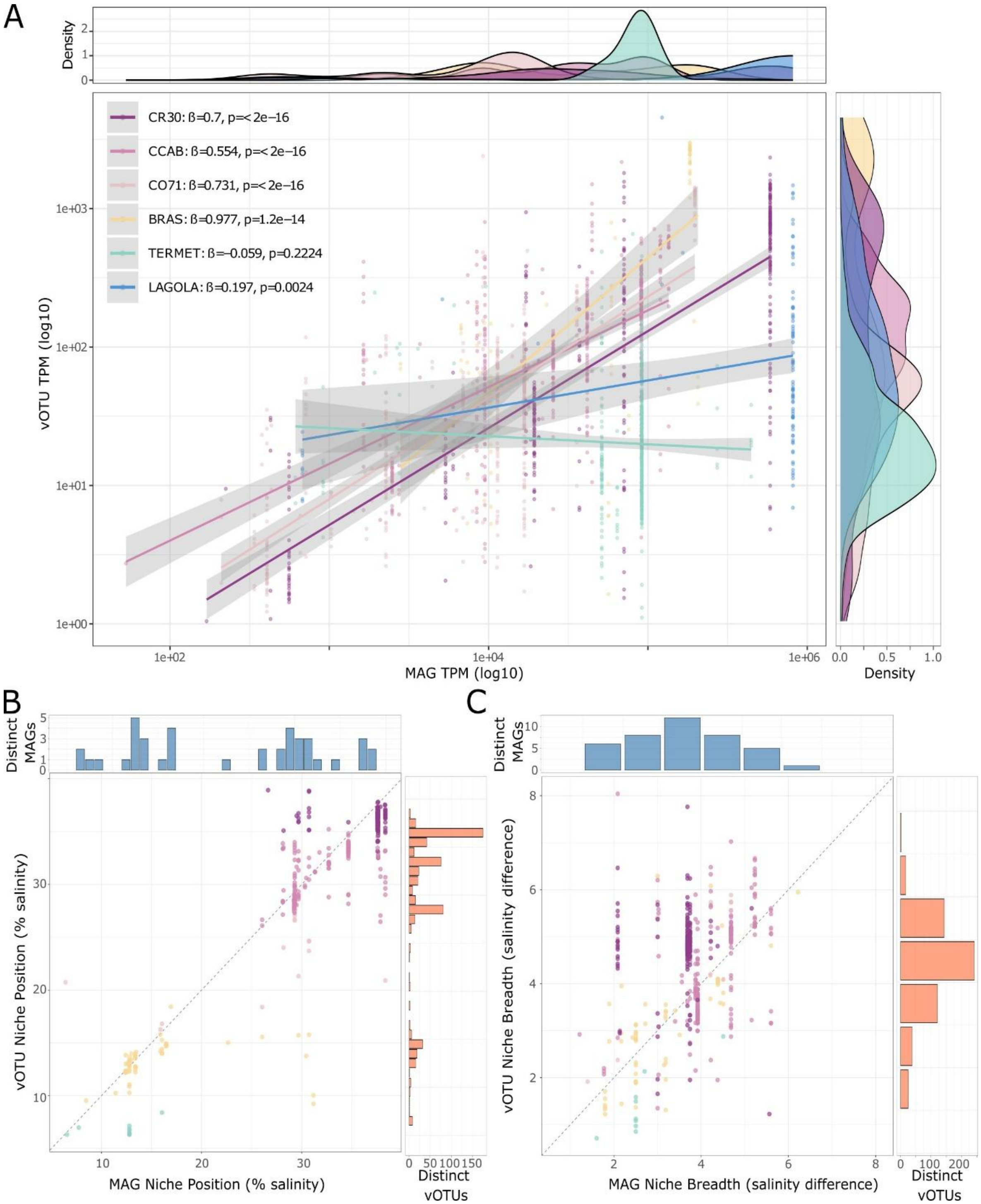


Figure S20. Abundance relationships and niche dynamics of virus-host pairs (relaxed vOTUs, PHIST-WiSH). A) Linear regression of vOTU and MAG abundances (TPM, $\log_{10}$-transformed) for pairs at each site (n total= 2088). Virus-host pairs predicted with PHIST-WiSH were quantified by read mapping with coverM (≥95% identity, ≥50% read coverage). Pairs with unmapped members in a given sample are excluded. Linear model statistics for each site include: number of pairs (n), regression slope (β), and p-value. B-C) Niche position and breadth analyses for n = 592 vOTU-MAGs pairs. B) Niche position of virus-host pairs calculated as the salinity optimum using ecospat. C) Niche breadth expressed as the salinity range surrounding niche position. Lateral histograms display the number of distinct MAGs and vOTUs across 1% salinity intervals. Dot colors indicate the sample where each vOTU had highest abundance.

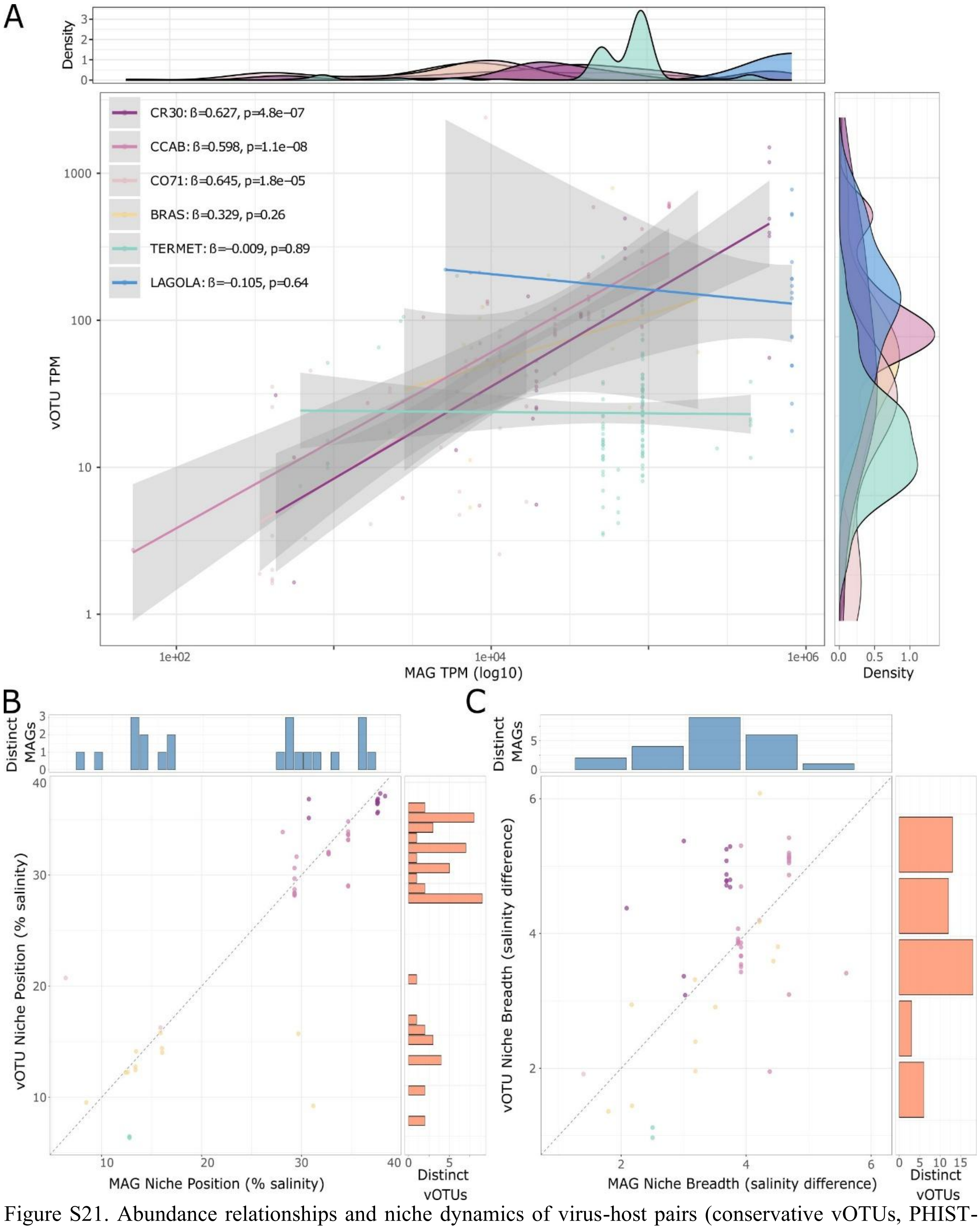


Figure S21. Abundance relationships and niche dynamics of virus-host pairs (conservative vOTUs, PHIST-WiSH). A) Linear regression of vOTU and MAG abundances (TPM, $\log_{10}$-transformed) for pairs at each site (n total= 264). Virus-host pairs predicted with PHIST-WiSH were quantified by read mapping with coverM (≥95% identity, ≥50% read coverage). Pairs with unmapped members in a given sample are excluded. Linear model statistics for each site include: number of pairs (n), regression slope (β), and p-value. B-C) Niche position and breadth analyses for n = 54 vOTU-MAGs pairs. B) Niche position of virus-host pairs calculated as the salinity optimum using ecospat. C) Niche breadth expressed as the salinity range surrounding niche position. Lateral histograms display the number of distinct MAGs and vOTUs across 1% salinity intervals. Dot colors indicate the sample where each vOTU had highest abundance.

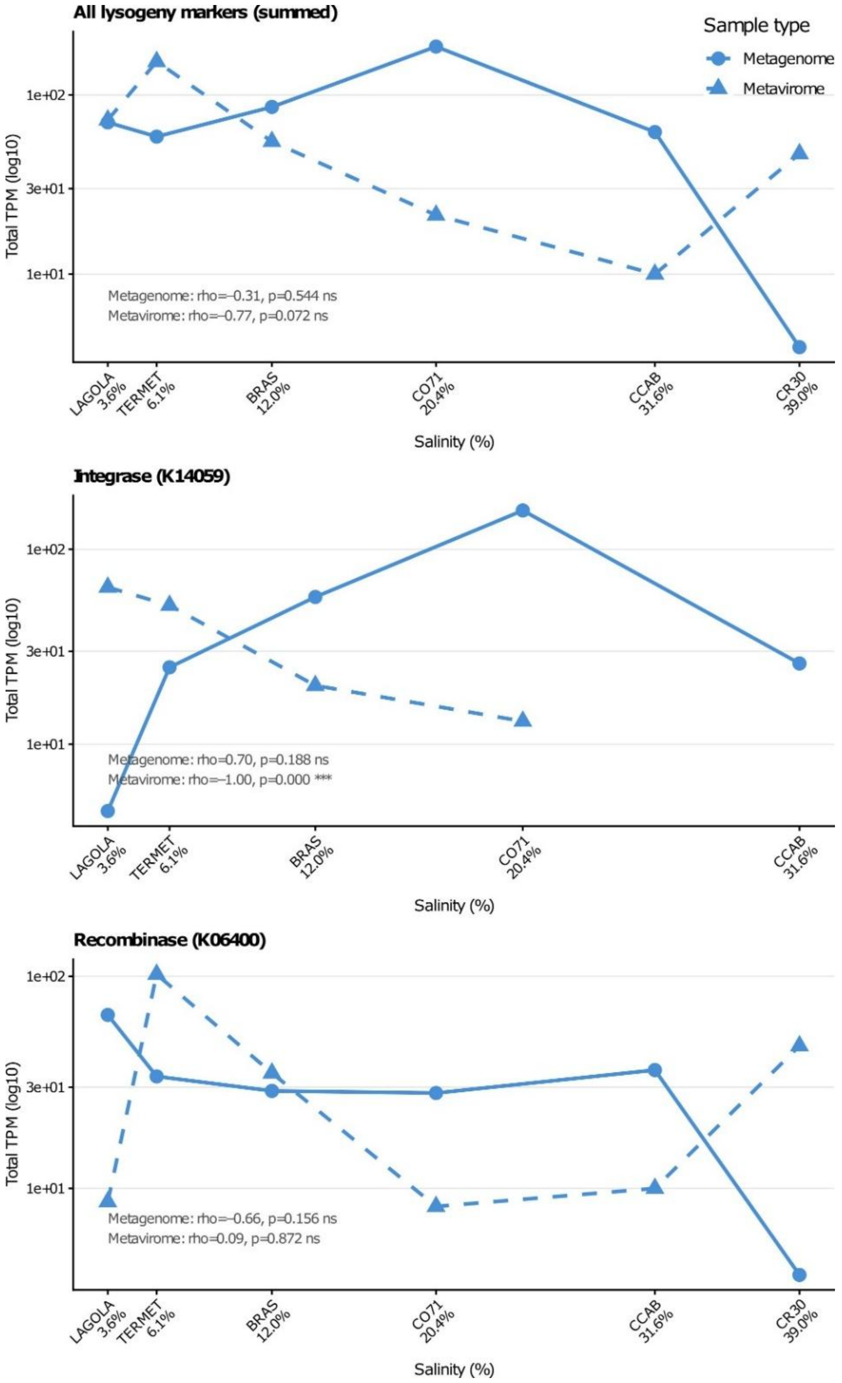


Figure 22. Abundance of lysogeny-associated marker genes across the salinity gradient at Bras del Port salterns. Total transcripts per million (TPM) of lysogeny marker genes detected in cellular metagenomes (solid line, circles) and viromes (dashed line, triangles) across six ponds spanning salinities from 3.6% (LAGOLA) to 39.0% (CR30). The y-axis is shown on the $\log_{10}$ scale. Three panels are shown: (top) all lysogeny markers combined; (middle) integrase (K14059); (bottom) recombinase/site-specific recombinase (K06400). Spearman's rank correlation coefficients (ρ) between log10-transformed TPM and salinity are indicated within each panel separately for metagenomes and viromes (significance levels: *$p < 0.05$, **$p < 0.01$, ***$p < 0.001$, ns: not significant). Lysogeny markers were defined as coding sequences annotated with the following KEGG Orthology identifiers: K14059 (tyrosine recombinase/integrase), K21039 (serine integrase), K21528 (serine recombinase), K06400 (site-specific recombinase, resolvase family), and K27464 (plasmid partition protein); of these, only K14059 and K06400 were detected in the dataset.

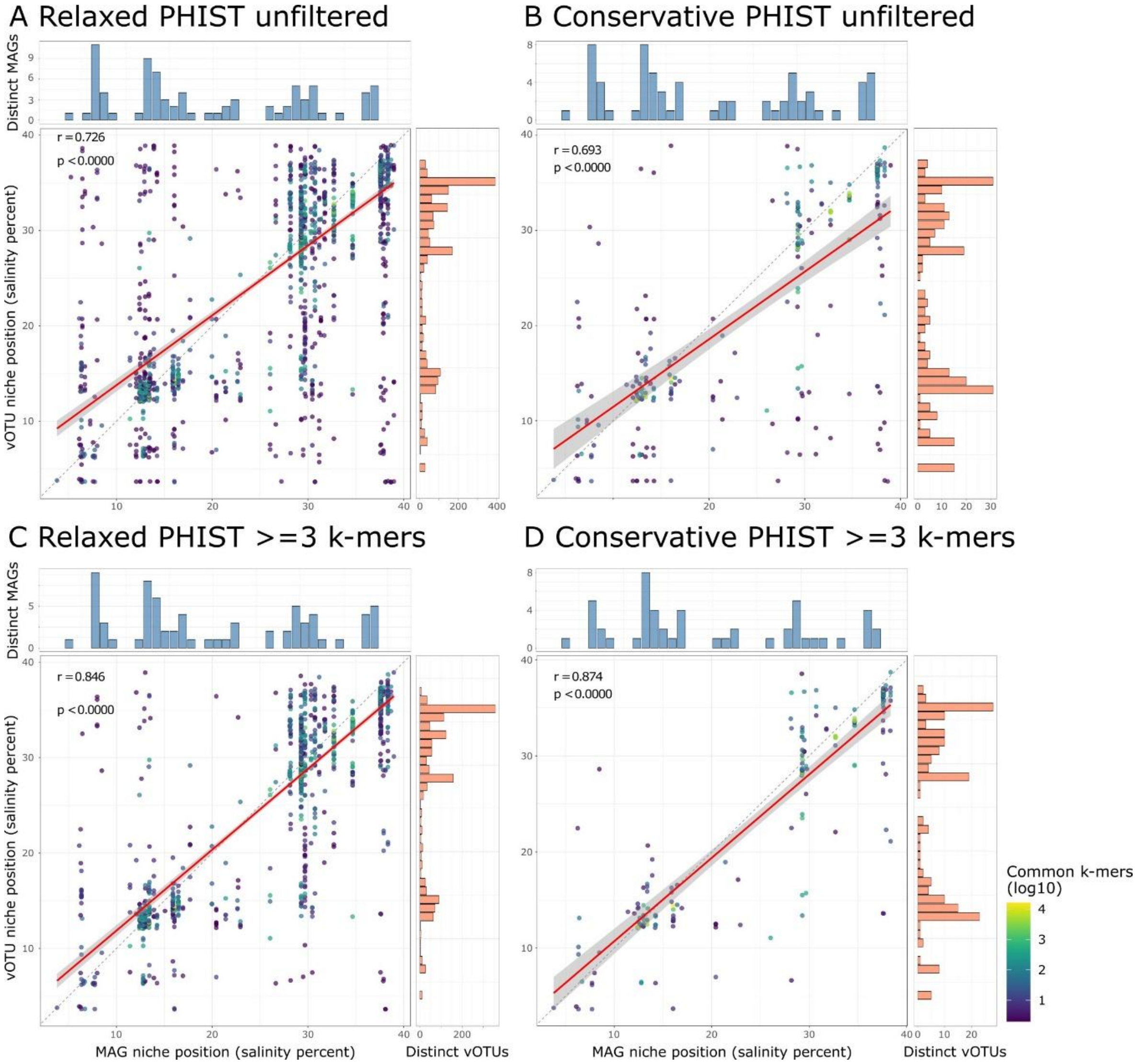


Figure S23. Niche position alignment of virus-host pairs across prediction stringency and vOTU dataset subsets. Scatter plots comparing the niche position of vOTUs versus their predicted host MAGs for each analytical subset, colored by the number of shared k-mers between the vOTU and MAG on a $\log_{10}$ scale. Marginal histograms show the number of distinct MAGs (top) and distinct vOTUs (right) across niche positions, allowing assessment of whether trends are driven by individual entities or distributed across multiple independent pairs. The dashed diagonal line represents the 1:1 relationship (perfect niche position alignment) and the solid red line shows the linear regression fit with 95% confidence interval. Pearson correlation coefficient r and associated p-value are shown in the upper left of each panel. (A) Relaxed vOTUs, unfiltered PHIST predictions (n = 1,875 pairs; r = 0.726, p < 0.0001). (B) Conservative vOTUs, unfiltered PHIST predictions (n = 267 pairs; r = 0.693, p < 0.0001). (C) Relaxed vOTUs, PHIST with ≥3 shared k-mers (n = 1,480 pairs; r = 0.846, p < 0.0001). (D) Conservative vOTUs, PHIST with ≥3 shared k-mers (n = 191 pairs; r = 0.874, p < 0.0001). Pearson correlations strengthen progressively from unfiltered to k-mer filtered subsets and from relaxed to conservative vOTU datasets. Pairs with higher k-mer counts (yellow-green) are concentrated near the 1:1 line across all subsets.

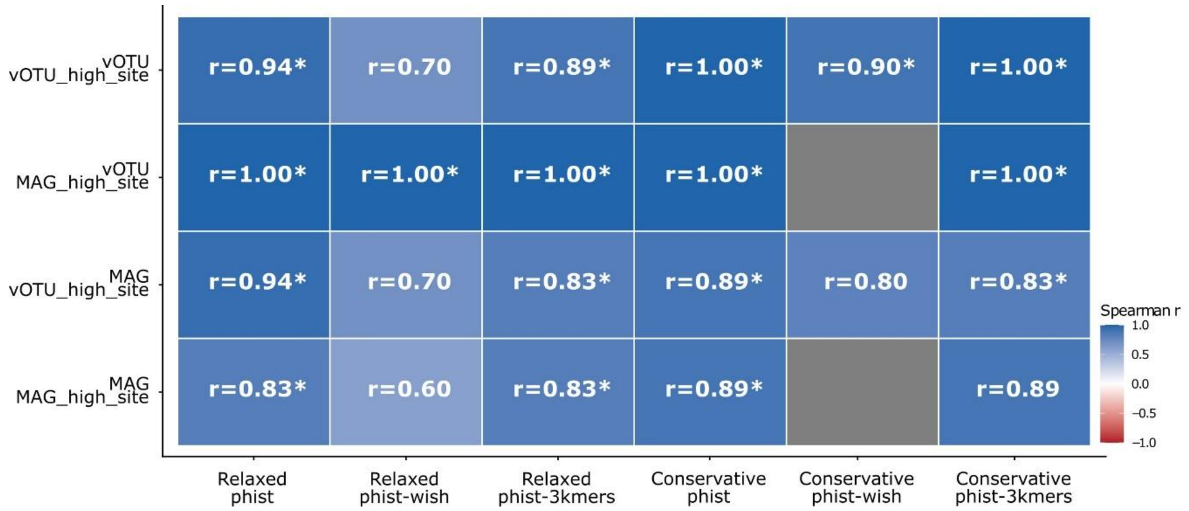


Figure S24. Spearman correlation between site-mean niche breadth and salinity for vOTUs and their predicted host MAGs across six analytical subsets. Each tile shows the Spearman rank correlation coefficient (r) between the mean niche breadth calculated per site and the corresponding salinity value, for vOTUs (virus) and MAGs (host) separately, and for two grouping strategies: pairs grouped by the site of highest vOTU abundance (vOTU_high_site) and pairs grouped by the site of highest MAG abundance (MAG_hig_hsite). Color intensity reflects the strength and direction of the correlation, with blue indicating a positive relationship (broader niches at higher salinities) and red indicating a negative relationship. Values marked with an asterisk (*) are statistically significant ($p < 0.05$). The six analytical subsets correspond to combinations of the relaxed and conservative vOTU datasets with three virus-host prediction filters: PHIST, PHIST-WiSH, and PHIST-3kmers. NA indicates subsets with fewer than five sites available for correlation, precluding statistical testing.

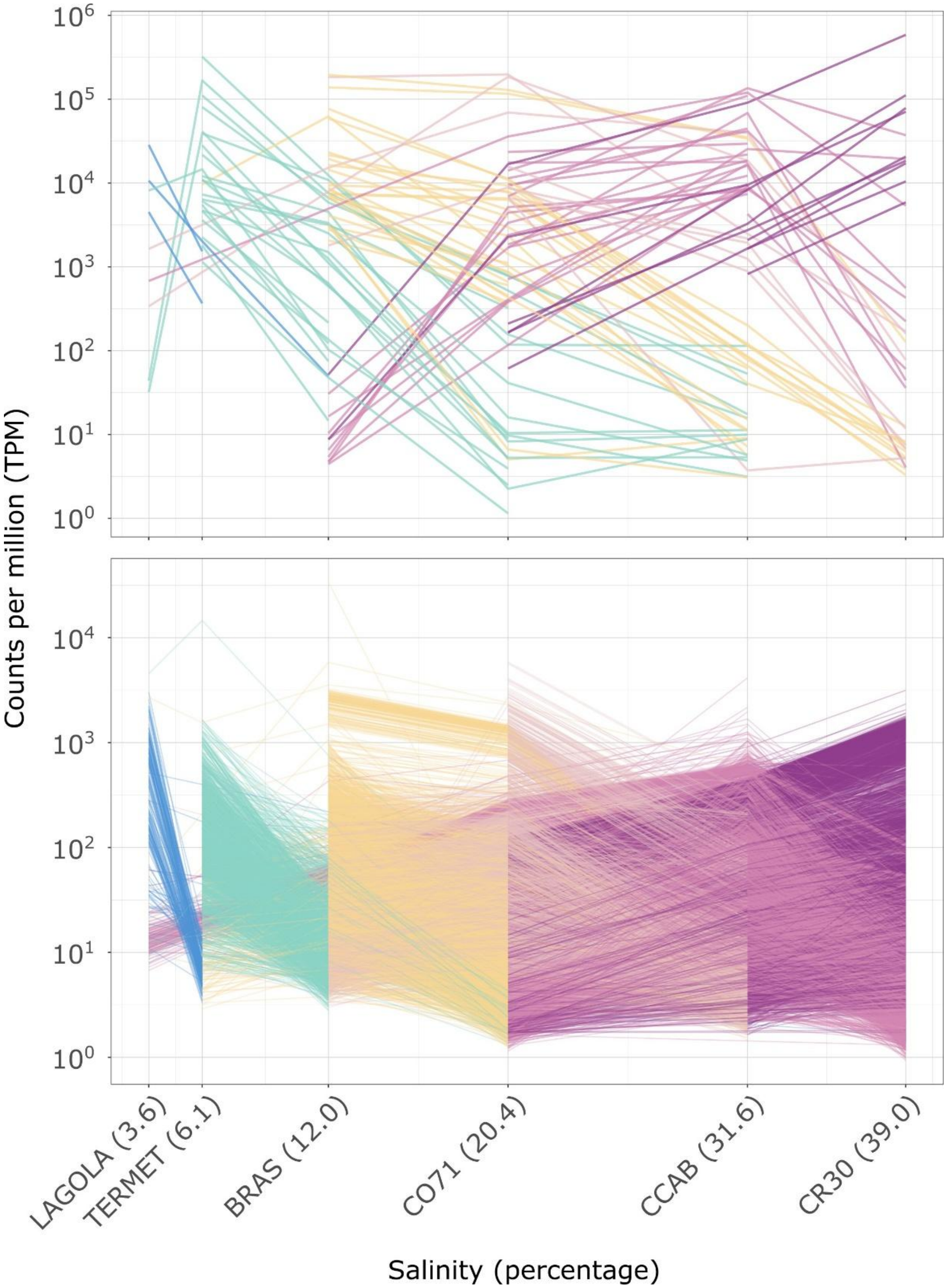


Figure S25. Distribution of MAGs and vOTUs across the salinity gradient. Line plots showing abundance trajectories of individual MAGs (upper panel, n=89) and vOTUs (lower panel, n=16,393) across six sampling sites along the salinity gradient. Each line represents a single entity colored by the sample where it achieved highest abundance (TPM). Y-axis is on a $\log_{10}$ scale. Only entities with at least two detected TPM values are included (MAGs detected with metapresence BER80_FUG50; vOTUs detected with ≥50% aligned fraction). Abundance values were quantified with coverM (≥95% identity, ≥50% read coverage). Patterns reveal site-specific prevalence and abundance dynamics of individual MAGs and vOTUs across the environmental gradient.

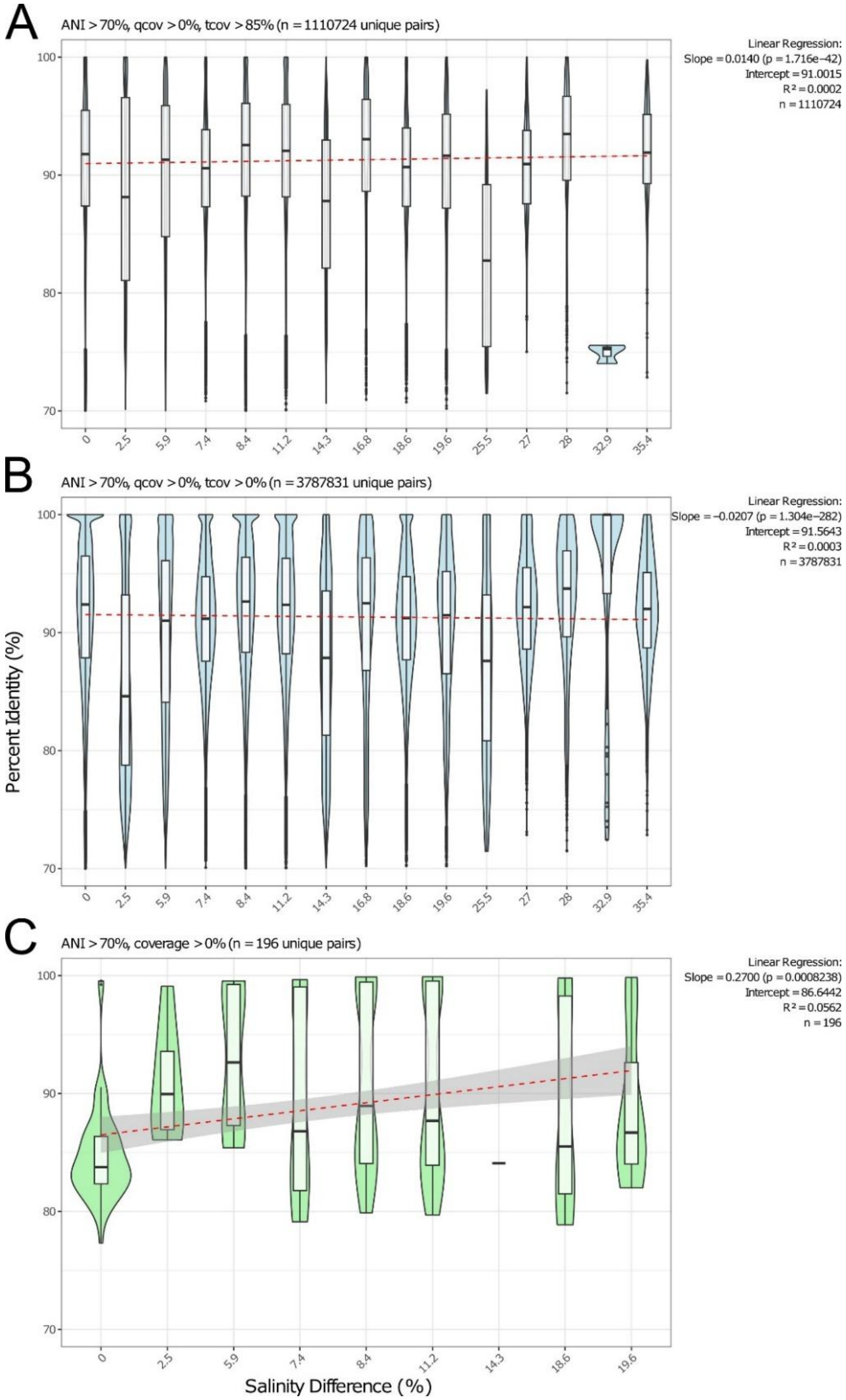


Figure S26. Relationship between genetic distance and salinity separation for MAGs and vOTUs. Violin, box, and whisker plots showing the relationship between average nucleotide identity (ANI) and absolute salinity difference for paired comparisons. A) vOTU pairs with >70% ANI, >0% query coverage, and >85% target coverage. B) vOTU pairs with >70% ANI, >0% query coverage, and >0% target coverage. C) MAG pairs with >70% ANI, >0% query coverage, and >0% target coverage. Numbers of paired comparisons are given in panel titles. Linear regression model statistics (slope, intercept, $R^2$, and p-value) are displayed for each dataset.

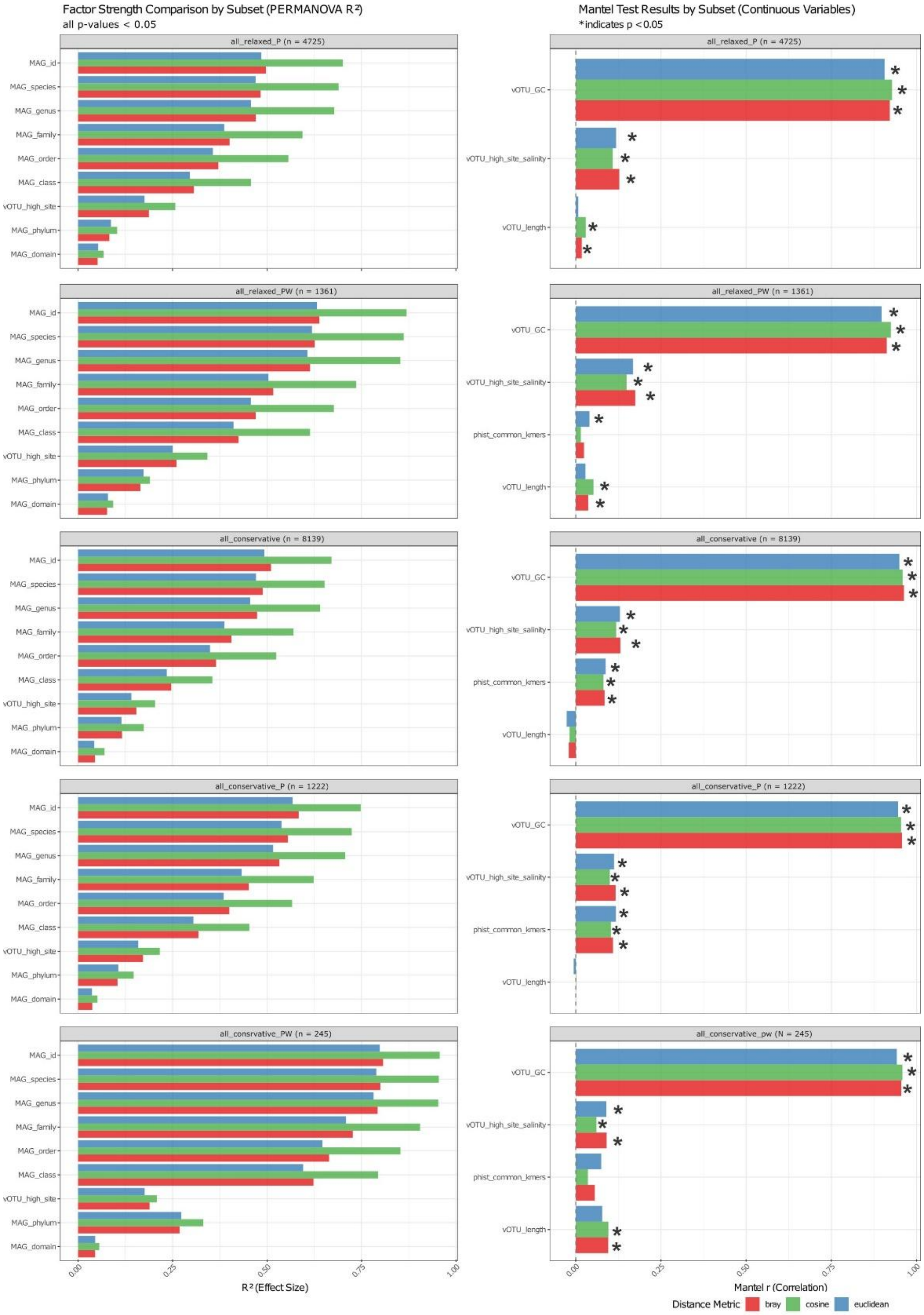


Figure S27. Effect sizes of variables on vOTU tetranucleotide frequency composition across prediction method subsets. Expansion of Figure 7B showing effect sizes for all variables using three distance metrics: Bray-Curtis (red), Cosine (green), and Euclidean (blue) distances, applied to different vOTU subsets. Left panels display effect sizes for categorical variables (host taxonomy and site with highest TPM) analyzed with PERMANOVA. Right panels display effect sizes for numerical variables (GC-content, salinity and contig length) analyzed with Mantel tests. Each row of panels represents a different vOTU subset defined by prediction method: conservative vOTUs with PHIST ≥3 shared k-mers, relaxed vOTUs with unfiltered PHIST, conservative vOTUs with unfiltered PHIST, relaxed vOTUs with PHIST-WiSH, and conservative vOTUs with PHIST-WiSH. Asterisks indicate significant associations (p<0.05).

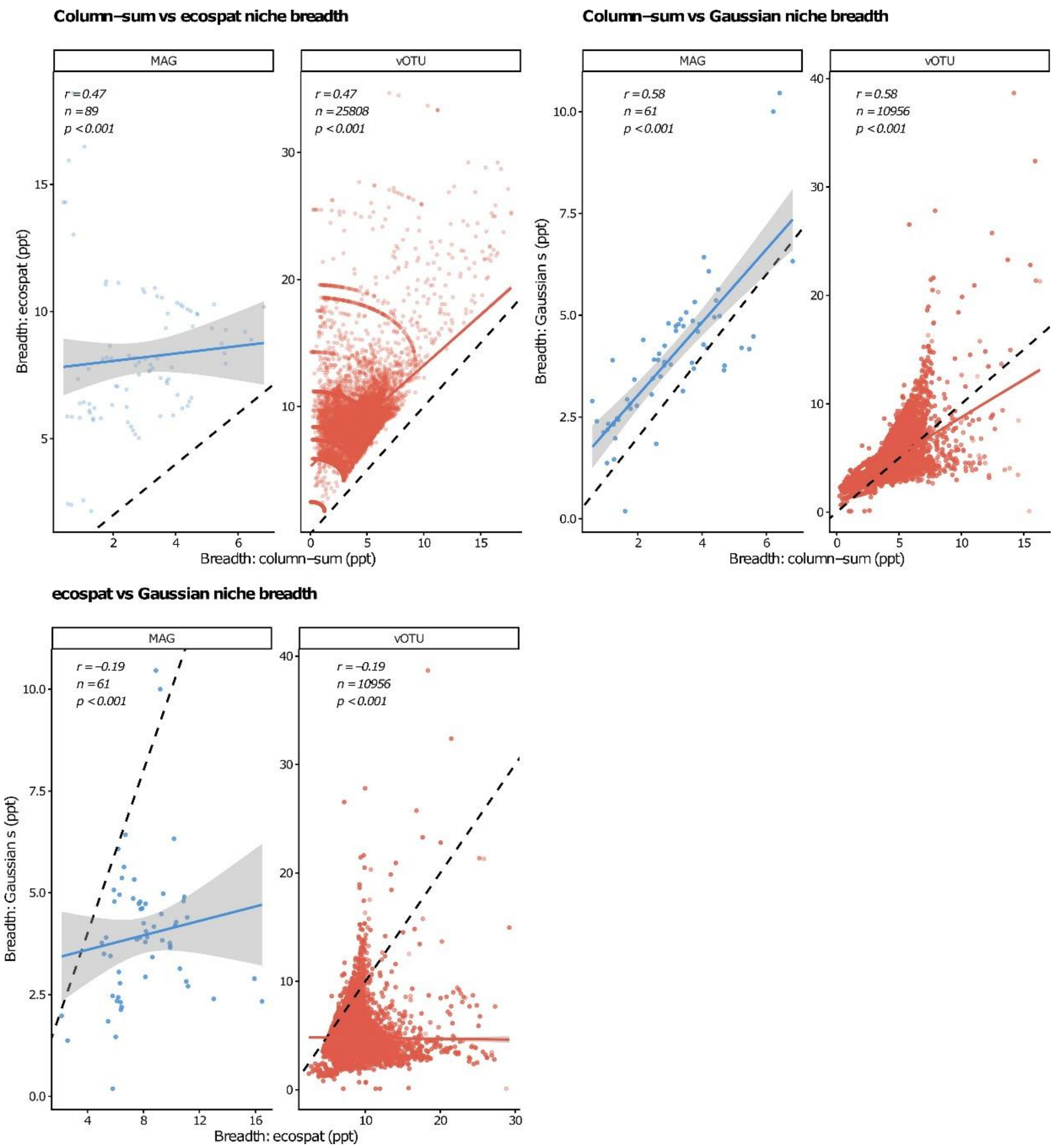


Figure S28. Comparison of niche breadth estimates between the column-sum normalisation method and two alternative approaches for MAGs and vOTUs. Upper left) Column-sum vs ecospat (ecospat::ecospat.nichePOSNB). Each point represents a single entity (MAG, blue; vOTU, red) detected in at least two sites. The x-axis shows niche breadth estimated by column-sum normalisation and the y-axis shows niche breadth estimated by ecospat. The dashed line indicates the 1:1 relationship. Ecospat breadths are systematically larger than column-sum breadths due to differences in weight normalisation and variance denominator (see Supplementary Text). The solid line shows the linear regression fit with 95% confidence interval, and Spearman r, n, and p-value are shown per panel. Upper right) Column-sum vs Gaussian σ. Niche breadth estimated by column-sum normalisation is compared against the σ parameter of a Gaussian response curve fitted to the TPM abundance of each entity across the six salinity values, using multi-start nonlinear least-squares fitting with three starting values for σ (range/8, range/4, range/2), retaining the fit with the highest $R^2$. Only entities where the Gaussian fit converged with $R^2 \geq 0.7$ are shown (MAGs: n = 61; vOTUs: n = 10,956). The solid line shows the linear regression fit with 95% confidence interval. Lower left) ecospat vs Gaussian σ. The same Gaussian-retained entities are used. The significant negative correlation (Spearman r = −0.19, $p < 0.001$ for both MAGs and vOTUs) indicates that ecospat and Gaussian breadths deviate from column-sum through independent mechanisms and in opposing directions, with entities assigned larger ecospat breadths tending to receive smaller Gaussian σ estimates and vice versa.

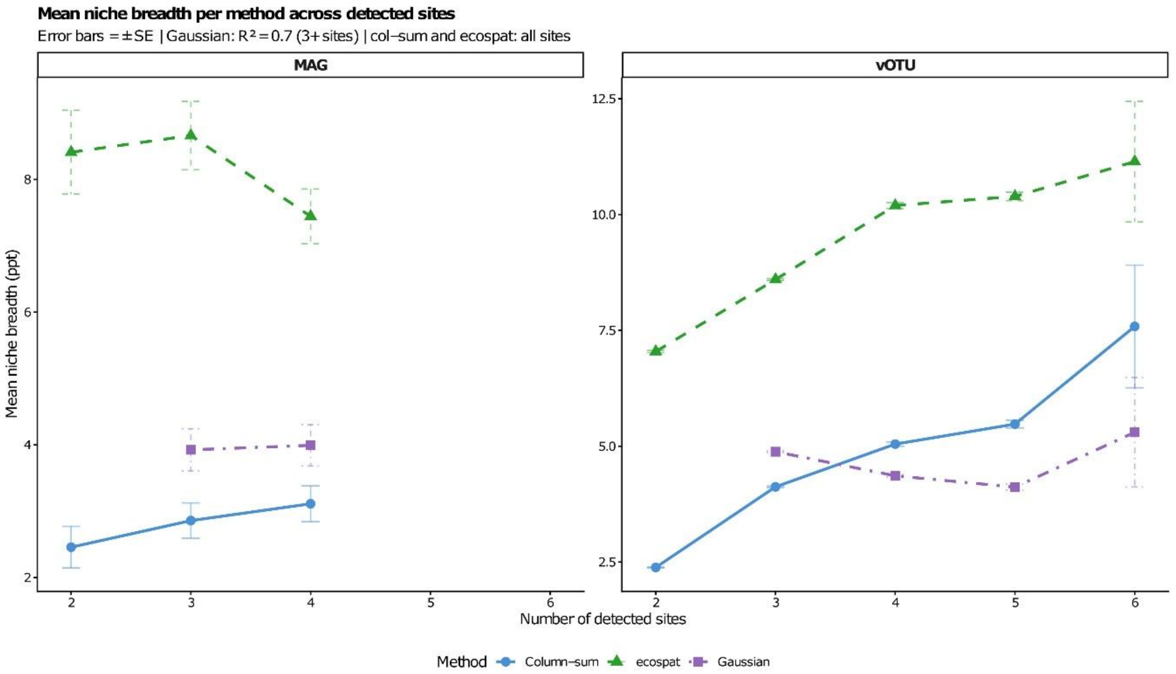


Figure S29. Mean niche breadth estimated by three methods across the number of detected sites for MAGs and vOTUs. Mean niche breadth (in ppt salinity) is shown for column-sum normalisation (blue solid line), ecospat (ecospat::ecospat.nichePOSNB; green dashed line), and Gaussian curve fitting using the σ parameter (purple dot-dash line). Column-sum and ecospat are shown for all entities detected at two or more sites; Gaussian fitting is shown only for entities detected at three or more sites where the fit converged with $R^2 \geq 0.7$, using multi-start nonlinear least-squares fitting. Error bars indicate ±1 standard error. Ecospat breadths are consistently and substantially larger than column-sum breadths across all numbers of detected sites, reflecting the systematic inflation introduced by max-normalised weights and the sum(w) - 1 variance denominator. Column-sum breadth increases steadily with the number of detected sites for both MAGs and vOTUs. Gaussian σ shows a non-monotonic relationship with the number of detected sites: for vOTUs it exceeds column-sum at three detected sites but falls below it at four and five sites before rising again at six, while for MAGs it remains stable between three and four detected sites at values above column-sum but well below ecospat. This non-monotonic pattern reflects the sensitivity of Gaussian σ to the shape and symmetry of the abundance distribution across the gradient, which varies with the number of constraining observations. Note that the MAG panel shows only two to four detected sites due to the limited number of MAGs with sufficient detections and retained Gaussian fits.

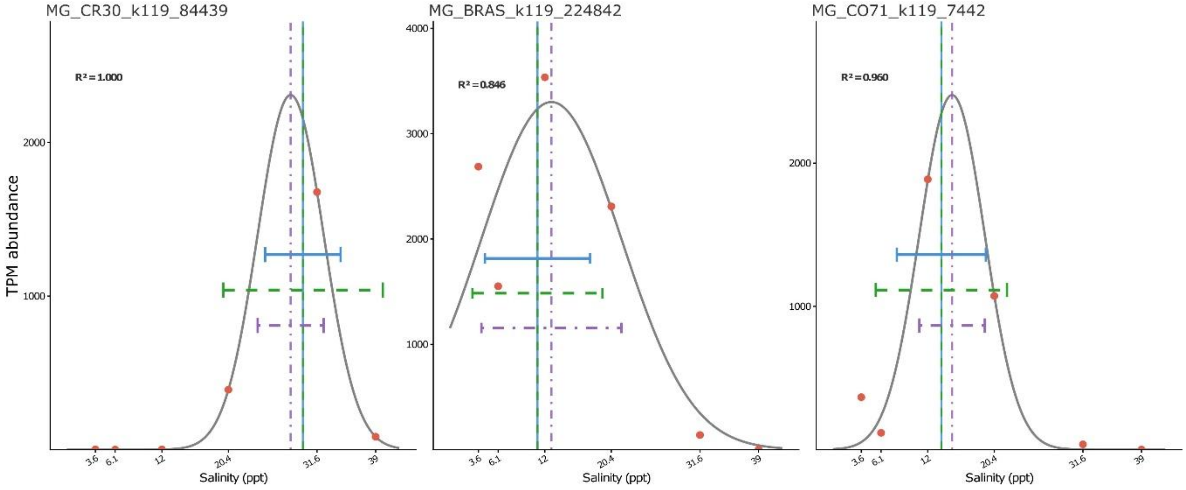


| Method | Position (ppt) | Breadth (ppt) |
|---|---|---|
| Column–sum | 29.82 | 4.77 |
| ecospat | 29.83 | 10.06 |
| Gaussian | 28.27 | 4.17 |

| Method | Position (ppt) | Breadth (ppt) |
|---|---|---|
| Column–sum | 11.06 | 6.65 |
| ecospat | 11.06 | 8.21 |
| Gaussian | 12.82 | 8.84 |

| Method | Position (ppt) | Breadth (ppt) |
|---|---|---|
| Column–sum | 13.72 | 5.62 |
| ecospat | 13.71 | 8.28 |
| Gaussian | 15.07 | 4.13 |



Figure S30. Gaussian niche fitting and comparison of three niche estimation methods for three example vOTUs detected at all six sampling sites. Each panel shows the TPM abundance of one vOTU across the salinity gradient (points) with the fitted Gaussian response curve (grey line). Vertical lines indicate the niche position estimated by each method: column-sum normalisation (blue solid), ecospat (green dashed), and Gaussian peak μ (purple dot-dash). Horizontal segments indicate the corresponding niche breadth estimate for each method, centred on the respective niche position, where breadth is defined as the abundance-weighted standard deviation for column-sum and ecospat, and as the Gaussian σ for the Gaussian method. The Gaussian curve was fitted using multi-start nonlinear least-squares optimisation, trying three starting values for σ (range/8, range/4, range/2) and retaining the fit with the highest $R^2$; the $R^2$ value is shown in the upper left of each panel. Numerical values of niche position and breadth for each method are shown in the table below each panel.

### Supplementary table captions

Table S1. Abundance of prokaryotic cells and viral-like particles across the salinity gradient. Quantification of prokaryotic cells and viral-like particles (VLPs) at each of the six sampling sites along the salinity gradient at Bras del Port salterns (Santa Pola, Spain). Cells/ml represents the concentration of prokaryotic cells enumerated by DAPI staining, with SD indicating standard deviation of replicate counts. VLPs/ml represents the concentration of viral-like particles enumerated by SYBR Gold staining, with SD indicating standard deviation of replicate counts. Ratio indicates the virus-to-cell ratio (VLPs/ml divided by Cells/ml).

Table S2. Relative abundances of CDSs by prokaryotic taxa. Relative abundance (as percentage of total mapped clean reads) of CDSs assigned to the ten most abundant prokaryotic taxa at 7 levels across the 12 samples. Taxonomic classification was performed using refineM against GTDB-R220 representative genomes. Read mapping was conducted with CoverM (≥95% identity, ≥50% read coverage). Remaining taxa below the top ten are grouped as "Other."

Table S3. Relative abundances of CDSs by functional category. Relative abundance (as percentage of total mapped clean reads) of CDSs assigned to the ten most abundant functional categories across the 12 samples. Functional annotations include: COG classifications (eggnog-mapper), KEGG and PFAM pathways (DRAM), and metabolic functions with individual HMMs and wider METABOLIC classifications. Values represent the proportion of reads from CDSs assigned to each functional classification at each site. Remaining functional categories below the top ten are grouped as "Other."

Table S4. Metagenome-assembled genome (MAG) characteristics and quality metrics. Detailed information for all 170 dereplicated MAGs recovered from metagenomic assemblies across the salinity gradient. NCBI genome and Biosample IDs are provided for MAGs > 90 % completeness. Taxonomic classification (gtdbtk_R220_classification) was assigned using GTDB-Tk v2.4.0 with database version R220. Quality assessment includes completeness, and contamination estimates from CheckM v1.0.12 and CheckM2 v1.0.2. Assembly metrics include genome size, number and length statistics of contigs (N50, mean length, longest contig), GC content, and coding density. Closest reference genome information (gtdbtk_closest_genome_reference, ANI, alignment fraction) links MAGs to cultured representatives from intermediate results from gtdb-tk. Gene content includes predicted CDSs, pseudogenes, hypothetical proteins, signal peptides, and small CDSs (sCDSs) annotated with bakta v1.9.4. Genomic features include tRNA and rRNA counts, CRISPR arrays, ncRNAs, and replication-associated features (oriCs, oriVs, oriTs). GSC_classification indicates genome quality status (high or medium) according to Genomic Standards Consortium criteria. Distribution across samples was determined by read mapping with CoverM v0.6.1 (covered_bases, mean, RPKM, TPM, coverage metrics; ≥95% identity, ≥50% read alignment). Presence-absence patterns were additionally evaluated using metapresence v1.0 with default parameters.

Table S5. Viral operational taxonomic unit (vOTU) characteristics, quality metrics, and host associations. Detailed information for all ~55,000 vOTUs recovered from viromic and metagenomic assemblies across the salinity gradient. Sequence properties include length (bp), GC content (%), and sample_origin indicating the primary source (metagenome or virome). Conservative classification (binary) indicates if the vOTU was from the relaxed or conservative set according to the MVP software. Viral and plasmid classifications were assigned using geNomad, including genome type (virus, plasmid, provirus), topology (circular/linear), gene count, hallmark proteins, taxonomic assignment, and quality scores. Plasmid metrics additionally include conjugation and antibiotic resistance genes. CheckV quality assessment includes completeness estimate, contamination level, quality classification (complete to low-quality), and potential quality warnings. Abundance across samples is quantified as TPM (transcripts per million) in each metagenome (MG_) and metavirome (MV_) sample. Columns maxTPM_site, maxTPM_site_MG, and maxTPM_site_MV indicate the site and sample type where maximum abundance was detected. Presence-absence patterns (presabs_*) indicate vOTU detection at each site based on AF10 threshold analysis (default parameters). Host MAG associations were predicted using PHIST (Host_MAG_id) with MAG_all_gtdbtk_R220_classification from the gtdb-tk data from the respective MAG. Reference genome names and strains were searched for each described genera through a search in the GTDB site (https://gtdb.ecogenomic.org/).

Table S6. Virus-host pair characteristics, niche metrics, and ecological strategies. Detailed information for virus-host pairs (vOTU-MAG) with sufficient detection across samples to calculate niche position and breadth (minimum two abundance values with >50 % of aligned fraction across the six sampling sites each). PHIST prediction parameters include: common_kmers (number of shared k-mers between vOTU and MAG), phist_pvalue (raw p-value), and phist_adj_pvalue (adjusted p-value). Filtering columns indicate which vOTU subsets each pair belongs to: conservative (vOTU detection ≥50% aligned fraction), phist_3kmers (PHIST ≥3 shared k-mers), and phist_wish (PHIST-WiSH predictions). MAG taxonomy includes domain through species level classification (GTDB-R220). Niche metrics calculated using our col-sum method include: MAG_NichePosition and vOTU_NichePosition (salinity optimum for each entity), and MAG_NicheBreadth and vOTU_NicheBreadth (salinity range surrounding niche position). Ecological strategies (MAG_Strategy, vOTU_Strategy) classify each entity as: Generalist (top 5% niche breadth), Intermediate (middle 90%), or Specialist (bottom 5% niche breadth).

Table S7. Experimentally confirmed viral reference genomes and associated vOTUs. Catalog of viral reference genomes retrieved from literature that infect prokaryotic strains associated with representative genomes of MAG species in the dataset. NCBI_id indicates the GenBank accession number for the reference viral genome. Host_strain_species identifies the prokaryotic host species infected by each virus. Reference provides the literature citation, with DOI linking to the original publication. Number_of_associated_vOTUs quantifies relaxed vOTUs from the dataset that share sequence similarity with each reference genome, identified through BLASTn searches with a relaxed threshold (≥90% identity, ≥85% query coverage).

Table S8. Site-specific paired comparisons of niche breadth between vOTUs and their predicted host MAGs across six analytical subsets calculated with the col-sum strategy. For each dataset, site, and grouping strategy, niche breadth values (expressed as the standard deviation of the salinity-weighted abundance distribution, in %) were compared between vOTUs and their cognate MAGs using paired statistical tests. The grouping strategy indicates whether pairs were assigned to sites based on the site of highest vOTU abundance (vOTU_high_site) or the site of highest MAG abundance (MAG_high_site). MAG_Mean and vOTU_Mean indicate the mean niche breadth per entity at that site; MAG_SD and vOTU_SD indicate the corresponding standard deviations. Mean_Diff is calculated as vOTU_Mean minus MAG_Mean, with positive values indicating broader vOTU niches. The Test_Used column indicates whether a paired Wilcoxon signed-rank test or a paired t-test was applied, selected based on the result of a Shapiro-Wilk normality test on the pairwise differences. The Statistic column reports the corresponding test statistic (W for Wilcoxon, t for paired t-test). Significant indicates whether the p-value fell below the 0.05 threshold. The final column (vOTU > MAG) indicates the direction of the significant difference where applicable. Rows marked as "Insufficient data" correspond to sites with fewer than three pairs available for testing. The six analytical subsets correspond to combinations of the relaxed and conservative vOTU datasets with three virus-host prediction filters: unfiltered PHIST, PHIST-WiSH, and PHIST-3kmers.
